# Coupling high-throughput protease enzymology with viral replication reveals biochemical constraints of viral fitness

**DOI:** 10.64898/2026.01.27.702130

**Authors:** Dylan Aidlen, William Vinh Thanh Vo, Nicholas J. Young, Julia Rosecrans, Anna Kurianowicz, Shih-Wei Chuo, Garrison P. R. Asper, Dashiell Anderson, Jacob A. Posner, Duncan F. Muir, Nicholas Freitas, Melanie Ott, Charles S. Craik, Taha Y. Taha, Margaux M. Pinney

## Abstract

Proteases govern essential biological processes and are key drug targets, yet how protease sequence variation quantitatively reshapes biochemical parameters and constrains biological fitness remains poorly understood. Here, we integrate high-throughput *in vitro* enzymology with cellular assays to link protease sequence, biochemistry, and fitness. We extend a microfluidic platform for high-throughput protease enzymology (HT-MEK^pro^), which is broadly applicable across protease families and catalytic classes, enabling measurement of catalytic turnover (*k*_cat_), Michaelis constant (*K*_M_), inhibitor potency (IC_50_), and relative substrate specificity for 10^2^-10^3^ variants. Applied to the SARS-CoV-2 main protease (M^pro^), HT-MEK^pro^ generated parallel catalytic and inhibitory landscapes for >400 variants. Integration with viral replication and in-cell cleavage assays reveals that variants with altered substrate specificity fail to support replication, suggesting imbalanced polyprotein processing as a constraint on viral fitness. More broadly, these data can enable mechanistically grounded modeling of protease sequence–property relationships and inform strategies for pharmacological modulation beyond active-site inhibition.

## Main Text

Proteases, enzymes that hydrolyze peptide bonds in proteins, are central to diverse biological processes, including cellular signaling, protein homeostasis, and viral replication, and account for ∼2% of the human genome.^1,2^ For decades, quantitative enzymology of proteases has shaped our molecular understanding of how enzymes work, and proteases remain the classic example used to describe enzyme mechanisms.^3,4^ Their roles in physiology and disease have made proteases prominent therapeutic targets, with inhibitors achieving clinical success against viral infections, diabetes, and cardiovascular diseases, including hypertension and blood coagulation.^5,6^ At the same time, proteases’ specificity and catalytic efficiency position them as promising therapeutics and versatile biotechnological tools.^7,8^ Yet a fundamental challenge remains: how do changes in protease sequence quantitatively reshape catalysis, substrate specificity, and inhibitor sensitivity, and how do these biochemical effects ultimately constrain the fitness of the cell or virus they operate in?

Despite decades of biochemical study, connecting protease sequence to quantitative function has remained a challenge of scale and context. Classical enzymology provides gold-standard kinetic and thermodynamic constants. Still, it is inherently low-throughput, typically limited to fewer than ten variants, because each must be individually expressed, purified, and assayed—although a few exceptional studies have extended this to several dozen or nearly one hundred variants.^9,10^ Cell-based selection and continuous evolution methods can rapidly identify variants with altered activity, specificity, or resistance. ^11–15^ However, these methods indirectly link sequence to biochemistry and can conflate catalytic effects with expression and folding. Substrate-profiling approaches, such as positional scanning libraries^16–20^, mass-spectrometry-based methods^21–25^, and label-free degradomics^26^, have mapped the specificity of wild-type (WT) proteases in remarkable detail; however, they have limited throughput to map the quantitative, variant-resolved landscapes necessary to dissect how protease specificity evolves or is redesigned molecularly. As a result, we still lack systematic, biochemically grounded maps that connect protease sequence to catalysis, specificity, and inhibition—and that directly link these biochemical properties to cellular or viral outcomes.

Here, we bridge this gap by integrating mechanistic, high-throughput *in vitro* enzymology with viral replication assays. We establish a microfluidic platform for quantitative protease enzymology (HT-MEK^pro^) that measures catalytic and inhibitory parameters across hundreds to thousands of variants, thereby expanding the scale of quantitative kinetic data by orders of magnitude beyond previous enzymology studies. We apply HT-MEK^pro^ to the SARS-CoV-2 M^pro^, defining its catalytic, specificity, and resistance landscapes and directly linking them to viral replication. Together, these results reveal how biochemical properties—particularly the specificity for polyprotein cleavage sites—constrain viral fitness and illustrate a general strategy for linking enzyme biochemistry to biological outcomes.

### Extension of HT-MEK to high-throughput protease enzymology (HT-MEK^pro^)

Here, we extend the High-Throughput Mechanistic Enzyme Kinetics (HT-MEK) microfluidic platform, previously developed with phosphatases and adapted to kinases,^27–29^ to proteases, representing the first application of this platform to protease enzymology. The quantitative analysis of catalysis and inhibition of clinically and evolutionarily relevant protease variants is enabled at scale by HT-MEK^pro^. We retained the underlying two-layer PDMS device while redesigning the on-chip biochemical workflow to support high-throughput protease enzymology. The device consists of ∼1800 isolated reaction chambers, and with an updated workflow, allows parallel *in vitro* expression, purification, activation, and kinetic characterization, with and without inhibitors, of hundreds to thousands of protease variants (Fig. 1a, Figs. S1 and S2, see Methods). This extended platform, termed HT-MEK^pro^, permits direct measurement of catalytic turnover (*k*_cat_), Michaelis constant (*K*_M_), and inhibitor potency (IC_50_) across protease variant libraries under uniform biochemical conditions.

**Figure 1.**
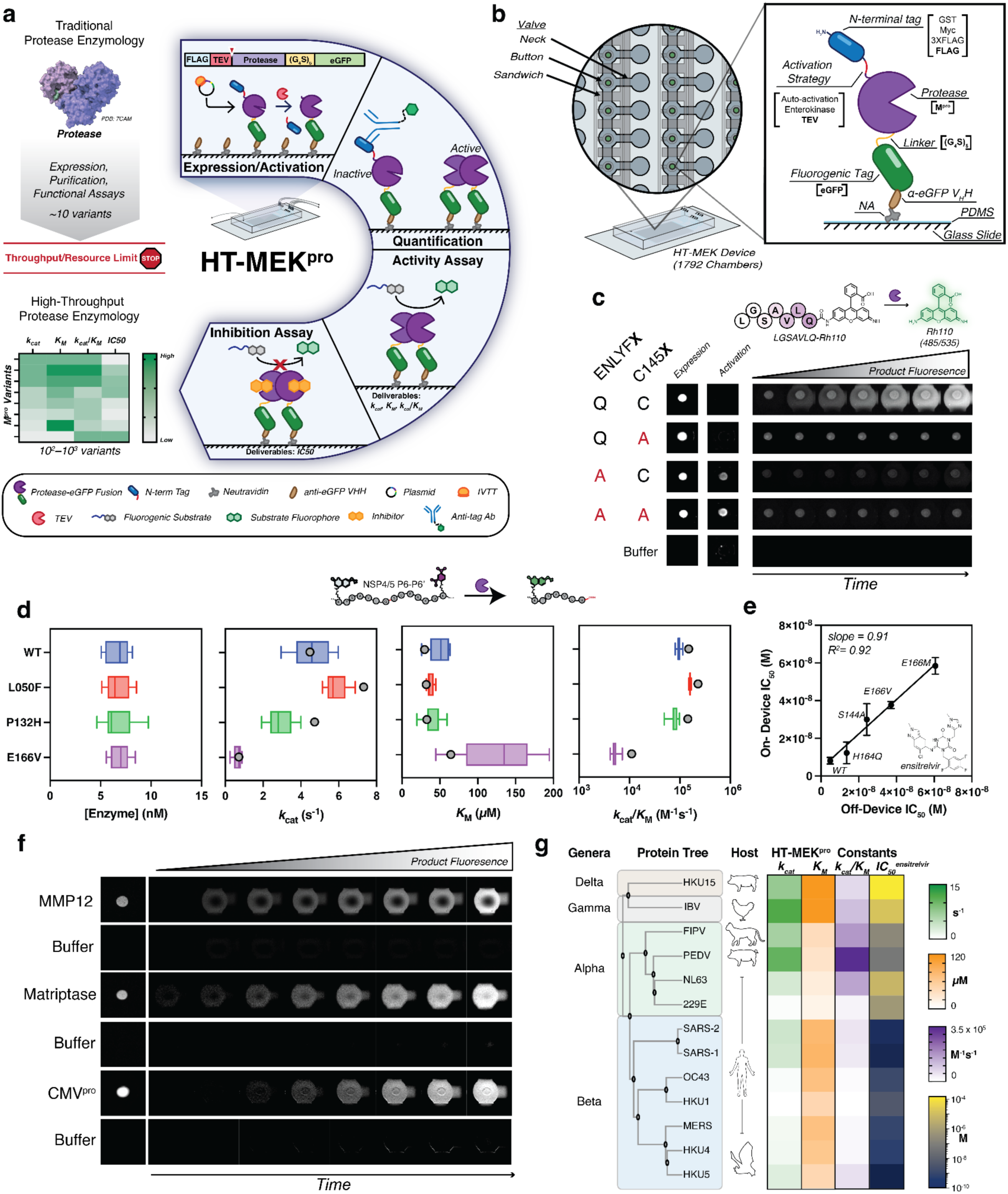
Adaptation and generality of HT-MEK for high-throughput protease enzymology. **(a)** Schematic of HT-MEK^pro,^ a microfluidic platform for parallel quantitative protease enzymology. Up to 1,792 protease variants are expressed *in vitro* as protease–eGFP fusions, captured via anti-eGFP nanobodies, and activated by TEV protease to generate native N-termini. Catalysis is quantified using compatible, internally quenched peptide substrates or dye release substrates to obtain Michaelis–Menten parameters (*k*_cat_, *K*_M_). Inhibitor dose–responses yield IC_50_ values across entire variant libraries, enabling quantitative maps of activity, specificity, and resistance. **(b)** Device layout and chamber workflow, including pull-down scheme and M^pro^ fusion design. **(c)** Representative on-device fluorescence microscopy images showing immobilized enzyme (left), anti-FLAG signal reporting activation (center), and product accumulation over time from an Rh110 substrate (right). Shown for WT, catalytically inactive C145A, and TEV-site controls (ENLYFQ/A). **(d)** Concordance between on-device Michaelis–Menten measurements and off-device (plate-based) kinetics. Bars, on-device (n = 7–9); grey points, off-device (n = 3). **(e)** Agreement of ensitrelvir IC50 values measured on-device (n = 7–9) versus off-device (n = 3), with linear fit shown. **(f)** Representative on-device activity assays for three additional, mechanistically distinct proteases: a viral serine protease (CMV protease), a human serine protease (matriptase), and a human metalloprotease (MMP12). **(g)** On-device *k*_cat_, *K*_M_, *k*_cat_/*K*_M_, and IC_50_ measurements with ensitrelvir for SARS-CoV-2 WT M^pro^ and 12 coronavirus M^pro^ orthologs spanning four genera and multiple hosts, assayed with an internally quenched peptide substrate derived from the SARS-CoV-2 NSP4-5 junction. See Table S3 for values.

As a test case, we focused on the SARS-CoV-2 M^pro^, also known as 3CL^pro^, a cysteine protease encoded by *nsp5* of Orf1a that is essential for coronavirus replication through its role in cleaving viral polyproteins at conserved Leu-Gln↓Ser motifs to release non-structural proteins 4–16 (nsp4–16).^30,31^ M^pro^ is an established antiviral target, exemplified by clinically approved inhibitors such as nirmatrelvir and ensitrelvir.^32–34^ M^pro^ is expressed as part of the viral polyproteins pp1a and pp1ab (nsp5) and undergoes self-cleavage at the N- and C-termini, releasing active M^pro^, and generating Ser1 at the N-terminus, which interacts with the active site of the adjacent protomer, stabilizing the S1 pocket.^35,36^ Thus, M^pro^ requires precise N-terminal processing to adopt its mature, catalytically competent conformation.

We first established a robust on-device expression and activation workflow for M^pro^. The M^pro^ variants were expressed as fusions, with an N-terminal FLAG tag and a C-terminal eGFP separated by a flexible Ser-Gly (G_4_S)_5_ linker. The N-terminal FLAG-tag supported higher expression relative to other tags tested (Fig. S3). Following expression, chamber-specific purifications were performed via an anti-GFP nanobody pull-down, and M^pro^ concentrations were quantified within each chamber via meGFP fluorescence (Fig. 1b). Variants were subsequently activated by cleavage of the N-terminal FLAG tag using TEV protease, which, relative to other activation strategies tested, yielded consistent activation across variant libraries and avoided off-target cleavage observed with alternative approaches (Fig. 1a–c, Figs. S4–S6). Together, the minimal N-terminal FLAG tag coupled to the TEV site supported high on-device expression, efficient activation, reproducible catalytic activity, and inhibition by nirmatrelvir (Fig. 1c, Fig. S8, Table S1).

We next optimized substrate design and benchmarked on-device kinetic measurements. Internally quenched fluorogenic peptide substrates commonly used in plate-based M^pro^ assays showed poor solubility on-device (Fig. S9). To ensure robust on-device kinetic measurements, we developed internally quenched fluorogenic peptide substrates based on native SARS-CoV-2 polyprotein junctions, with improved solubility and optical compatibility with microfluidic assays (Fig. 1c, Fig. S9). Using a P6-P6’ NSP4–5–derived substrate with a N-terminal 5-carboxyfluorescein (5-FAM) dye and C-terminal dinitrophenolate (DNP) quencher group, HT-MEK^pro^ measurements of *k*_cat_ and *K*_M_ for reference variants—including WT and clinically relevant mutants—aligned with off-device assays and reproduced known trends in catalytic efficiency (Fig. 1d).

Finally, we benchmarked HT-MEK^pro^ for high-throughput, quantitative inhibitor profiling. Unlike traditional approaches that screen tens to hundreds of inhibitors against a limited number of enzyme variants, HT-MEK^pro^ enables systematic evaluation of inhibition across extensive protease variant libraries and a focused set of key inhibitors. IC_50_ values for ensitrelvir,^32–34^ measured on device for WT and established resistance mutants agreed closely with off-device measurements, demonstrating the ability of HT-MEK^pro^ to dissect inhibitor resistance at scale (R² = 0.92; Fig. 1e, Fig. S10). Across kinetic and inhibition measurements at the library scale (see below), coefficients of variation were typically <0.2, indicating strong reproducibility (Fig. S11).

### HT-MEK^pro^ is broadly generalizable across protease families and catalytic classes

Proteases are a highly diverse class of enzymes, spanning distinct catalytic mechanisms and evolutionary origins.^1,2^ To test the breadth of HT-MEK^pro^, we applied it to proteases from diverse evolutionary and functional contexts. We first assayed three proteases that are structurally and mechanistically distinct from coronavirus M^pro^: the cytomegalovirus (CMV) serine protease,^37^ the human serine protease matriptase,^38^ and the human metalloprotease MMP-12^39^ (Fig. 1f). Together with M^pro^, these enzymes span the three major catalytic classes: cysteine, serine, and metalloproteases. For each target, optimized construct designs and assay conditions supported robust *in vitro* expression and catalysis, demonstrating that HT-MEK^pro^ is adaptable across diverse catalytic chemistries and structural frameworks.

We then tested a panel of 12 coronavirus M^pro^ orthologs, spanning all major genera (alpha, beta, gamma, delta) and hosts (human, porcine, avian, murine, feline, and bat), with 43–95% sequence identity to SARS-CoV-2 M^pro^ (Fig. 1g, Table S2). All twelve orthologs exhibited expression comparable to that of SARS-CoV-2 and measurable activity (Fig. 1g). Across orthologs, catalytic efficiencies (*k*_cat_/*K*_M_) on the SARS-CoV-2-derived NSP4-5 substrate varied over two orders of magnitude, despite highly conserved active sites, with several exceeding SARS-CoV-2 M^pro^, with differences being driven by changes in *k*_cat_, *K*_M_, or both (Fig. 1g, Table S3, Fig. S12).

Finally, we examined the susceptibility of these orthologs to the clinically approved noncovalent inhibitor ensitrelvir.^32–34^ IC_50_ values spanned five orders of magnitude, with SARS-CoV, and the two bat coronaviruses, HKU4, and HKU5 being potently inhibited (Fig. 1g). In contrast, others, such as NL63 and 229E, showed less potent inhibition, consistent with clinical reports of natural resistance (Fig. 1g). These data highlight the utility of HT-MEK^pro^ for identifying inhibitors with broad-spectrum inhibitory potential and for dissecting natural sequence variation that confers resistance.

### Mapping the catalytic and resistance landscapes of SARS-CoV-2 M^pro^

To quantify how clinical and evolutionary sequence variation reshapes M^pro^ catalysis and inhibitor sensitivity, we generated a library of 397 SARS-CoV-2 M^pro^ variants spanning functional, clinical, and evolutionary sequence space (Fig. 2a; Figs. S13–S16). The first half of this library includes mutations from variants of concern and high-frequency clinical isolates, targeted substitutions at residues involved in substrate and inhibitor recognition, and hyperactivating mutations identified in a yeast-based selection screen (see Methods).^15^ To place SARS-CoV-2 sequence M^pro^ variation in an evolutionary context, the second half of the library was composed of substitutions to amino acids found in the human alphacoronavirus NL63, whose M^pro^ is naturally resistant to ensitrelvir and exhibits elevated activity on the SARS-CoV-2 NSP4-5 substrate (Fig. 1g). Together, this library enables quantitative mapping of catalytic and inhibition landscapes at a scale far exceeding prior protease enzymology studies.

**Figure 2.**
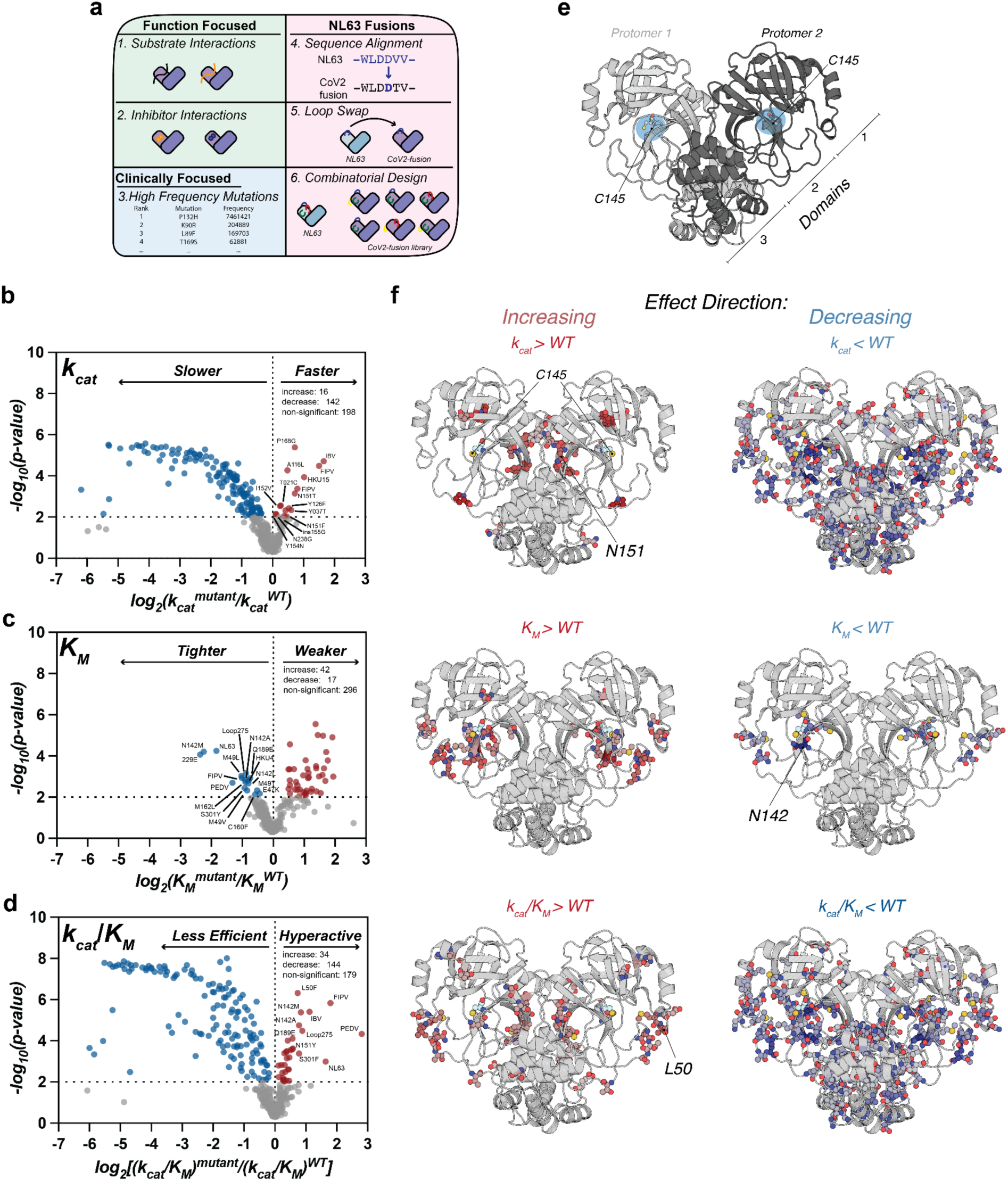
Catalytic landscape of SARS-CoV-2 M^pro^ revealed by HT-MEK^pro^. **(a)** Variant library design. Schematic of 397 M^pro^ variants spanning clinical/VOC sites, targeted substitutions at substrate/inhibitor-contacting residues, yeast-screen hits enriched for hyperactivity, and NL63-inspired evolutionary swaps (single residues, loops, and combinatorial designs). **(b–d)** Volcano plots showing variant effects relative to WT SARS-CoV-2 on **(b)** *k*_cat_, **(c)** *K*_M_, and **(d)** *k*_cat_/*K*_M_; function-enhancing mutations are annotated (complete values in Tables S4–S5). Orthologs are included for comparison (complete values in Table S3). **(e)** Structural architecture of SARS-CoV-2 M^pro^ (PDB 7CAM) with domains and active site indicated (blue) **(f)** Structural mapping of variant effects onto SARS-CoV-2 M^pro^ (PDB 7CAM); for simplicity, only significantly altered variants (blue and red points from b–d) are shown, and only SARS-CoV-2 point mutations (no deletions or loop swaps). Residues are colored according to the magnitude of the largest significant effect observed at that position. Residues highlighted in the main text are labeled.

Using HT-MEK^pro^, we obtained Michaelis–Menten parameters (*k*_cat_, *K*_M_, and *k*_cat_/*K*_M_) for 332 variants with catalytic activity above background, with an average of 6-7 biological replicates per variant, and strong reproducibility across independent experiments (R^2^ = 0.87-0.97; Fig. S17; Tables S4 and S5). Most substitutions were neutral or deleterious relative to WT; however, many increased *k*_cat_/*K*_M_, which arose through diverse kinetic mechanisms, including increased *k*_cat_, lowered *K*_M_, or contributions from both (Fig. 2b–d; Tables S4–S5). These data underscore the ability of HT-MEK^pro^ to resolve distinct kinetic routes to similar net catalytic efficiencies, which may have different functional implications in cellular contexts despite comparable overall activity. For example, we observed that the N142M mutation in the substrate binding pocket decreases *K*_M_ by ∼5-fold, possibly due to altered interactions with the P3 Val residue in NSP4-5, a finding supported by molecular dynamics (MD) simulations (Fig. S18).

Catalytic enhancement was not limited to residues directly interacting with the substrate. We also observed increased *k*_cat_ and decreased *K*_M_ from mutations located more than 15–20 Å from the catalytic cysteine, consistent with long-range allosteric coupling within M^pro^ (Fig. 2e,f). In one case, cooperative effects were evident: the swap of an entire loop from NL63 for the corresponding residues in SARS-CoV-2 M^pro^ (residues 275–292, “Loop 275”) produced a modest but significant 2-fold decrease in *K*_M_ not recapitulated by single-residue substitutions alone (Fig. S19 and S20). Community analysis of MD trajectories further suggested that introduction of the NL63 Loop 275 enhances inter-protomer dynamic coupling and coordinated motions between active sites (Fig. S21).

To place these HT-MEK^pro^-derived biochemical measurements in context with other methods, we compared HT-MEK^pro^-derived catalytic parameters with prior cellular and evolutionary fitness datasets. Correlations between HT-MEK^pro^-derived catalytic efficiency (*k*_cat_/*K*_M_) and yeast-based deep mutational scanning readouts depended strongly on the assay modality. Readouts based on cleavage of a single NSP4–5 site—measured either by loss of fluorescence resonance energy transfer (FRET) or by inactivation of a transcription factor driving GFP expression—showed weak correlations with *k*_cat_/*K*_M_ (Spearman ρ = 0.11 and 0.20, respectively; Fig. S22).^15^ In contrast, a growth-based assay, which leverages the toxicity of active M^pro^ variants to yeast, exhibited moderate agreement with *k*_cat_/*K*_M_ (Spearman ρ = 0.62; Fig. S22).^15^ Comparisons to phylogeny-inferred fitness showed more modest agreement (Spearman ρ = 0.43; Fig. S23).^40^ These results strongly suggest that while catalytic efficiency contributes to “fitness” as measured in these contexts, cellular and evolutionary readouts integrate additional constraints beyond intrinsic enzyme kinetics, and therefore capture overlapping but distinct aspects of protease function. Consistent with this result, many mutations with apparent biochemical effects *in vitro* appeared nearly neutral in cell-based assays, underscoring the complementary value of direct enzymological measurements.

We next quantified inhibitor sensitivity across the variant library by measuring IC_50_ values for the noncovalent inhibitor ensitrelvir.^32–34^ IC_50_ values spanned several orders of magnitude and arose from mutations distributed throughout M^pro^ structure, with enrichment near the active-site cavity (Fig. 3a,b). While single substitutions at established resistance sites reduced inhibitor potency, the most substantial resistance was observed in M^pro^ orthologs from other coronaviruses—IBV, NL63, 229E, FIPV, and PEDV—emphasizing that robust resistance often emerges from combinatorial sequence contexts, rather than isolated point mutations (Fig. 1g, Fig. 3a). We additionally identified previously unreported SARS-CoV-2 substitutions with increases in IC_50_ comparable to canonical variants: P52I, C44A, and N51Δ (deletion of N51) each conferred increases in IC_50_ similar to S144A and E166A (Fig. 3a). MD simulations of N51Δ and P52I, located on a loop adjacent to the “active site gateway”, a region implicated in conformational switching and substrate access, suggest that these mutations confer resistance by either increasing loop flexibility in N51Δ, or by favoring a more open conformation and alternative ensitrelvir positioning in P52I (Fig. S24).^41^ N51Δ and P52I are WT residues in NL63 M^pro^, which may contribute to this ortholog’s >10,000-fold higher IC_50_ for ensitrelvir (Fig. 1g). Notably, N51Δ and P52I have not emerged in SARS-CoV-2 passaging experiments with ensitrelvir, suggesting that they may have adverse collateral effects on other biochemical properties influencing viral replication (see below).

**Figure 3.**
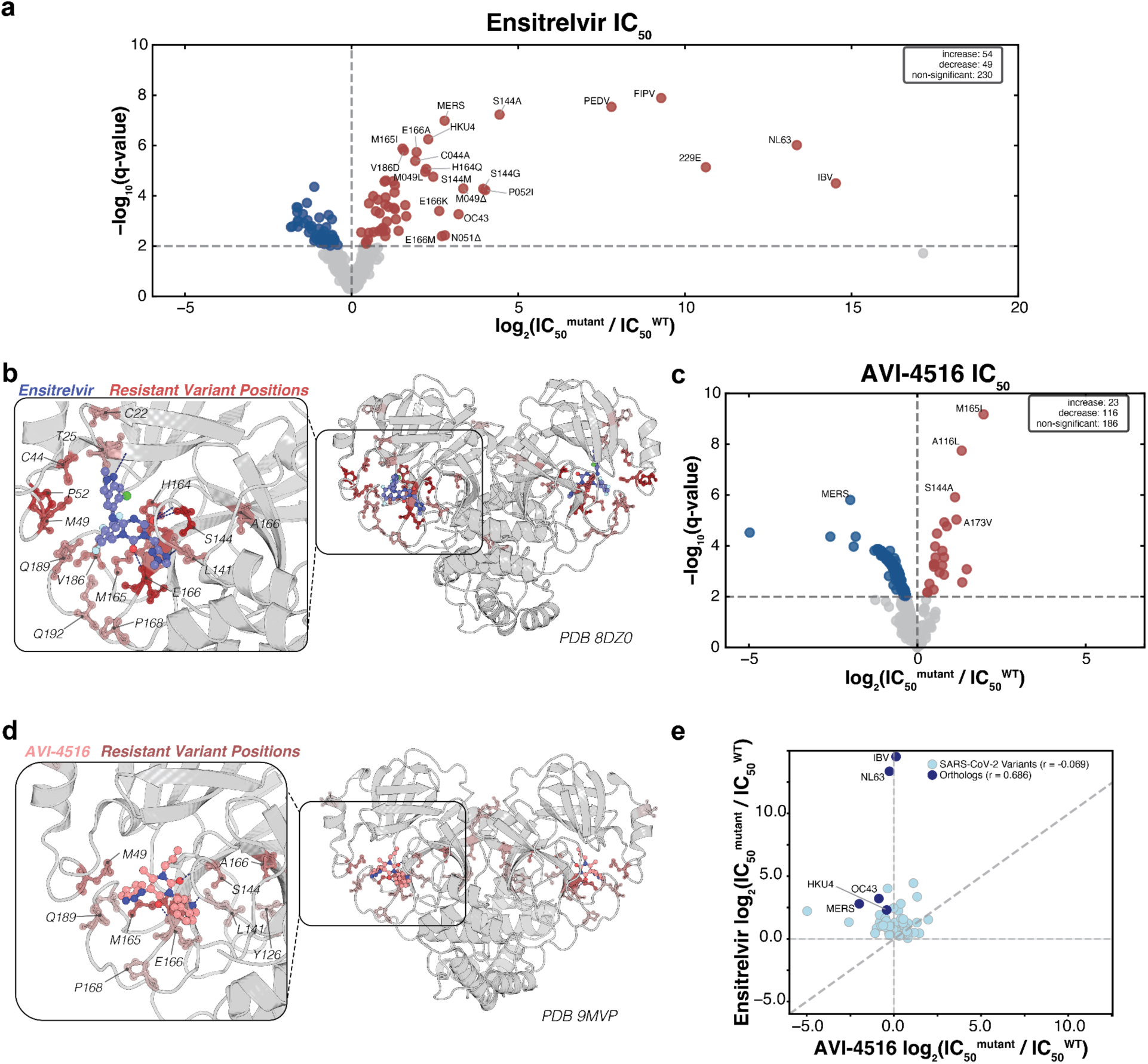
Inhibition landscapes of SARS-CoV-2 M^pro^ with two inhibitors. **(a)** Volcano plot showing variant effects on ensitrelvir IC_50_ relative to WT SARS-CoV-2 M^pro^. Points represent mean log_2_ fold changes in IC_50_ (Δlog_2_ IC_50_) from n = 6–9 biological replicates. Statistical significance of variant effects relative to WT was assessed using two-tailed Welch’s *t*-tests, followed by Benjamini–Hochberg correction for multiple hypothesis testing. Reported *q*-values represent FDR-adjusted *p*-values, with *q* < 0.01 considered significant. Only points with a coefficient of variation (CV = SD/mean across replicates) < 0.6 are shown to emphasize reproducible estimates. All values are reported in Tables S6 and S7. Variants at established resistance sites (M49, S144, E166) and newly identified hits (P52I, C44A, N51Δ) are labeled. For comparison, orthologs from other coronaviruses (e.g., IBV, NL63-CoV, 229E-CoV, FIPV, PEDV; Fig. 1f) are shown. **(b)** Structural mapping of resistance mutations onto the SARS-CoV-2 M^pro^ dimer bound to ensitrelvir (PDB 8DZ0). Residues are colored by Δlog_2_ IC_50_ according to the magnitude of the largest significant IC_50_ (≥ 2-fold, q < 0.01) effect observed at that position; the inset shows a zoomed active-site view with polar contacts to ensitrelvir as dashed lines. **(c)** Volcano plot showing variant effects on AVI-4516 IC₅₀ relative to WT, as in (a). Selected variants with significant resistance (≥ 2-fold increase, q < 0.01; CV < 0.6) are labeled. **(d)** Structural mapping of AVI-4516 resistance mutations onto SARS-CoV-2 M^pro^, colored by Δlog_2_ IC_50_ as in (b). **(e)** Comparison of resistance profiles for ensitrelvir and AVI-4516 across the variant library from (a) and (c). The thick dashed line denotes the 1:1 relationship. For clarity, only variants that increase IC_50_ for at least one inhibitor in (a) or (c) (red points) are shown

Prior studies with a limited number of variants have shown that nirmatrelvir and ensitrelvir exhibit distinct resistance profiles, reflecting their different active site interactions.^42^ Expanding the repertoire of inhibitors with non-overlapping resistance liabilities could broaden therapeutic options for SARS-CoV-2. To assess whether the resistance landscapes captured by HT-MEK^pro^ were sensitive to inhibitor chemotype, we additionally profiled the covalent inhibitor AVI-4516 (Fig. 3c and 3d).^43^ For four variants previously characterized with AVI-4516, on-chip IC_50_ values correlated closely with published measurements (R² = 0.96; Fig. S25).^43^ Across our variant library, AVI-4516 showed lower resistance liabilities than ensitrelvir, perhaps because its distinct active site interactions reduce susceptibility to common resistance pathways (Fig. 3e).

A major goal in antiviral development is identifying inhibitors with durable, pan-coronaviral activity. Notably, several M^pro^ orthologs that were moderately to highly resistant to ensitrelvir (including IBV, NL63, OC43, HKU4, and MERS) remained potently inhibited by AVI-4516 at levels comparable to SARS-CoV-2 M^pro^. These data underscore the pan-coronaviral potential of AVI-4516 and illustrate how HT-MEK^pro^ can identify inhibitor scaffolds with broad activity across evolutionary space, moving beyond the “one virus–one drug” paradigm.^43^

### Linking catalytic landscapes to viral fitness

Having mapped M^pro^ catalytic and resistance landscapes *in vitro*, we next asked how these biochemical properties relate to viral fitness. In principle, variants with severe catalytic defects should be strongly selected against in circulating viruses, whereas variants with near–WT or enhanced catalytic efficiency should be tolerated or even favored. Consistent with this expectation, substitutions that reduced catalytic efficiency (*k*_cat_/*K*_M_) as measured with HT-MEK^pro^ were rare in the GISAID database (https://gisaid.org/), the largest repository of COVID-19 sequences (Fig. 4a).^44^ However, this catalytic efficiency alone did not predict prevalence: several hyperactive variants identified by HT-MEK^pro^ were rare or absent in circulating viruses, suggesting additional constraints. For example, L50F, N151T, N142M, N142A, and S301F all increased catalytic efficiency on the NSP4-5 substrate, yet their clinical frequencies spanned orders of magnitude: L50F was reported 5,289 times, N151T 98 times, N142M 10 times, N142A once, and S301F not at all. Despite similar or enhanced catalytic efficiency on NSP4-5 *in vitro*, these variants differ dramatically in clinical prevalence, illustrating that NSP4-5 biochemical activity alone is insufficient to predict viral success.

**Figure 4.**
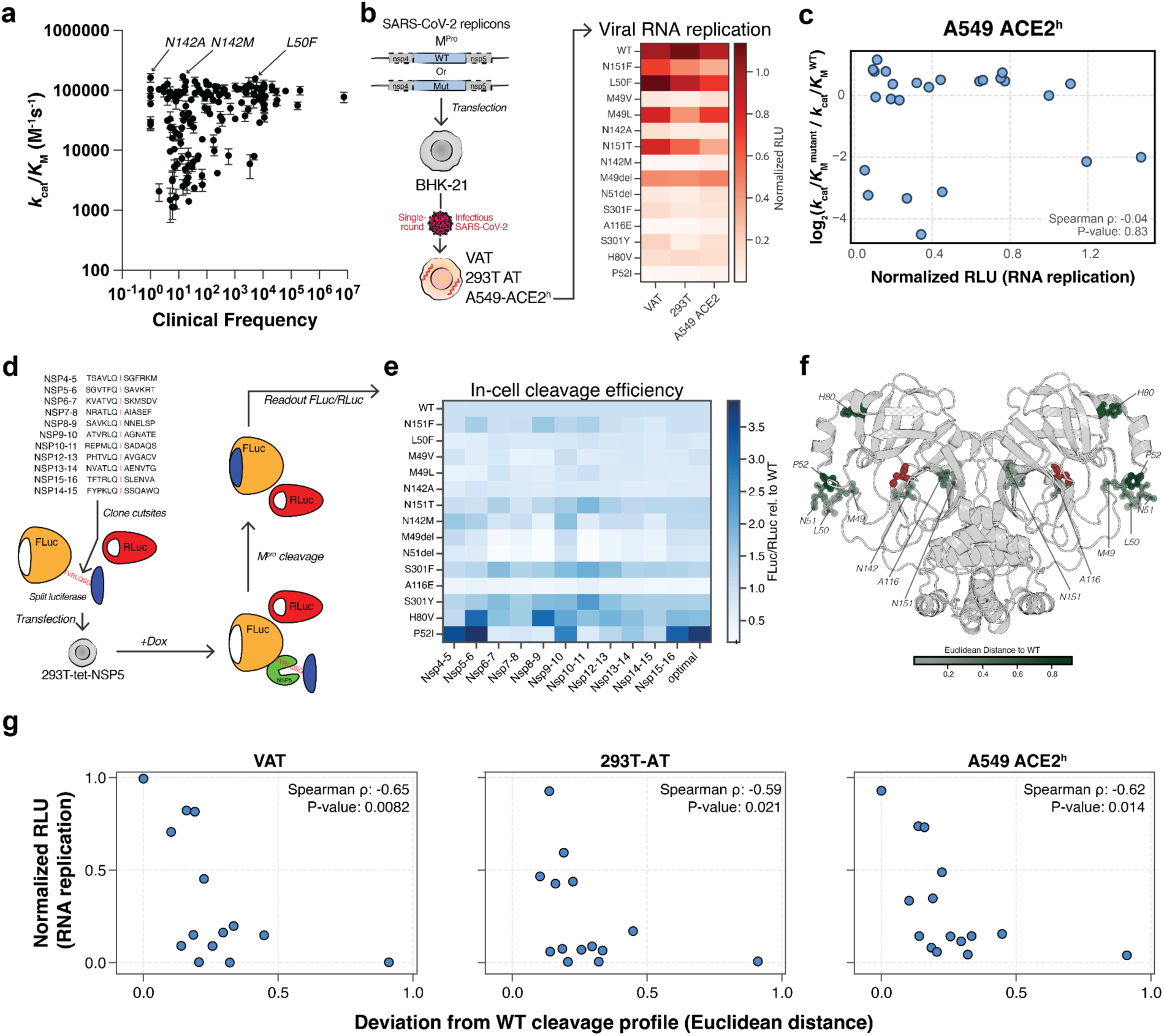
Linking catalytic landscapes of SARS-CoV-2 M^pro^ to viral fitness. **(a)** Relationship between M^pro^ catalytic efficiency (*k*_cat_/*K*_M_) measured by HT-MEK^pro^ and variant frequency in the GISAID database. **(b)** Replication efficiency of 14 M^pro^ variants in a SARS-CoV-2 replicon system. Variants were selected based on HT-MEK^pro^ measurements (either hyperactive or ensitrelvir-resistant). RNA replication was quantified in BHK-21 cells post-transfection and in VAT, 293T-AT, and A549-ACE2^h^ cell lines post-infection (see also Fig. S26). **(c)** Scatter plot of viral RNA replication in A549-ACE2^h^ versus catalytic efficiency (*k*_cat_/*K*_M_). Plots with other cell types versus *k*_cat_ and *K*_M_ can be found in Fig. S27. **(d)** Schematic of the in-cell cleavage assay, in which each viral polyprotein junction is inserted between the N- and C-terminal fragments of split firefly luciferase (FLuc); Renilla luciferase (RLuc) serves as an independent normalization control for expression and transfection. **(e)** Cleavage efficiency profiles across all eleven SARS-CoV-2 polyprotein junctions, plus one non-native “optimal” sequence,^46^ were measured using the assay in (d). Variants are ordered by their deviation from the WT profile, quantified as the Euclidean distance across the 11 junctions. **(f)** Structural mapping of cleavage-profile deviation (Euclidean distance from WT) onto the SARS-CoV-2 M^pro^ dimer (PDB 7CAM). The catalytic cysteine, C145, is shown in red. **(g)** Scatter plots showing the relationship between deviation from the WT cleavage profile (Euclidean distance) and relative RNA replication in VAT (Spearman ρ = –0.65, p = 0.0082), 293T-AT (Spearman ρ = –0.59, p = 0.021), and A549-ACE2^h^ (Spearman ρ = –0.62, p = 0.014) cells for each variant.

To directly test how these biochemical changes impact viral replication, we introduced a panel of 14 M^pro^ variants into a SARS-CoV-2 replicon system (Fig. 4b).^45^ Variants were selected based on HT-MEK^pro^ measurements: either hyperactive or conferred ensitrelvir resistance with catalytic efficiencies comparable to circulating mutations such as E166V. In BHK-21 cells, which report RNA replication independent of viral entry, particle production, and innate immune signaling, most variants supported RNA replication at or above WT levels (Fig. S26), indicating that they retain sufficient protease function and are not broadly cytotoxic.

In contrast, infection-based assays in three cell lines selected for graded capacity for interferon (IFN) signaling, from deficient to largely intact [Vero cells stably expressing ACE2 and TMPRSS2 (VAT), 293T cells stably expressing ACE2 and TMPRSS2 (293T-AT), and A549 cells stably expressing high levels of ACE2 (A549-ACE2^h^)], showed a wide range of infectivity as measured through an RNA replication-based luciferase reporter, with some variants similar to WT and others undetectable (Fig. 4b, Fig. S26).^45^ Across variants, viral replication correlated poorly with *k*_cat_, *K*_M_, or *k*_cat_/*K*_M_ measured with HT-MEK^pro^ on the canonical NSP4–5 substrate (all Spearman ρ < 0.4; Fig. 4c and Fig. S27). Replication phenotypes were nonetheless consistent across cell types, indicating that the observed defects reflect M^pro^-intrinsic constraints rather than cell-specific host responses (Fig. 4b).

One potential explanation for the disconnect between *in vitro* catalytic parameters and viral RNA replication is differences in assay substrate scope. Our initial HT-MEK^pro^ assays were performed with a substrate based on the NSP4-5 cleavage site; however, M^pro^ processes eleven distinct polyprotein junctions, each sharing a conserved P1 Gln but differing at surrounding positions (Fig. S28). Mutations that preserve activity on NSP4–5 while perturbing cleavage at other junctions could therefore disrupt viral polyprotein processing without reducing apparent catalytic efficiency in standard *in vitro* single-substrate assays.

To test this hypothesis, we measured cleavage efficiencies across all 11 SARS-CoV-2 polyprotein junctions in 293T cells using a split-luciferase reporter system (Fig. 4d).^46^ This assay reports the relative cleavage of each junction under cellular conditions, enabling the construction of variant-specific cleavage profiles. Several variants displayed marked deviations from the WT profile (Fig. 4e). Among these, P52I—located in the active-site gateway loop—showed the largest overall shift, consistent with MD simulations that show a more open loop conformation that may differentially affect substrate positioning across junctions (Fig. 4f, Fig. S24). Other variants with divergent profiles, such as H80V and S301Y, are located distal to the active site, consistent with allosteric modulation of the active-site conformational ensemble.

We next asked whether divergence in polyprotein cleavage profiles predicts viral replication. We quantified the divergence of each variant’s cleavage profile from WT using the Euclidean distance across junctions and compared this metric to each variant’s viral RNA replication. Indeed, cleavage-profile divergence correlated strongly with relative RNA replication in three cell lines: VAT (Spearman ρ = –0.65, p = 0.0082), 293T-AT (ρ = –0.59, p = 0.021), and A549-ACE2^h^ (ρ = –0.62, p = 0.014) (Fig. 4g).

Together, this in-cell and the above HT-MEK^pro^ data support a model in which viral fitness depends not on activity at a single junction, but on the coordinated balance of cleavage across all eleven, or a critical subset, of polyprotein sites. Variants that retain near–WT activity on NSP4–5 but disproportionately alter processing at other junctions can disrupt the ordered release of nonstructural proteins, impairing replication. Similar constraints have been observed in other viruses, such as Human Immunodeficiency Virus (HIV) and Semliki Forest virus, where the order and timing of polyprotein processing are critical for replication, and even subtle perturbations can attenuate infectivity.^47,48^ Consistent with this model, mass spectrometry studies of wild-type SARS-CoV-2 M^pro^ processing NSP7–11 suggest that cleavage proceeds in a defined sequence, perhaps to ensure proper assembly of the replication–transcription complex.^49,50^ These constraints are invisible to conventional single-substrate enzymology, where robust catalytic activity on the measured substrate can be maintained while altering specificities for others; however, they emerge when high-throughput biochemical measurements are integrated with cellular assays.

Beyond viral polyprotein processing, coronavirus M^pro^ enzymes cleave host factors: M^pro^ orthologs from multiple coronaviruses cleave NF-κB essential modulator (NEMO), dampening IFN signaling^51–54^, and SARS-CoV-2 M^pro^ has been reported to cleave >100 host substrates, including the IFN-stimulated lectin galectin-8 and gasdermin D, a pore-forming effector of antiviral pyroptosis.^55,56^ We therefore considered whether altered host substrate processing could substantially contribute to the replication phenotypes observed for specificity-altering M^pro^ variants. However, because many reported host M^pro^ substrates are involved in IFN signaling, we would expect replication phenotypes to vary with host IFN competence. Instead, infectivity defects were highly consistent across cell types spanning deficient, intermediate, and intact IFN signaling (Fig. 4b). Moreover, most variants supported robust RNA replication in the BHK-21 replicon system despite failing to sustain replication in infection-based assays, indicating preserved cellular viability (Fig. 4b, Fig. S26). Together, these observations are consistent with replication defects arising from impaired viral polyprotein processing than from altered host substrate cleavage or nonspecific cytotoxicity.

### High-throughput substrate specificity across polyprotein cleavage sites

Building on the observation that mutations can alter M^pro^ substrate specificity, we returned to HT-MEK^pro^ to systematically quantify how sequence variation reshapes polyprotein processing across the variant library. We extended HT-MEK^pro^ measurements to encompass all eleven SARS-CoV-2 polyprotein cleavage junctions (Fig. S28). Internally quenched peptide substrates spanning P6–P6′ of each NSP junction were assayed against our M^pro^ variant library (Fig. 5a, Table S9). For each variant, we measured initial cleavage rates for all junctions sequentially in a matched series at a single subsaturating substrate concentration (15 µM), resulting in more than 3,400 variant–substrate measurements. This design enables quantitative, variant-resolved comparisons of substrate preference across native substrate sequences under uniform biochemical conditions, a regime difficult to access with existing specificity-profiling approaches.

**Figure 5.**
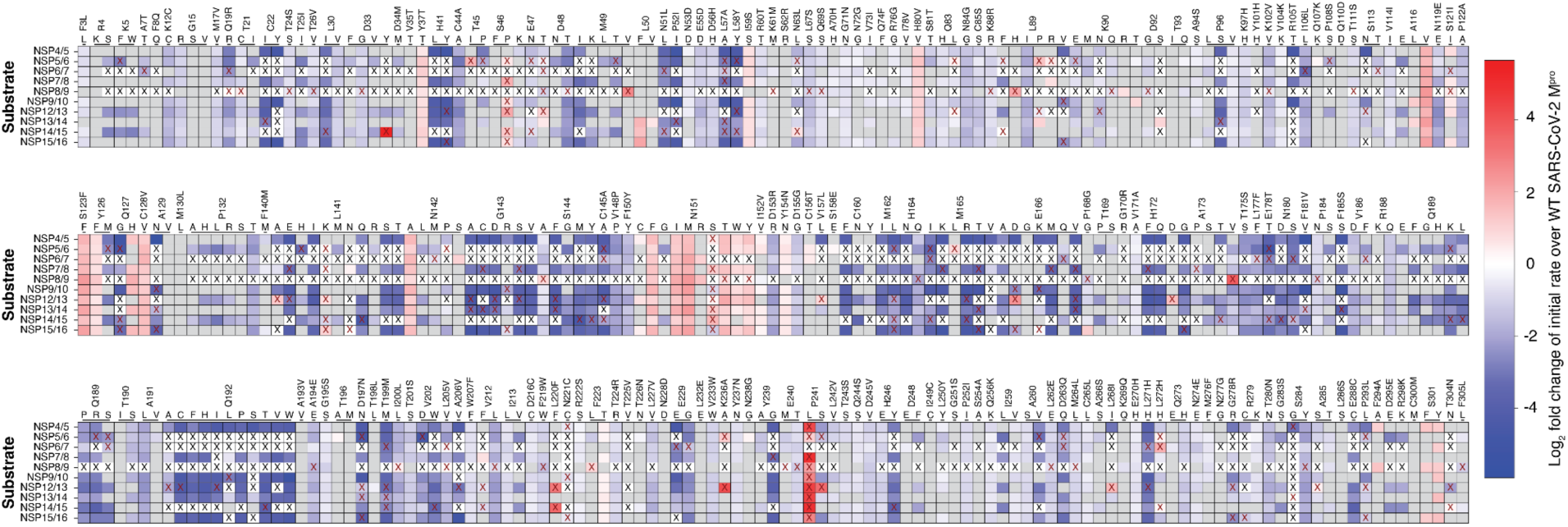
NSP cleavage specificity of SARS-CoV-2 M^pro^. Heatmap showing substrate-specificity profiles of SARS-CoV-2 M^pro^ variants across ten viral polyprotein junctions. Values represent log_2_ fold-changes in normalized initial cleavage rates relative to WT, measured at 15 μM peptide substrate (P6–P6′). Columns correspond to M^pro^ variants; rows correspond to NSP junctions. Colored cells indicate significant differences in initial rates between mutant and WT (p ≤ 0.05), with color intensity reflecting the magnitude and direction of the log_2_ fold-change (red, increased cleavage; blue, decreased cleavage). Grey cells indicate non-significant differences (p > 0.05). Red X denotes cases with only a single replicate for the mutant. Black X indicates undetectable initial rates above background for that mutant-substrate combination.

We used subsaturating substrate conditions because several NSP-derived peptides were solubility-limited, precluding full Michaelis–Menten titrations. Under these conditions, initial rates provide a proxy for relative specificity (*k*_cat_/*K*_MMut_ / *k*_cat_/*K*_MWT_), an assumption supported by the correlation between NSP4–5 initial rates under these conditions and the independently measured *k*_cat_/*K*_M_ values from above (Spearman ρ = 0.81, p-value = 2.8 × 10^-52^, Fig. S29). Consistent with prior reports, NSP8–9–derived peptides were cleaved far less efficiently than NSP4–5 by WT M^pro^ (∼3 orders of magnitude lower *k*_cat_/*K*_M_), and most variants exhibited slowed or undetectable NSP8–9 activity (Fig. 5a). Notable exceptions included A116V and S301Y, which showed increased activity on NSP8–9 (Fig. 5a). Under on-chip conditions, WT M^pro^ showed no detectable activity toward NSP10–11; however, several variants (e.g., L50F, A116V, N151M, and S301Y) exhibited significant NSP10–11 cleavage above background (Fig. S30).

Across the library, multiple mutations altered substrate preferences at one or more junctions, and these specificity-altering substitutions were distributed throughout the structure, including many distal from the active site (Fig. 5b). Notably, P52I and H80V—identified in cell-based assays as having the most divergent cleavage profiles—also ranked among the most shifted variants in the in vitro specificity landscape (Fig. 5a; Fig. S31), and additional variants selected based on HT-MEK^pro^ profiles similarly showed junction-specific effects in cells, with S123F and Q127H exhibiting broadly elevated but biased cleavage and Q127G showing reduced activity, particularly outside NSP4-5 (Fig. S32).

Finally, having established that mutational perturbations of substrate specificity constrain viral fitness, we asked whether similar effects could be achieved pharmacologically. The specificity-altering substitutions S301Y and S301F, which reduced viral replication and reshaped both in-cell and *in vitro* cleavage profiles (Figs. 4e and 5), map to an allosteric pocket between domains 2 and 3, approximately 15 Å from the catalytic cysteine (Fig. S33a), previously identified in an X-ray drug-repurposing screen as a binding site for the kinase inhibitor pelitinib.^57^ *In vitro* assays showed that pelitinib caused no detectable inhibition of WT M^pro^ activity on the NSP4–5 substrate *in vitro* (Fig. S33b), indicating that it does not act as a conventional active-site inhibitor. In contrast, in-cell cleavage assays revealed a biased cleavage profile distinct from untreated WT, suggesting that pelitinib selectively perturbed cleavage at specific polyprotein junctions (Fig. S34). Notably, pelitinib had minimal effect in the S301Y background, consistent with occlusion of the binding pocket. In contrast, its effects were amplified in the specificity-altering P52I variant, which lies at the active-site gateway and exhibits one of the largest shifts in cleavage profiles (Fig. 4e, Fig. S34). Together, these results suggest that small-molecule engagement of this allosteric site can modulate proteolytic balance and timing without directly inhibiting catalysis, mirroring the effects of specificity-altering mutations identified through HT-MEK^pro^. Extending similar mutational maps to other proteases could reveal sites that are pharmacologically targetable to modulate catalytic activity, reprogram specificity, or rescue function in disease-associated mutants, analogous to classic chemical rescue strategies.^58^

## Conclusions and Implications

HT-MEK^pro^ overcomes the scale and resource limitations of traditional protease enzymology, providing a broadly applicable approach for the systematic and quantitative mapping of protease catalytic, inhibitory, and specificity landscapes at an unprecedented depth. Applied to SARS-CoV-2 M^pro^, this approach reveals activating and resistance mutations as well as distinct resistance profiles across inhibitor chemotypes. Notably, our initial library includes hundreds of clinically observed SARS-CoV-2 M^pro^ variants, making this the first instance in which HT-MEK has been applied to clinically circulating variants rather than engineered or purely evolutionary substitutions. Quantitative maps of catalysis and inhibition like these can provide a foundation for anticipating resistance and guiding the design of next-generation antivirals. The observation that AVI-4516 remains potent across evolutionarily divergent coronavirus M^pro^ orthologs underscores how HT-MEK^pro^ can be used prospectively to identify inhibitor scaffolds with pan-coronaviral potential before resistance emerges in the clinic. As new M^pro^ inhibitors are developed for SARS-CoV-2 and related coronaviruses, systematic identification of both observed and latent resistance mutations will be critical for assessing durability and achieving pan-coronavirus activity, particularly given the repeated emergence of pathogenic coronaviruses in the 21st century.^59^

Beyond identifying resistance mutations, HT-MEK^pro^ can map how substitutions reprogram extended substrate specificity—beyond the P1–P1′ positions—across all eleven polyprotein junctions. These substrate specificity maps capture effects of hundreds of protease mutations, including distal substitutions, in a way not accessible through traditional specificity-profiling methods. Prior work on VOC M^pro^ mutants reported mutation-dependent changes in substrate specificity using HyCoSuL, including broadened P2 permissivity for specific mutants (e.g., A260V, S284G).^9^ Here, we extend these observations by measuring relative cleavage of all native polyprotein cut sites in parallel, providing a systematic view of how viral evolution reshapes substrate specificity and, consequently, proteolytic timing across hundreds of variants sampled from circulating viruses and evolutionary space. More broadly, this framework offers a general strategy for identifying mutations that bias protease selectivity and for pinpointing sites at which specificity may be engineered or pharmacologically modulated.

Looking forward, HT-MEK^pro^-derived biochemical landscapes provide not only mechanistic insight but also quantitative training data for prediction. The scale of the HT-MEK^pro^ dataset can enable data-driven modeling of protease sequence–function relationships. Predicting how amino-acid substitutions alter *k*_cat_, *K*_M_, or IC_50_ has been challenging in part because most available training data are sparse, indirect, and heterogeneous across assays and conditions.^29^ While large IC_50_ datasets spanning inhibitor panels for reference sequences exist,^60–63^ analogous datasets spanning hundreds of protein variants are rare, limiting quantitative, variant-aware resistance prediction. By contrast, HT-MEK^pro^ provides hundreds of variant-resolved ground-truth biochemical constants collected under identical experimental conditions. As an initial proof of concept, both lightweight models and a supervised neural network trained on HT-MEK^pro^-derived measurements utilizing protein language model embeddings were moderately predictive of *k*_cat_ and IC_50_, with performance improving both as the training set size increased and with the neural network architecture (Fig. S35). While further expansion of datasets and exploration of improved model architectures will be needed for robust quantitative resistance prediction, these results establish a foundation for predictive frameworks aimed at anticipating resistance mutants or engineering proteases with customized catalytic properties.

Together, this work establishes high-throughput protease enzymology using HT-MEK^pro^ as a powerful complement to cellular assays, enabling quantitative dissection of sequence–function–fitness relationships that are inaccessible to either approach alone. By linking variant-resolved enzymology to viral replication, HT-MEK^pro^ reveals how catalytic efficiency and substrate specificity constrain viral fitness. More broadly, this framework provides a general strategy for mapping and modeling enzyme sequence–function–fitness landscapes, offering a comprehensive framework for uncovering biochemical constraints on biological systems and guiding both antiviral strategies and enzyme engineering.

## Acknowledgements

We thank members of the Pinney laboratory for their discussions and comments on the manuscript. We thank members of the Coyote-Maestas laboratory for experimental advice. We thank members of the Quantitative Biosciences Institute Coronavirus Research Group (QCRG) AViDD Program for their guidance and feedback throughout this work. M.M.P. and M.O. are Biohub, San Francisco, Investigators. For the analysis of SARS-CoV-2 sequences from the GISAID database, we gratefully acknowledge all data contributors, i.e., the Authors and their Originating laboratories responsible for obtaining the specimens, and their submitting laboratories for generating the genetic sequence and metadata and sharing them via the GISAID Initiative, on which these analyses were based.

## Author Contributions

N.J.Y., C.S.C., T.Y.T., and M.M.P. designed the study. N.Y. and G.P.R.A. cloned and prepared the M^pro^ library for HT-MEK^pro^ experiments. D.A. synthesized quenched fluorescent peptide substrates. N.J.Y., W.V.T.V., and D.A. performed *in vitro* experiments on and off-device for SARS-CoV-2 M^pro^ and orthologs. S.C. and D.A. performed *in vitro* off-device experiments for MMP12, CMV, and matriptase. N.J.Y., D.F.M., N.F., and D.A. performed HT-MEK^pro^ data analysis. D.A. and S.C. performed downstream computational and statistical analysis. D.A. performed machine-learning and deep-learning experiments. J.R. and A.K. performed viral replicon and in-cell cleavage efficiency assays under the supervision of M.O. and T.Y.T. D.A. and M.M.P. assembled figures and wrote the manuscript with input from all authors.

## Data and materials availability

Summary tables of kinetic and inhibition parameters for each M^pro^ variant are provided in the Supplementary Materials. Full specificity datasets, along with the complete kinetic and inhibition parameters, are available at Zenodo (DOI: 10.5281/zenodo.18382886).

## List of Supplementary Materials

Materials and Methods

Tables S1 to S10

Figs. S1 to S35

## Materials and Methods

### Reagents

All reagents were of the highest purity commercially available (≥97%). Unless otherwise specified, chemicals were purchased from MilliporeSigma or Thermo Fisher Scientific. Q5 High-Fidelity DNA polymerase, restriction enzymes, and ligases were from New England Biolabs (NEB). PURExpress in vitro translation components were from NEB. Plasmid miniprep kits were from Qiagen (QIAprep Spin Miniprep Kit) or New England Biolabs (Monarch Plasmid Miniprep Kit), and 96-well plasmid purification was performed using the Wizard SV 96 Plasmid DNA Purification System (Promega). All oligonucleotides were synthesized by Integrated DNA Technologies or Twist Biosciences.

### Cloning, Mutagenesis, and Plasmid Purification

PCR fragments were amplified using Q5 polymerase (NEB) according to the manufacturer’s instructions. An expression vector containing a T7 promoter served as the cloning backbone for all protease–eGFP fusion constructs. Inserts were assembled into the backbone using Gibson assembly or T4 DNA ligase–mediated cloning and then transformed into chemically competent DH5α cells (NEB) via heat shock.

Plasmids used for off-device expression were prepared with QIAprep or Monarch miniprep kits. Plasmids used for on-device HT-MEK^pro^ experiments were purified in 96-well format using the Wizard SV 96 Plasmid DNA Purification System (Promega). All constructs were sequence-verified by Sanger sequencing (Genewiz) or nanopore sequencing (Primordium Labs or Plasmidsaurus Inc.).

### Cells

BHK-21 and 293T cells were obtained from ATCC and cultured in standard media: DMEM (Corning) supplemented with 10% fetal bovine serum (FBS) (GeminiBio), 1x Glutamax (Corning), 1x non-essential amino acids (NEAA) (Corning), and 1x penicillin-streptomycin (Corning) at 37 °C, 5% CO_2_. Vero cells stably expressing human ACE2 and TMPRSS2 (VAT) (gifted from A. Creanga and B. Graham at NIH) were maintained in standard media with the addition of 10 μg/mL of puromycin. 293T cells stably expressing ACE2 and TMPRSS2 (293T-AT) have been described previously^64^ and were maintained in standard media with 10 μg/mL blasticidin and 250 μg/mL hygromycin. A549 cells stably expressing high levels of ACE2 have been previously described and were maintained in standard media supplemented with 10 μg/mL blasticidin.^43,65^ All cell lines are confirmed mycoplasma-free by quarterly testing.

### Peptide Synthesis of HT-MEK^pro^ Substrates

The peptide substrates were synthesized as described previously^43^ on a Syro II peptide synthesizer (Biotage) using standard N-(9-fluorenylmethoxycarbonyl) (FMOC) solid-phase synthesis. All amino acids and HCTU were purchased from Anaspec, and all other chemicals and solvents were purchased from Sigma-Aldrich. All the peptides were synthesized using 12 µmol of preloaded lysine(DNP) resin at ambient temperature. All coupling reactions were done with 4.9 eq of O-(1H-6-chlorobenzotriazole-1-yl)-1,1,3,3-tetramethyluronium hexafluoro-phosphate (HCTU), 5 eq of FMOC-AA-OH, and 20 eq of N-methylmorpholine (NMM) in 500 μL of N, N-dimethyl formamide (DMF). Each amino acid position was double-coupled with 8-minute reactions. FMOC-protected N-termini were deprotected with 500 μL 40% 4-methypiperidine in DMF for 3 minutes, followed by 500 μL 20% 4-methypiperidine in DMF for 10 minutes and six washes with 500 μL of DMF for 3 minutes. After solid-state peptide synthesis, peptides were cleaved from the resin using a cleavage solution (75% TFA, 5% Phenol, 5% Triisopropylsilane, 5% H 2 O, 5% Anisol, 5% Thioanisol) at ambient temperature with shaking for 3 h before precipitation with 45 mL of ice-cold 1:1 diethyl ether and hexane. The suspension was pelleted via centrifugation for 20 minutes at 4,000 × g, and the supernatant was decanted. The resulting pellet was allowed to dry overnight at ambient temperature. The crude pellet was dissolved in 200 µL of dimethyl sulfoxide (DMSO). Each peptide was purified via high-performance liquid chromatography (HPLC) on an Agilent Pursuit 5 C18 column (5 mm bead size, 150 × 21.2 mm) using an Agilent PrepStar 218 series preparative high-performance liquid chromatography suite. The mobile phase consisted of water (0.1% TFA) and an increasing gradient of acetonitrile (0.1% TFA) from 20% to 80%. The solvent was removed from the purified material through lyophilization using the Freezone 2.5L lyophilizer and confirmed by LC-MS (Agilent 1260 Infinity II).

### Selection and Construction of Protease Variant Libraries

#### SARS-CoV-2 M^pro^ point mutations

To map the catalytic and resistance landscapes of SARS-CoV-2 M^pro^, we generated a library of 397 variants spanning functional, clinical, and evolutionary sequence space (Fig. 2a, Fig. S13–16). The first half of the library emphasized functional and clinically relevant substitutions, including 136 mutations from variants of concern (VOCs), variants of interest (VOIs), and high-frequency clinical isolates. It targeted Hamming-distance scans of residues directly involved in substrate or inhibitor recognition (e.g., M49, N142, G143, S144, M165, E166, R188, Q189, T190). To complement these, we incorporated 52 top-ranked mutations from a published yeast-based selection screen, prioritizing substitutions with potential hyperactivating effects, as increased catalytic activity represents a possible resistance mechanism beyond decreased inhibitor binding.^15^

The second half of the library focused on evolutionary comparisons, introducing NL63-derived sequence features into SARS-CoV-2 M^pro^ through a combination of targeted single-residue substitutions, multi-residue swaps, and combinatorial loop designs (Fig. S14). NL63 M^pro^ was chosen because we found it was naturally hyperactive on the SARS-CoV-2 NSP4-5 substrate and resistant to ensitrelvir (Fig. 1g), providing a valuable evolutionary comparison for dissecting the molecular basis of enhanced catalysis and natural resistance. This evolutionary-focused library subset included single amino acid swaps, loop exchanges, and combinatorial loop designs targeting active-site loops, enabling us to probe both discrete and synergistic contributions of NL63 sequence elements to SARS-CoV-2 M^pro^ function.

SARS-CoV-2 M^pro^ variants were generated using Deep Indel Missense Programmable Library Engineering (DIMPLE), a Golden Gate–based pipeline for pooled deep mutational scanning libraries with barcoding.^66^ Briefly, the 918-bp M^pro^ coding sequence (306 amino acids) was divided computationally into six overlapping gene fragments, each flanked by unique primer barcodes and BsaI recognition sites for assembly. The designed oligo pool contained 6,120 unique oligos. We filtered this pool to retain only oligos corresponding to variants in our targeted library.

Each sub-pool was PCR-amplified from the oligo pool (Twist Biosciences) using sub-pool–specific primers. In parallel, the parental M^pro^–eGFP backbone plasmid was amplified into six overlapping fragments, each lacking the region replaced by the corresponding oligo sub-pool. Golden Gate assembly using BsaI and T4 DNA ligase was then used to reconstruct pooled M^pro^–eGFP plasmids containing the desired mutations.

For HT-MEK^pro^, individually isolated plasmids were required. Pooled libraries were transformed into competent cells, plated at low density, and single colonies were picked using an automated colony picker (QPix XE, Molecular Devices). Colony picking, miniprepping using the Wizard SV 96 Plasmid DNA Purification System (Promega), and Sanger sequencing were performed in plate format. Sequence-verified variants were then arrayed into library plates for use in on-device experiments.

#### Loop swaps and combinatorial variants

Loop swap and deletion constructs compatible with the DIMPLE strategy (fully contained within a single fragment) were designed manually and incorporated into the appropriate oligo sub-pools. Loop swaps spanning multiple DIMPLE fragments and multi-site combinations that could not be encoded within a single oligo were generated using standard site-directed mutagenesis or Gibson assembly, followed by sequence verification.

#### Coronavirus M^pro^ orthologs

M^pro^ orthologs from 14 coronaviruses spanning alpha, beta, gamma, delta, and epsilon genera and diverse hosts (e.g., human, porcine, feline, avian, murine, bat) were synthesized as codon-optimized genes (Twist) and inserted into the M^pro^–eGFP backbone using Gibson assembly. All ortholog constructs contained the same C-terminal (G_4_S)_5_–eGFP fusion and Q306A substitution used for SARS-CoV-2 M^pro^ to prevent C-terminal self-cleavage, and were sequence-verified by Sanger sequencing (Elim Biopharmaceuticals Inc.). Of the fourteen orthologs, twelve expressed and were active above background.

#### Other proteases

The human cytomegalovirus (CMV) serine protease, human serine protease matriptase, and human metalloprotease MMP-12 were cloned into the same expression backbone used for M^pro^, with C-terminal (G_4_S)_5_–eGFP fusions. Constructs were assembled by Gibson cloning and sequence-verified. Sequences of these other proteases are listed in Table S10, along with their tested substrate sequences. All substrates were synthesized similarly to the SARS-CoV-2 substrates (see *Michaelis–Menten Kinetics on the HT-MEK^pro^ Platform: Substrate preparation*, below).

### Off-Device *in vitro* expression and characterization of protease-eGFP fusions

Protease–eGFP fusions were expressed in vitro using the PURExpress In Vitro Protein Synthesis Kit (NEB). Reactions (25 μL total) contained 10 μL Solution A, 7.5 μL Solution B, RNasin Ribonuclease Inhibitor (Promega), and 50-250 ng plasmid DNA. For activation testing, the reaction mix was supplemented with water, TEV protease, or enterokinase as appropriate. Reactions were incubated at 37°C for 2–3 h and then held at 4°C to allow eGFP maturation.

Fluorescent protease–eGFP yields were quantified on a plate reader (BioTek NEO2 or Synergy H1, Agilent) using excitation/emission settings appropriate for eGFP, and concentrations were calculated from a standard curve of purified eGFP. Expression and activation states were assessed by SDS–PAGE followed by imaging on a Bio-Rad ChemiDoc MP system.

### Off-device pelitinib inhibition assays

Pelitinib (HY-32718, MedChem Express) was tested off-device under the same buffer conditions as those used for on-chip NSP substrates. Cleavage of internally quenched fluorogenic peptides derived from NSP4-5 was monitored at 30 μM substrate in the presence of pelitinib across a titration spanning and exceeding its reported EC50 (1.25 μM).^57^ Concentrations tested were 0, 0.94, 1.88, 3.75, 7.5, 15, and 30 μM pelitinib. All reactions were performed with a constant 2% DMSO concentration. Each condition was assayed in triplicate, with two WT DMSO controls per plate, over a 4-h reaction at 37°C. Initial rates were determined from the linear regime of fluorescence increase.

### Adaptation of High-Throughput Microfluidic Enzyme Kinetics (HT-MEK) Platform for proteases (HT-MEK^pro^) and Specifications

#### Photolithographic mold fabrication

Briefly, flow- and control-layer master molds for PS1.8K MITOMI 2.0 PDMS devices (1792 chambers; design files at https://www.fordycelab.com/microfluidic-design-files) were fabricated on 4-inch, single-sided silicon test-grade wafers (University Wafer) using 30,000 DPI transparency masks (Fineline Imaging), following standard photolithography procedures described previously for MITOMI 2.0 device molds.^28,67–69^

#### PDMS device fabrication

Two-layer “push-down” MITOMI devices were fabricated using RTV615 PDMS (Momentive; batch RTV615 010-Pail Kit No. 23BWFA015) as described previously.^28,67–69^ For the control layer, PDMS was mixed at a 5:1 (A: B) ratio, poured over the control-layer master, degassed, and baked at 80°C for 40 min. For the flow layer, 2” × 3” glass slides (Corning) were spin-coated with a 20:1 (A: B) PDMS mixture (10 s at 500 rpm, then 75 s at 1820 rpm on a Laurell WS-400B-6NPP/LITE spinner) and baked at 80°C for 40 min. All fabrication was performed in a laminar flow hood (AirScience Purair FLOW-36) to minimize particulate contamination. The partially cured control layer was aligned and bonded to the flow layer, then fully cured at 80°C. Inlet and outlet holes were punched before bonding to the glass.

#### Device layout and valve architecture

Each PS1.8K device contains 1792 chambers arranged in a 32 × 56 array. Every chamber is divided into a DNA compartment and a reaction compartment, separated by a Neck valve. Chambers are isolated from neighbors by Sandwich valves, and a Button valve in each reaction compartment enables on-chip surface patterning and enzyme immobilization. The shared flow layer allows sequential delivery of cell-free expression components, activation protease, fluorescent antibody, and substrates.

#### Plasmid Arraying and Device Alignment

Sequence-verified plasmids encoding M^pro^ variants, orthologs, and additional proteases were normalized to 50–400 ng/μL in printing buffer and arrayed in a 32 × 56 format in 384-well plates. Arrays were printed onto PDMS-coated quartz slides using a SciFLEXARRAYER S3 piezoelectric arrayer (Scienion). Spot quality and cross-contamination were assessed by test prints and fluorescence imaging of control spots. Devices were aligned over the printed plasmid arrays using a stereomicroscope and baked at 80°C to bond PDMS to the glass slide and isolate the DNA spots within the DNA compartments of each chamber.

#### Imaging

Fluorescence imaging was performed on a Nikon Ti2-E inverted microscope equipped with a Lumencor Spectra light source, a Photometrics Kinetix camera, and appropriate filter sets for eGFP, DyLight550, and 5-FAM. For each time point, a 5 × 5 raster of images was acquired and stitched into a whole-chip image (WCI) using custom Python code. Exposure times were typically 1–5 ms for eGFP and DyLight550 channels and 5–30 ms for 5-FAM.

#### Surface Patterning, On-Device Protease Expression, and Activation

Devices were passivated with ultrapure BSA (Invitrogen, AM2618) and patterned with neutravidin (NA, Thermo Scientific catalog no. PI31050) and a biotinylated anti-eGFP nanobody (ProteinTech, catalog no. GTB-250), as described previously.^29^ PURExpress components were flowed into the DNA chambers, Neck and Sandwich valves were closed to isolate DNA and reaction compartments, and cell-free expression was carried out at 37°C for 2–3 h. Protease–eGFP fusions were captured within the reaction chambers by pre-patterned anti-GFP nanobodies under the Button valve.

We evaluated several activation strategies to facilitate M^pro^ maturation, including auto-activation and orthogonal activation using enterokinase (EK) or Tobacco Etch Virus (TEV) protease (Fig. S4a–f). Auto-activation achieved high but incomplete cleavage of WT M^pro^ and, as expected, no detectable activation of the catalytically inactive C145A mutant (Fig. S4a-d). Because the extent of auto-activation depends on both enzyme yield and intrinsic activity, it will produce inconsistent maturation across variant libraries. Supplementary activation with an unrelated protease provides a more reliable, variant-independent solution. EK achieved near-complete activation but introduced unintended secondary cleavage products (Fig. S4c,e). By contrast, TEV protease produced complete, off-target–free activation of both WT and C145A M^pro^ (Fig. S4c,f), and TEV-activated WT M^pro^ showed robust activity on a commercial fluorogenic substrate (Fig. S5). The activation step was performed at 37°C for 90 min to allow complete cleavage at the engineered N-terminal site and the generation of the native Ser1 N-terminus for M^pro^. Activation was assessed by flowing DyLight550-conjugated anti-FLAG antibody, imaging using the compatible channel, and comparing signal across constructs.

The identity of the N-terminal tag also strongly influenced on-device expression levels: larger tags (e.g., GST) abolished expression, while progressively smaller tags improved yields (Fig. S3). A minimal FLAG tag coupled to the TEV site supported high on-device expression, efficient activation, strong catalytic activity, and inhibition by nirmatrelvir (Fig. 1c, Fig. S6–8, Table S1).

### Michaelis–Menten Kinetics on the HT-MEK^pro^ Platform

#### Substrate preparation

Internally quenched fluorogenic peptide substrates spanning P6–P6′ of NSP4-5 and each of the other 10 SARS-CoV-2 NSP junctions were synthesized internally with N-terminal 5-carboxyfluorescein (5-FAM) dye and C-terminal dinitrophenolate (dnp) quencher group. Sequences for these substrates can be found in Table S9. Substrates were dissolved in DMSO and diluted into assay buffer (50 mM Tris, 150 mM NaCl, 1 mM EDTA, 1 mM TCEP, 0.05% Tween-20, pH 7.4) such that the final DMSO concentration was identical across all conditions in a run at 2%.

#### On-device Michaelis–Menten measurements

For Michaelis–Menten experiments, a 10-point, ∼1.8-fold serial dilution of substrate (1–200 μM) was prepared in assay buffer. For each concentration, substrate was flowed through the device with Button valves closed for 5 min to exchange solution in the flow layer, followed by a 10-min activity assay with Button valves open and Sandwich valves closed to isolate the chambers. Time-lapse images (whole-chip) were acquired every few seconds over the reaction period.

Background-subtracted 5-FAM fluorescence within each chamber was converted to product concentration using a per-device 5-FAM standard curve. Initial rates were extracted from the linear regime for each substrate concentration and chamber. For each variant, rates across 6–9 independent chambers were fit to the Michaelis–Menten equation to obtain *k*_cat_, *K*_M_, and *k*_cat_/*K*_M_, and parameters were averaged across chambers. The full M^pro^ library was measured across two devices; WT and 12 reference variants were included on both for reproducibility assessment. Michaelis–Menten parameters (*k*_cat_, *K*_M_, and *k*_cat_/*K*_M_) measured for twelve shared variants across independent experiments were strongly correlated (R^2^ = 0.87-0.97; Fig. S17).

### Inhibition Assays on the HT-MEK^pro^ Platform

#### Ensitrelvir

On-chip inhibition assays for ensitrelvir were performed at 30 μM NSP4-5 substrate. A 12-point gradient of ensitrelvir concentrations (20 pM–90 μM) was flowed sequentially through the device. For each inhibitor concentration, the device was perfused for 5 minutes with the Button valves closed, followed by a 10-minute activity measurement with the Button valves open, as described above. Initial rates at each concentration were determined and normalized to the DMSO control. Percent inhibition was fit to a sigmoidal model to obtain IC50 values per chamber and per variant. IC50 values were averaged across 6–9 replicates per variant. On-device IC50 values for WT and canonical resistance mutants (S144A, H164Q, E166M, E166V) showed excellent agreement with off-device measurements (Fig. 1e, R^2^ = 0.92, slope = 0.91).

#### AVI-4516

Covalent inhibition by AVI-4516 was assessed similarly, using a 12-point concentration series (20 pM–90 μM) at a 30 μM NSP4-5 substrate. After initial filtering of outliers (see below), dose–response curves were fit to obtain IC_50_ values. Agreement between on- and off-device measurements for WT M^pro^ validated the quantitative performance of covalent inhibitors (Fig. S25, R^2^ = 0.96).

### Substrate Specificity Assays on the HT-MEK^pro^ Platform

For specificity profiling, substrates corresponding to each of the 11 NSP junctions (P6–P6′) were assayed at a single subsaturating concentration (15 μM) across the library of 328 catalytically active M^pro^ variants. Reactions were performed in assay buffer (50 mM Tris, 150 mM NaCl, 1 mM EDTA, 1 mM TCEP, 0.05% Tween-20, pH 7.4) containing 2% DMSO.

For each NSP substrate, the solution was flowed through the device for 5 minutes with the Button valves closed, followed by a 15-minute kinetic acquisition with the Button valves open and the Sandwich valves closed. Whole-chip eGFP and 5-FAM images were stitched and background-subtracted using images acquired before expression (eGFP) or before substrate addition (5-FAM). Button and chamber locations were identified by interpolating the 32 × 56 array from four manually picked corner chambers. These rough positions were refined using opencv.HoughCircles, and then summed pixel intensity was obtained across all images within each chamber or button, then converted to [NADPH] (μM) or [eGFP] (nM) using the appropriate standard curve (the eGFP standard curve was measured in the manner of Markin et al. ^28^).

Summed pixel intensities were converted to enzyme (eGFP) and product (5-FAM) concentrations using per-device calibration curves. A lower limit of detection for enzyme concentration (typically 1–2 nM) was imposed; chambers below this threshold were excluded. Initial rates were extracted for each chamber and substrate, and rates were normalized by enzyme concentration to approximate *k*_cat_/*K*_M_ under subsaturating conditions. NSP4-5 rates from these single-concentration experiments correlated well with independently measured *k*_cat_/*K*_M_ values from full Michaelis–Menten fits, supporting this approximation (Spearman ρ = 0.81, p-value = 2.8 × 10^-52^, Fig. S29).

### Euclidean Distance Calculations

#### In-cell cleavage profiles

For in-cell split-luciferase cleavage assays (Fig. 4d), relative cleavage efficiencies for each variant across peptide substrates containing the 11 polyprotein junctions were normalized to the WT value for each substrate. To quantify the overall divergence of a variant’s cleavage profile from WT, we computed the Euclidean distance in 11-dimensional junction space using the Euclidean function from scipy.spatial.distance in Python.

### Inhibitor Data Filtering and Plotting

For ensitrelvir and AVI-4516, raw on-device IC_50_ values and enzyme expression levels were first filtered based on Z-scores. For each variant, per-chamber enzyme concentrations were converted to Z-scores; chambers with an absolute Z-score greater than 1.5 were removed. From the remaining chambers, IC_50_ values were converted to Z-scores (log10-transformed) and outliers (|Z| > 3.5) were excluded. Variants with fewer than three remaining replicates were discarded. Finally, the coefficient of variation (CV) across replicates was calculated, and variants with a CV greater than 0.6 were excluded from the volcano plots. All values, including filtered variants, are reported in Tables S6-S8.

### Heatmap Generation and Statistical Analysis for Specificity Profiles

For each NSP substrate and variant, the normalized initial rates (product concentration per time per enzyme concentration) were averaged across chambers after filtering (enzyme concentration≥ 1–2 nM and outlier removal as described above). Log_2_ fold-changes relative to WT within the same device were calculated. For each mutant–substrate pair, Welch’s two-sample t-test was used to compare normalized initial rates between mutant and WT. A p-value ≤ 0.05 was considered significant for individual substrate comparisons.

Heatmaps were generated as follows:

● Grey cells: non-significant differences (p > 0.05) with ≥ 2 replicates.
● Colored cells: significant differences (p ≤ 0.05) encoded by log_2_ fold-change.
● Red X: only one replicate for mutant or WT.
● Black X: undetectable initial rates in either mutant or WT.

NSP10–11 was excluded from the main log_2_ fold-change heatmap because no WT activity was detectable; instead, raw initial rates for mutants and WT at NSP10–11 were analyzed separately, and significance was assessed relative to a null hypothesis of zero activity.

### Comparisons to Prior Fitness Measurements

To compare HT-MEK^pro^ biochemical parameters with prior M^pro^ fitness datasets (Fig. S22 and S23), we aligned our variant set to mutations present in (i) Bloom and Neher^40^, which report predicted fitness effects for SARS-CoV-2 proteins, and (ii) Bolon and co-workers^15^, who report deep mutational scanning scores for M^pro^ based on FRET, split transcription factor, and growth readouts.

Variants were matched by residue position and amino acid substitution. For Bloom and Neher, predicted fitness scores for M^pro^ mutations were plotted against our *k*_cat_, *K*_M_, and *k*_cat_/*K*_M_ values, and Spearman rank correlations (ρ) and p-values were computed. For Bolon et al.^15^, we averaged the two reported replicate scores for each metric (or used a single replicate if only one was available). They plotted these against our kinetic parameters for each matching mutation. For each combination of biochemical parameter and cellular readout, Spearman’s ρ and p-values were calculated, along with Pearson’s r and associated p-values.

### PyMOL visualizations and in silico mutagenesis

Structural visualizations were generated using PyMOL (Schrödinger, LLC; open-source executable build; OpenGL 2.1, GLSL 1.20). Structures of SARS-CoV-2 M^pro^ were taken from PDB IDs 7CAM and 7AXM, as indicated in the figure legends. The point mutation S301Y was introduced in silico using PyMOL’s mutagenesis wizard, which selects the highest-probability rotamers from the internal rotamer library. The models were then relaxed (default parameters)^70^, implemented within the Rosetta3 suite. A single representative relaxed pose is shown in Fig. 6c (light blue). Final molecular renderings were ray-traced in PyMOL with ambient occlusion enabled.

### Molecular dynamics simulations and dynamical network analysis

All molecular dynamics (MD) simulations were performed using AMBER22.^71^ Protein residues were described with the ff14SB force field,^72^ and small-molecule components, when present, were parameterized using the GAFF force field.^73^ Structural models of the NSP4-5 substrate–bound SARS-CoV-2 M^pro^ and the ensitrelvir-bound enzyme were obtained from PDB IDs 7T70 and 8HBK, respectively.^74,75^ For each variant, the local regions containing the mutations or the loop-275 replacement were modeled with AlphaFold3,^76^ and the resulting structures were incorporated into the template using AmberTools.^77^

Each protein monomer was capped with an acetyl (ACE) group at the N terminus and an N-methylamide (NME) at the C terminus to neutralize terminal charges. Systems were solvated in a truncated octahedral TIP3P water box with a minimum solute–box distance of 14 Å in all directions.^78^ Sodium and chloride ions were added to neutralize total charge and to achieve a physiological ionic strength of 0.15 M NaCl, calculated based on the initial box volume.

Energy minimization was performed in three sequential stages using the conjugate gradient method: (i) with positional restraints on all atoms, (ii) with restraints on backbone atoms only, and (iii) without restraints. Systems were then gradually heated from 0 to 300 K under the NVT ensemble, followed by equilibration under NPT conditions at 300 K and 1 bar through eight successive steps with gradually reduced restraints. Temperature and pressure were maintained using the Langevin thermostat. Long-range electrostatics were treated with the particle mesh Ewald (PME) method^79^ using a 10 Å cutoff, and all bonds involving hydrogen atoms were constrained with the SHAKE algorithm.^80^ Production MD simulations were run for 100 ns. Trajectories were processed and analyzed using cpptraj in AmberTools.^81^

Dynamical network analysis was performed using the NetworkView plugin in VMD.^82,83^ Networks were constructed by defining each residue Cα atom as a node, and communication between residues was quantified using Cartesian covariance of atomic positional fluctuations. A pair of nodes was connected by an edge if the node distance between the corresponding residues remained within 4.5 Å for at least 75% of the trajectory. Edge weights were assigned proportional to the Cartesian covariance between node positions, such that strongly correlated residues formed high-weight connections. The resulting weighted networks were resolved into communities, defined as groups of residues exhibiting highly correlated motions, using the Girvan–Newman algorithm. The strength of communication between communities was quantified by the total betweenness of all edges that connect the communities.^84^ The networks were visualized using Chimera^85^ and Cytoscape.^86^

### Viral genome assembly and replication assays

All M^pro^ mutations introduced into SARS-CoV-2 replicon plasmids were done as described previously.^87^ SARS-CoV-2 single-round infectious particles were generated as previously described with some modifications.^87^ BHK-21 cells were seeded in a 6-well plates (2×10^5^) and were transfected the next day with 1 μg of pBAC SARS-CoV-2 Spike replicon plasmid, 0.5 μg of Spike Delta variant plasmid, and 5 μg of Nucleocapsid R203M plasmid using Xtremegene 9 DNA transfection reagent (Sigma-Aldrich).^88^ The medium was changed 4-6 hours after transfection and the cells were incubated at 37°C and 5% CO_2_. At 72 hours post-transfection, 2×10^4^ VAT, 293T-AT, or A549-ACE2^h^ cells in 50 μl of culture medium were mixed with 250 μl of 0.45 μm filtered supernatant and plated in a 96-well plate for 6 to 8 hours at 37°C and 5% CO_2_. The cells were washed once with 300 μL culture medium and 100 μl of fresh culture medium was added. The cells were incubated for 24 hours and 50 μl of supernatant was transferred to a white 96-well plate. Promega nanoGlo reagent (50 μl) was added and luminescence was recorded in a Tecan plate reader.

### Split-luciferase cleavage reporter assays

To construct a split-luciferase cleavage reporter vector, the VSV-G coding sequence in pMD2.G (Addgene Plasmid #12259) was replaced with a permuted Firefly luciferase-IRES-Renilla luciferase containing an “optimal” M^pro^ cleavage sequence using HiFi DNA assembly (NEB).^46^ Natural M^pro^ cleavage sequences were introduced into this construct using HiFi DNA assembly (NEB). To construct an M^pro^ doxycycline-inducible expression vector, the Cas9-T2A-EGFP sequence in TCLV2 (Addgene Plasmid #87360) was replaced with NSP5 human codon optimized sequence,^89^ and the puromycin resistance gene replaced with hygromycin resistance gene using HiFi DNA assembly (NEB). NSP5 mutations were introduced into this vector using HiFi DNA assembly (NEB).

To generate 293T cells conditionally expressing NSP5, 293T cells were transfected with 1 μg tet-inducible NSP5 lentiviral vector along with 0.5 μg pMD2.G (Addgene Plasmid #12259) and 1 μg psPAX2 (Addgene Plasmid #12260) in 6-well plates. The media were changed 4-6 hours after transfection. At 48 hours post-transfection, the supernatant was 0.45 μm filtered and stored at -80C. 293T cells in 6-well plate were transduced with 200 μL of the supernatant with 10 μg/mL polybrene (Sigma-Aldrich). After 24 hours, the culture media was changed and 250 μg/mL hygromycin added. After 2 passages, the cells were expanded and frozen.

To conduct cleavage reporter assays across multiple M^pro^ variants and cleavage sites, 1×10^4^ 293T cells conditionally expressing M^pro^ variants were seeded in 96-well plates. The next day, cells were transfected with 100 ng of 12 different split-luciferase cleavage reporter vectors bearing the “optimal” and the 11 natural cleavage sequences using Xtremegene 9 DNA transfection reagent (Sigma-Aldrich). At the time of transfection, 1 μg/mL doxycycline was added. At 48 hours post-transfection, the cells were washed with 200 μL PBS and lysed in 20 μL 1xPLB (Promega). Firefly and Renilla luciferase signals were determined using the Dual-Luciferase Reporter Assay System (Promega) in 5 μL of lysate. For experiments with pelitinib, the inhibitor was added at the time of transfection along with doxycycline. Cell viability was determined using the CellTiter-Glo Assay (Promega).

### Machine learning model generation for biochemical parameter prediction with bootstrapping over increasing training sizes

Random forest, support vector regression, and ridge regression models used as baselines were imported from Sci-Kit Learn. ESM1v per residue embeddings were utilized as input features while output labels were log10 transformed. For robust model performance metrics, we utilized the dataset ablation procedure from Muir et. al 2025.^29^ Our neural net architecture was supervised, used ESM1v embeddings, and had dual-head attention (one focused on per-residue embedding change and one focused on mutant embeddings directly) to form pooled representations that could be used for prediction.

