## Supplemental Materials for "Coupling high-throughput protease enzymology with viral replication reveals biochemical constraints of viral fitness"

The PDF file includes:
Figs. S1 to S35
Tables S1 to S10

### Supplementary Figures:

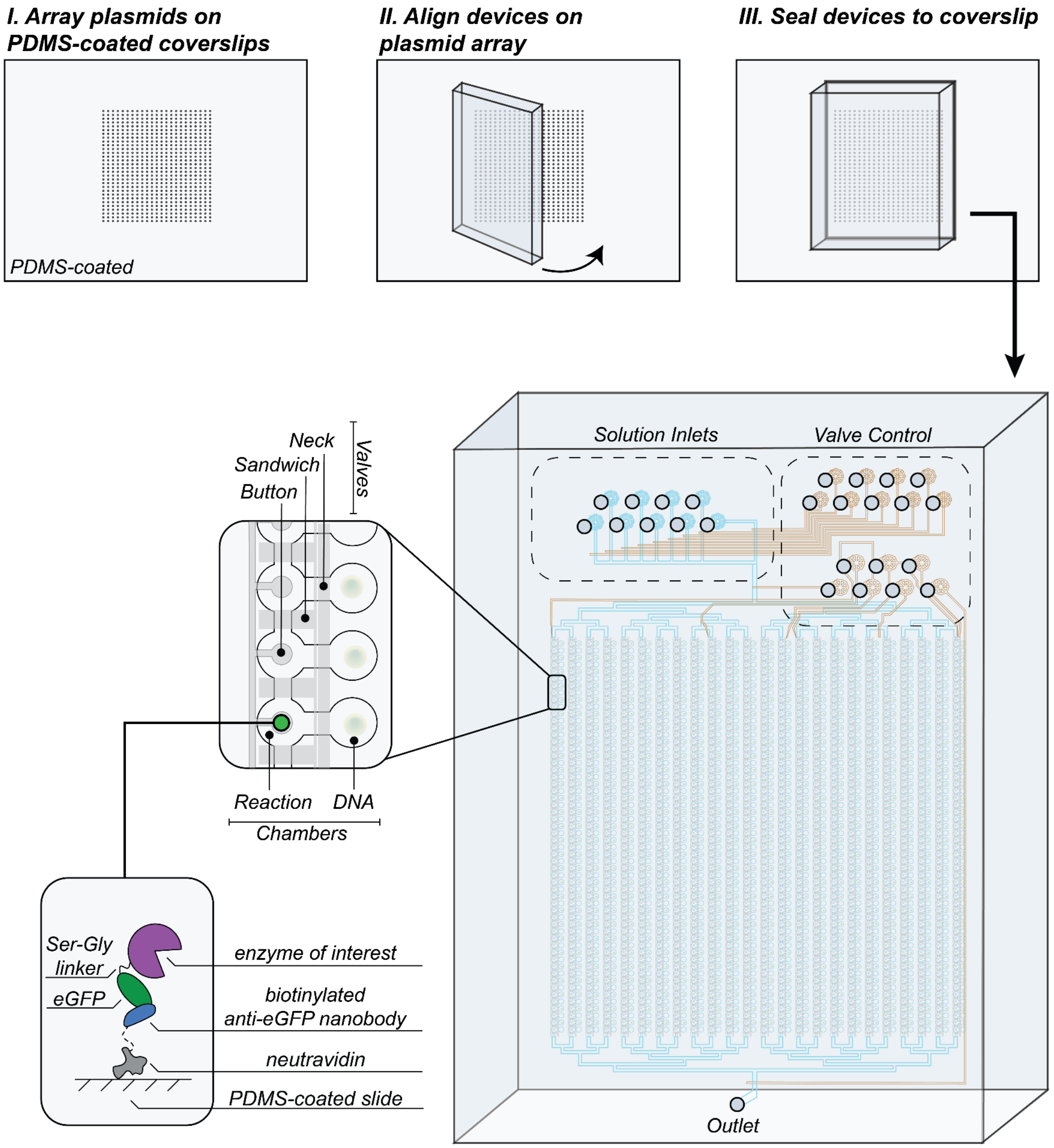


**Figure S1. Device setup, alignment, and architecture.**

The plasmid arraying and slide-mounted device assembly are shown schematically. An enlarged view highlights reagent inlets for introducing substrates and other solutions, along with pneumatic control lines that actuate the Neck, Sandwich, and Button valves (inset). A representative reaction chamber illustrates the “pedestal” region and capture scheme used to immobilize the enzyme of interest. More details on chamber and valve architecture can be found in Figure S2.

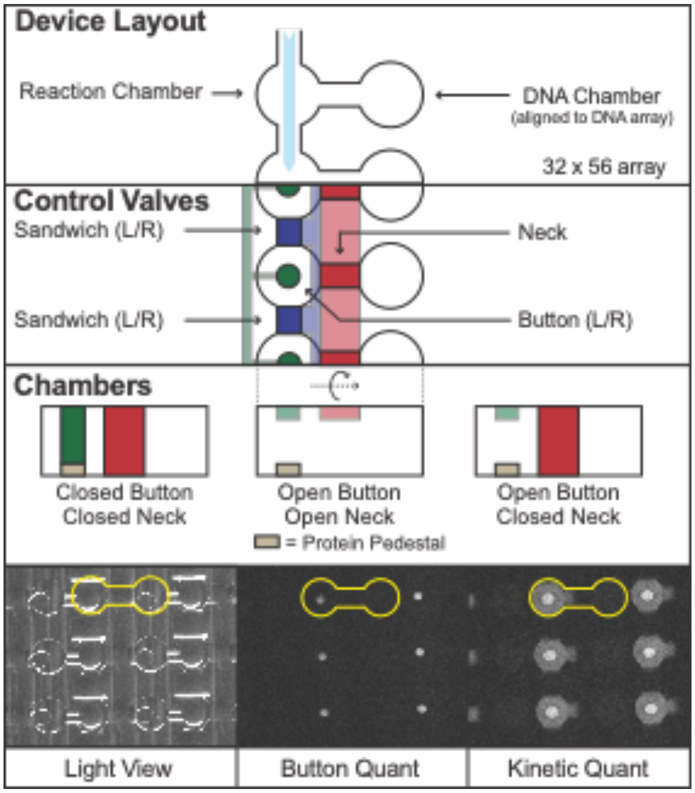


**Figure S2. Overview of HT-MEK device architecture, layout, and visualization.**
The layout of the DNA and reaction chambers is shown, with the direction of flow indicated by a light blue arrow. Sandwich, Button, and Neck valves protect the protein pedestal and isolate each DNA and reaction chamber. Representative microscope images are shown (left to right): device features under room light, the eGFP channel visualizing $M^{pro}$ variant "buttons," and the substrate channel reporting substrate cleavage activity in the reaction chambers.

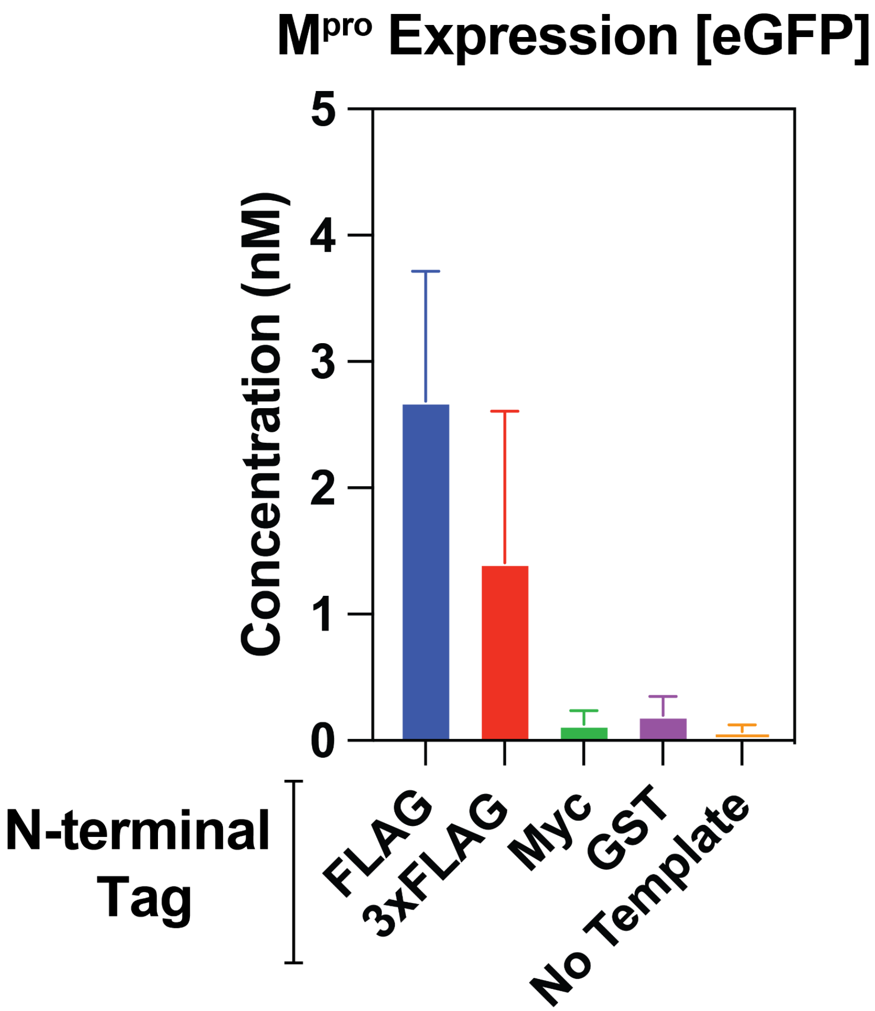


**Figure S3. On-chip expression of SARS-CoV-2 M$^{pro}$ with different N-terminal tags.**
All templates were concentration normalized with the exception of the no template control.

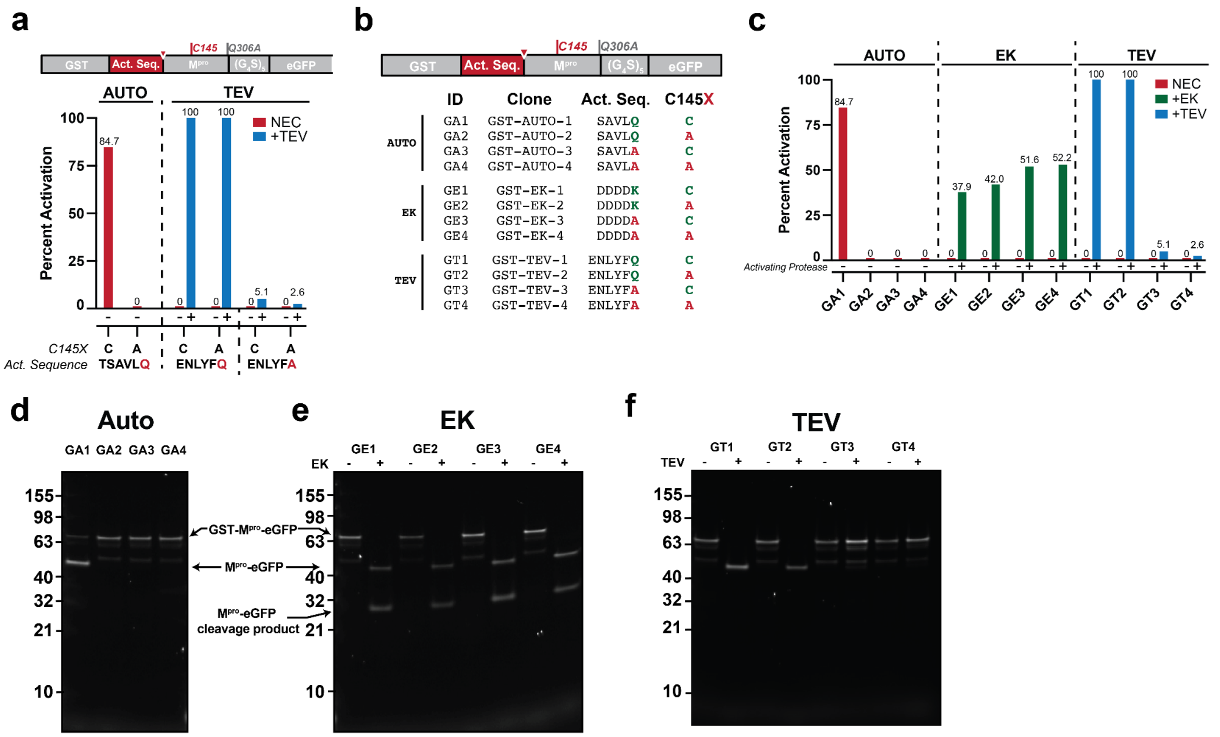


**Figure S4. Off-device activation of M$^{pro}$-eGFP using auto-activation, enterokinase, and TEV protease.**

**(a)** Quantification of activation of WT M$^{pro}$ and catalytically-dead cysteine control mutant (C145A) from in gel fluorescence. Activation sequences are: auto-activation (SAVLQ|SGFRK), TEV activation (ENLYFQ|SGFRK), and TEV control (ENLYFA|SGFRK). **(b)** Table of M$^{pro}$-egfp clones used as activation controls. Each of the four activation strategies contained two WT M$^{pro}$ and two M$^{pro}$ C145A (catalytically inactive). Additionally, the cleavage site of the activating protease was knocked out with an alanine. **(c)** Quantification of in gel fluorescence for each of the three activation strategies. Gels are shown in d-f. **(d-f).** Gel fluorescence images used for each construct comparing inactive and active forms for each of the M$^{pro}$-eGFP constructs.

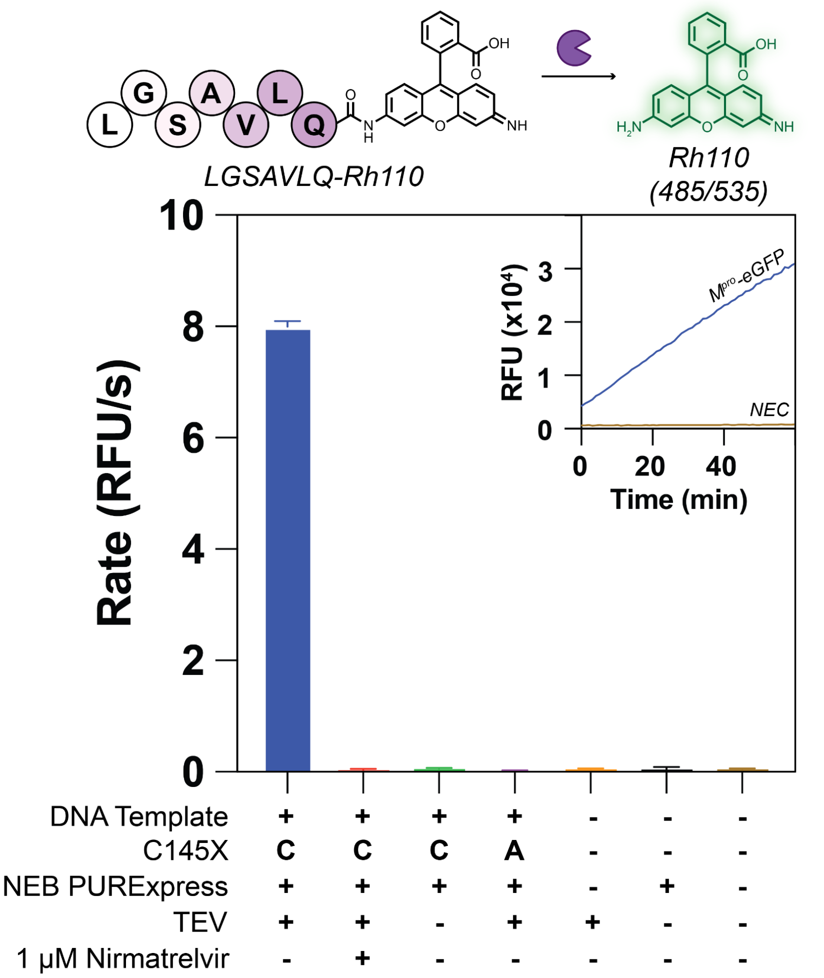


**Figure S5. TEV-activated WT M$^{pro}$ displays robust activity on a commercial fluorogenic substrate.**

Reaction scheme of Rhodamine-110-labelled fluorogenic substrate (top). Proteolytic activity of WT Mpro under different conditions quantified off-chip on a plate reader. Inset contains representative progress curves for activated WT Mpro and no enzyme controls (NEC).

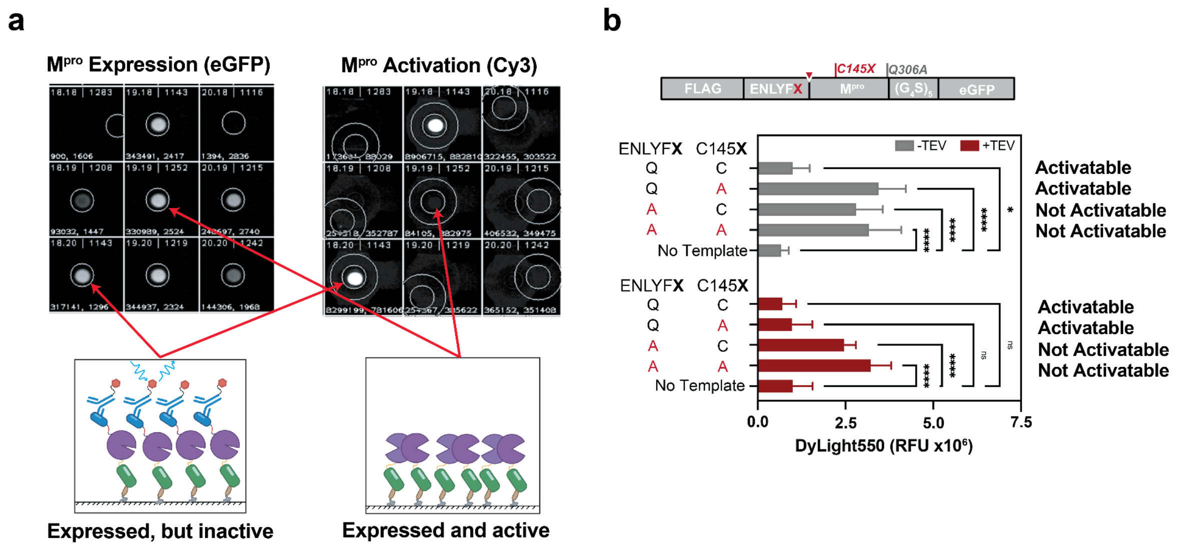


**Figure S6. Efficient on-chip activation of FLAG-TEV-M$^{pro}$ constructs.**
**(a)** Representative images and cartoon representation of activation visualized by binding of N-terminal tag with dye-conjugated antibody (DyLight550). **(b)** Activation of M$^{pro}$-eGFP controls on device visualized by anti-FLAG antibody conjugated to DyLight550.

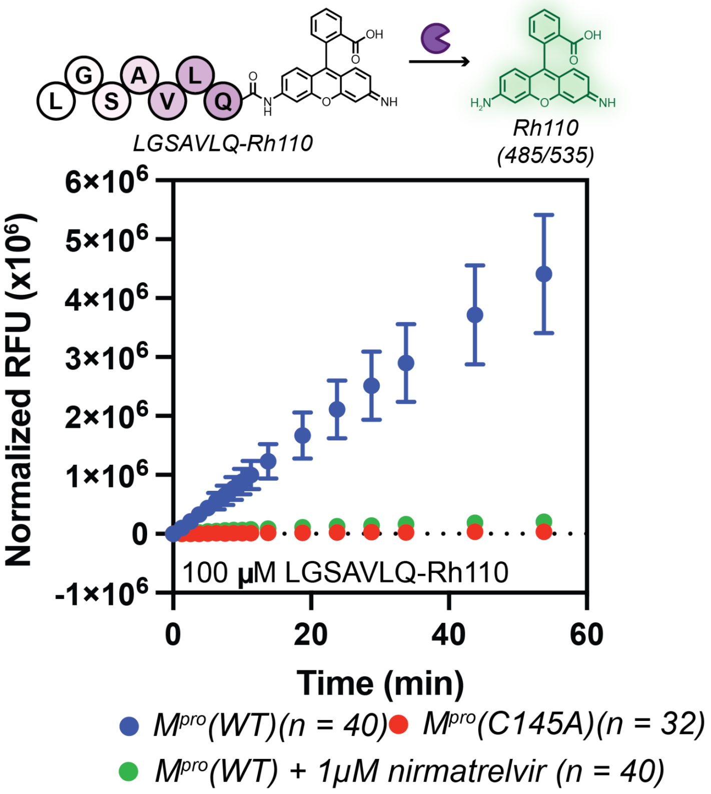


**Figure S7. Quantification of on-chip M$^{pro}$ activity for the optimized M$^{pro}$-eGFP construct with N-terminal FLAG-tag and TEV cleavage site.**

Example Rh110-labelled fluorogenic progress curve for chambers containing WT M$^{pro}$ enzyme and active site control (C145A) with and without nirmatrelvir (1µM).

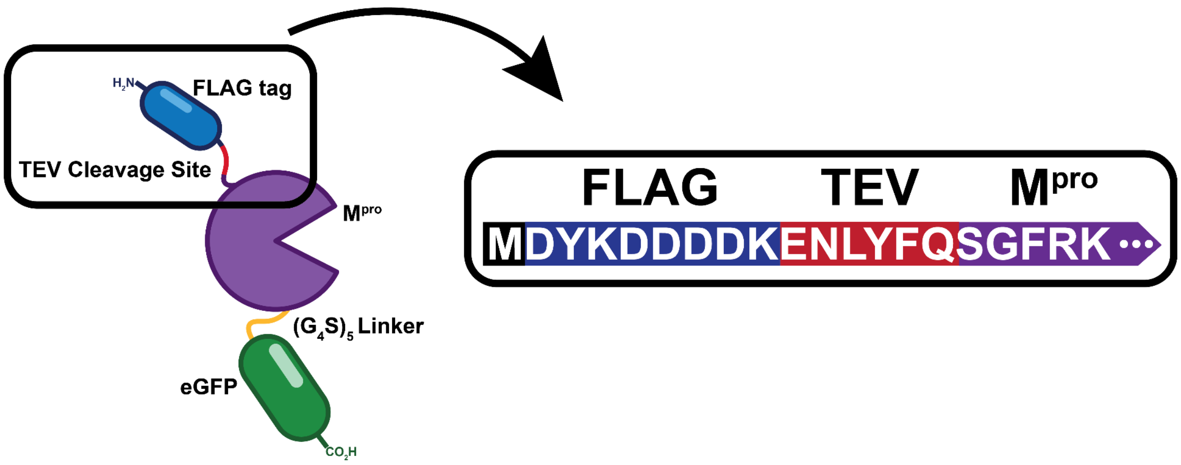


**Figure S8. Optimized M$^{pro}$-eGFP construct with N-terminal FLAG-tag and TEV cleavage site.**

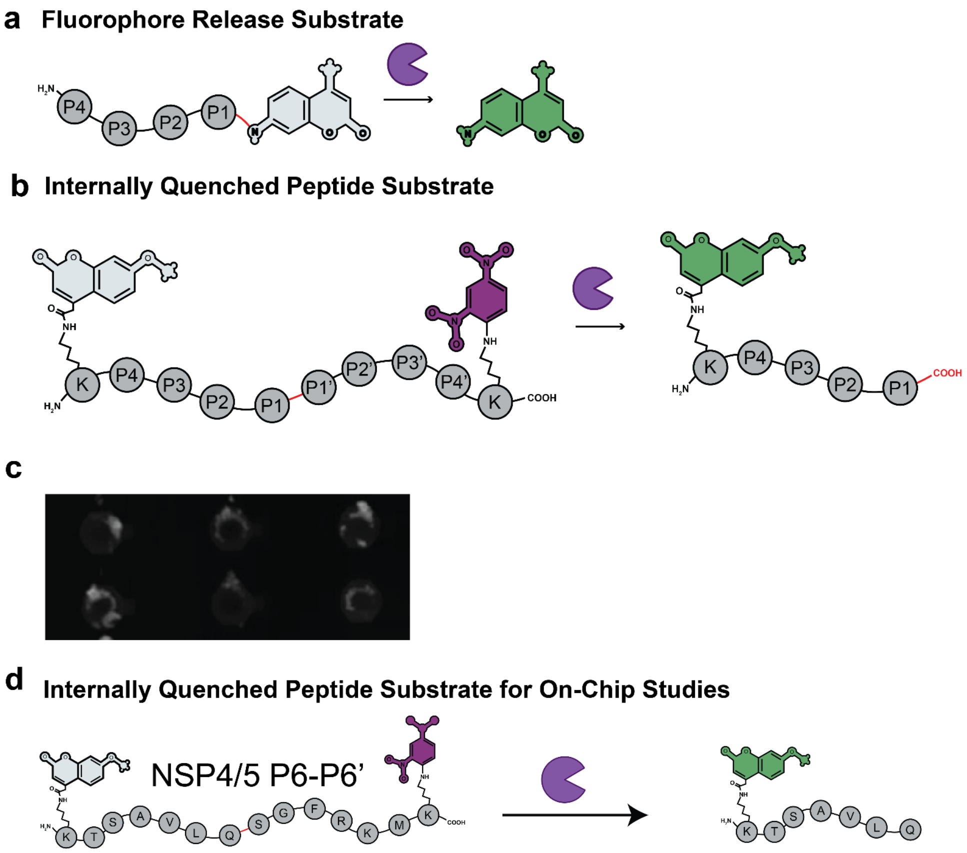


**Figure S9. Protease substrate design strategies.**
**(a)** Fluorophore-release substrates generate signal only upon cleavage of the P1–dye bond, ensuring specificity, but typically exhibit weaker $K_M$ values requiring higher substrate concentrations and incorporate a synthetic, non-natural P1′ residue. **(b)** Internally quenched peptide substrates provide tighter $K_M$ values, allow inclusion of natural P1′–P4′ residues and cleavage natural cleavage sequences, and report directly on bond scission, but can also generate signal from off-site cleavage events. **(c)** Example on-chip image of an insolubale, presumably aggregated internally quenched peptide substrate based on the P5-P5' of the nsp7/8 cleavage site. **(d)** Internally quanched substrate used for on-chip studies based on P6-P6' of the nsp4/5 cleavage site flanked by a 5-carboxyfluorescein (5-FAM) donor and a 2,4-dinitrophenyl (Dnp) quencher pair. 5-FAM was chosen because of its high solubility and negative charge.

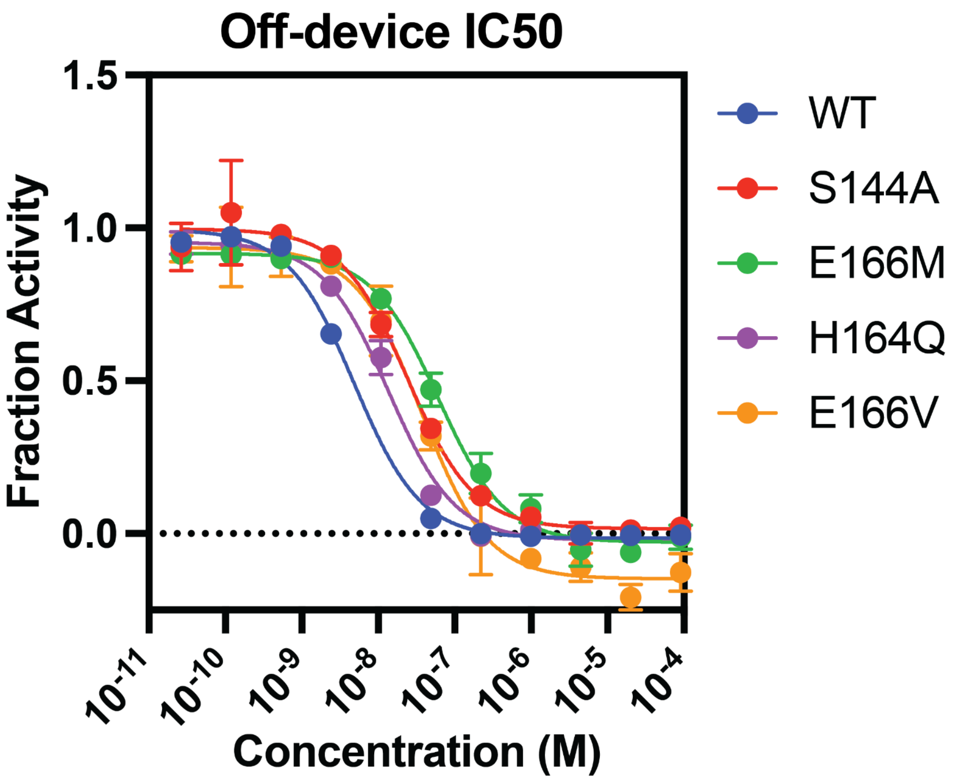


**Figure S10. Inhibition curves determined in a plate reader (off-chip) for select SARS-CoV-2 variants with ensitrelvir.**
$IC_{50}$s from fits to these curves are compared to on-chip values in Fig. 2g.

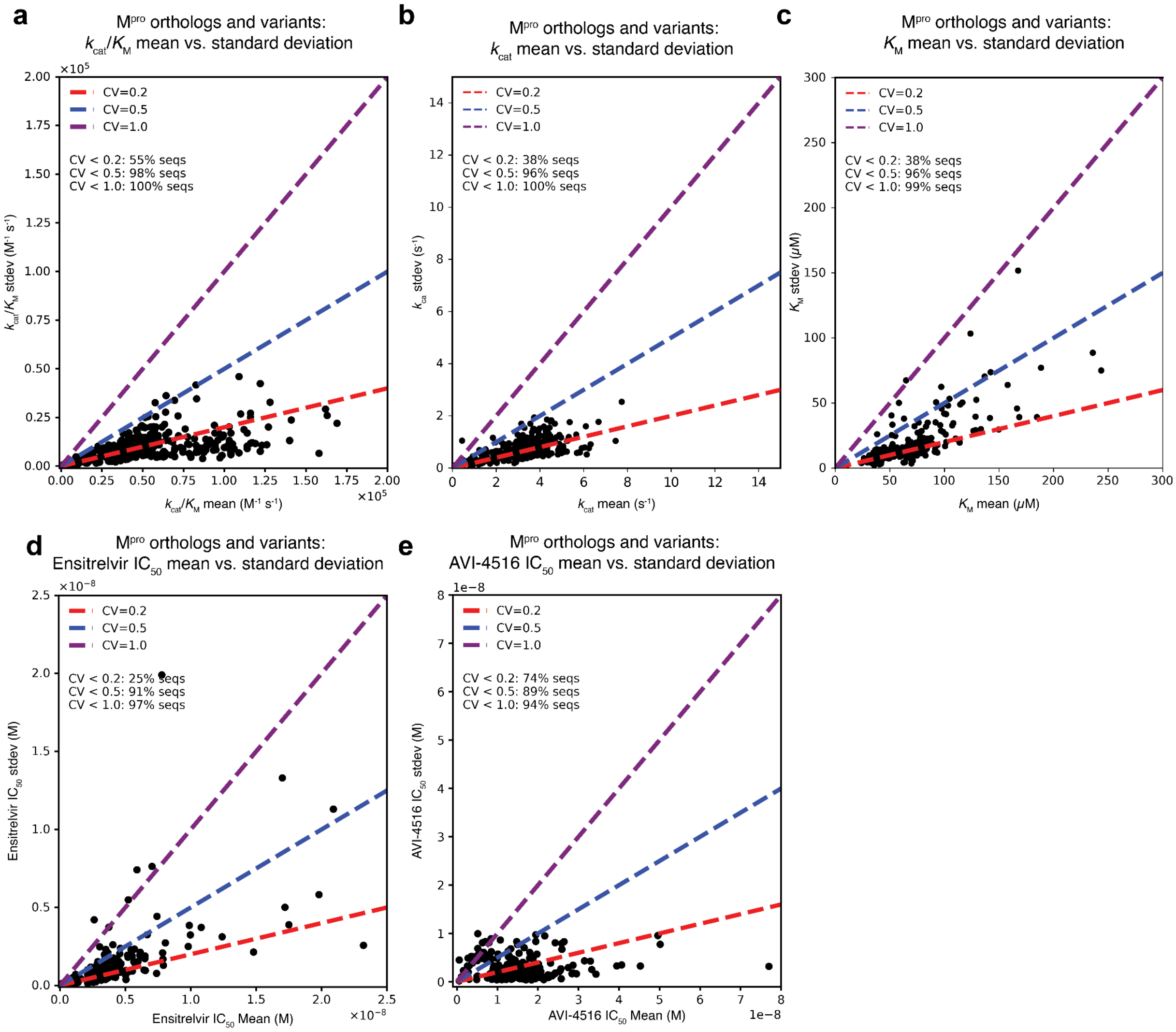


**Figure S11. Coefficient of variation for on-chip M$^{pro}$ catalysis and inhibition measurements.** The coefficient of variation (CV = standard deviation/mean) is plotted for **(a)** ortholog and SARS-CoV-2 M$^{pro}$ variant $k_{cat}/K_M$, (b) $k_{cat}$, (c) $K_M$, (d) ensitrelvir $IC_{50}$, and (e) AVI-4516 $IC_{50}$. Dashed lines representing a CV of 0.2, 0.5, and 1.0 are plotted (red, blue, and purple respectively). The percentage of sequences with CV less than these thresholds for eachplot is indicated by text under the legends.

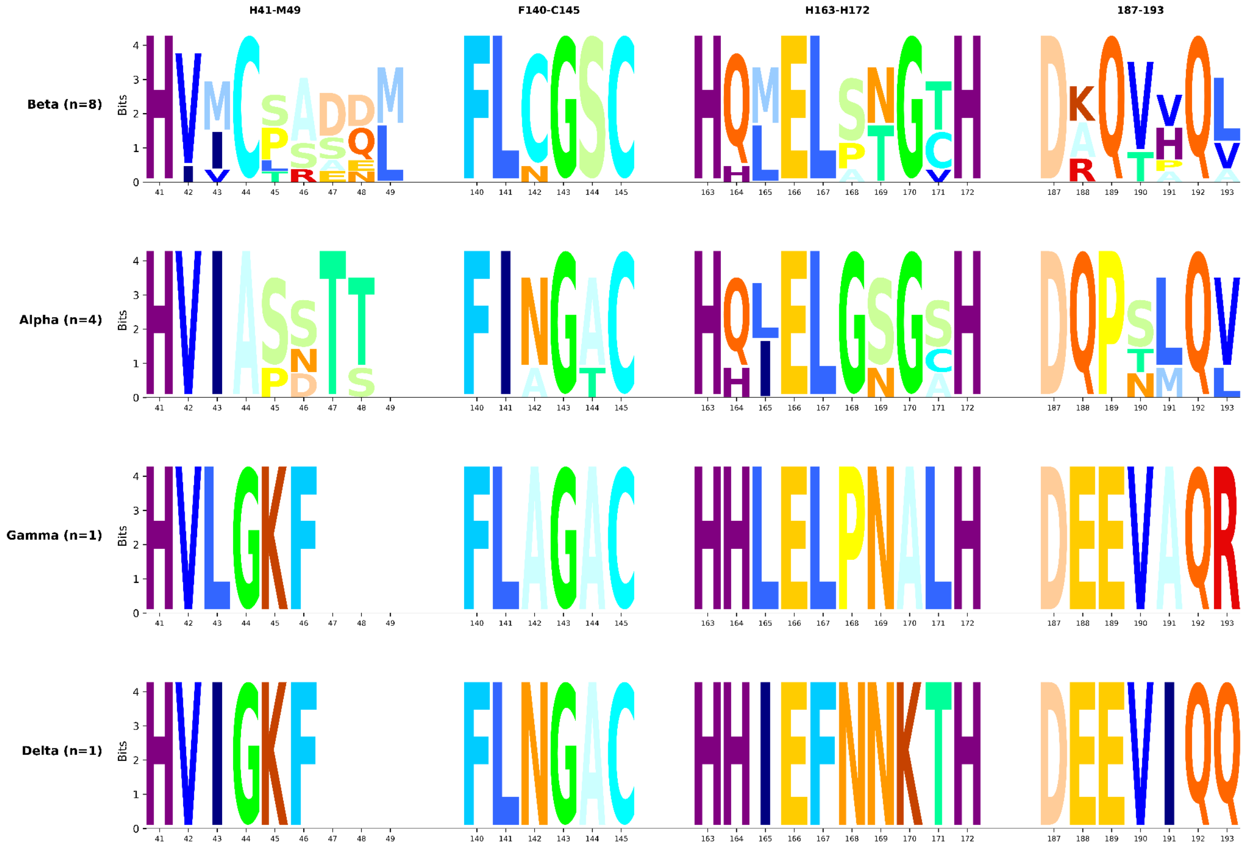


**Figure S12. Active site residue logos of M$^{pro}$ orthologs in Fig 1f.**
Active site residue logos were generated for each genera for the fourteen orthologs tested. All orthologs were aligned to the SARS-CoV-2 reference. Orthologous M$^{pro}$ sequences can be found in Table S2.

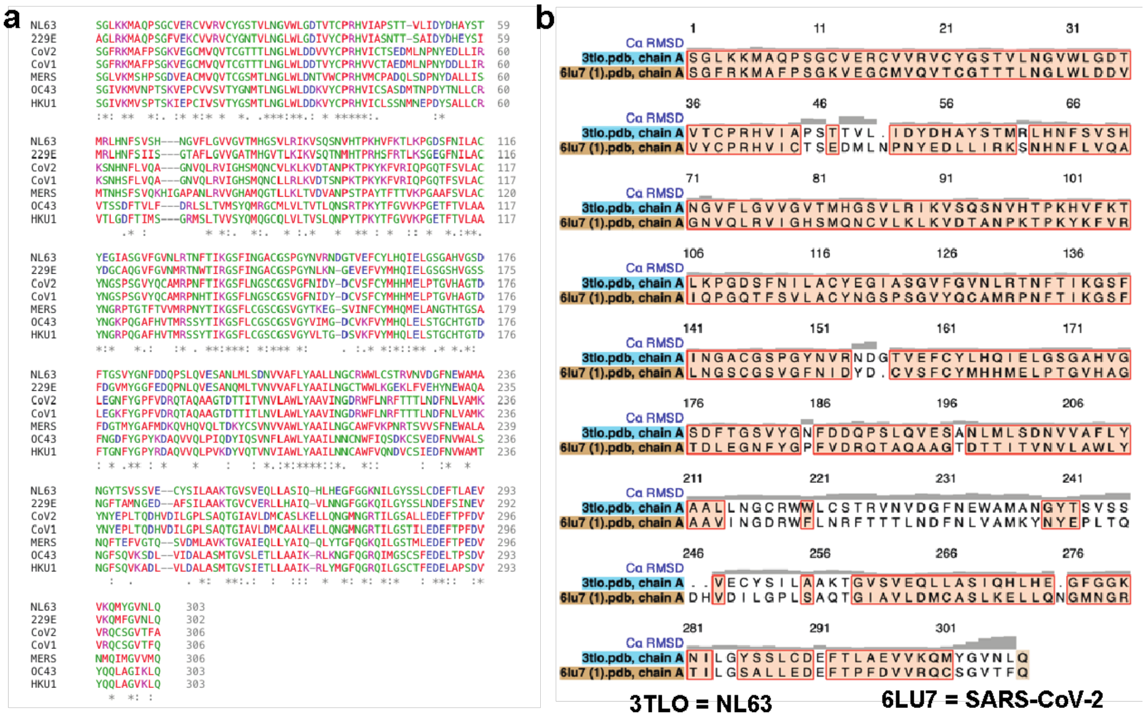


**Figure S13. Sequence and Structural alignment between SARS-CoV-2 M$^{pro}$ and other orthologs.**

**(a)** Sequence alignment of SARS-CoV-2 M$^{pro}$ and other orthologs measured herein. **(b)** Structural alignment of SARS-CoV-2 M$^{pro}$ and NL63.

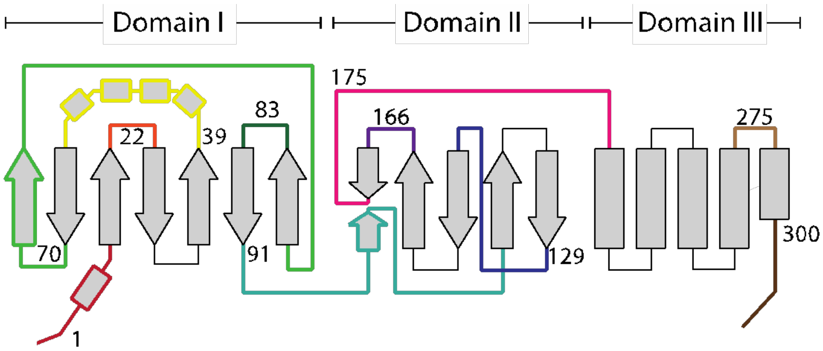


| Region | Sites | CoV-2 | NL63 |
|---|---|---|---|
| 1 | 1-16 | SGFRKMAFPSGKVEGC | SGLKKMAQPSGCVERC |
| 22 | 22-25 | CGTT | YGST |
| 39 | 39-66 | PRHVICTSEDMLNPNYEDLLIRKSNHNF | PRHVIAPSTTVLdelIDYDHAYSTMRLHNF |
| 70 | 70-77 | AGNVQLRV | IINGVFLGV |
| 83 | 83-86 | QNCV | HGSV |
| 91 | 91-11 | VDTANPKTPKYKFVRIQPGQT | VSQSNVHTPKHVFKTLKPGDS |
| 129 | 129-148 | AMRPNFTIKGSFLNGSCGSV | NLRTNFTIKGSFINGACGSP |
| 166 | 166-172 | ELPTGVH | ELGSGAH |
| 175 | 175-200 | TDLEGNFYGPFVDRQTAQAAGTDTTI | SDFTGSVYGNFDDQPSLQVESANLML |
| 275 | 275-292 | GMNGRTILGSALLEDEFT | GFGGKNILGYSSLCDEFT |
| 300 | 300-306 | CSGVTFA | MYGVNLA |

**Figure S14. Selected loops from NL63 $M^{pro}$ swaped into SARS-CoV-2 $M^{pro}$**

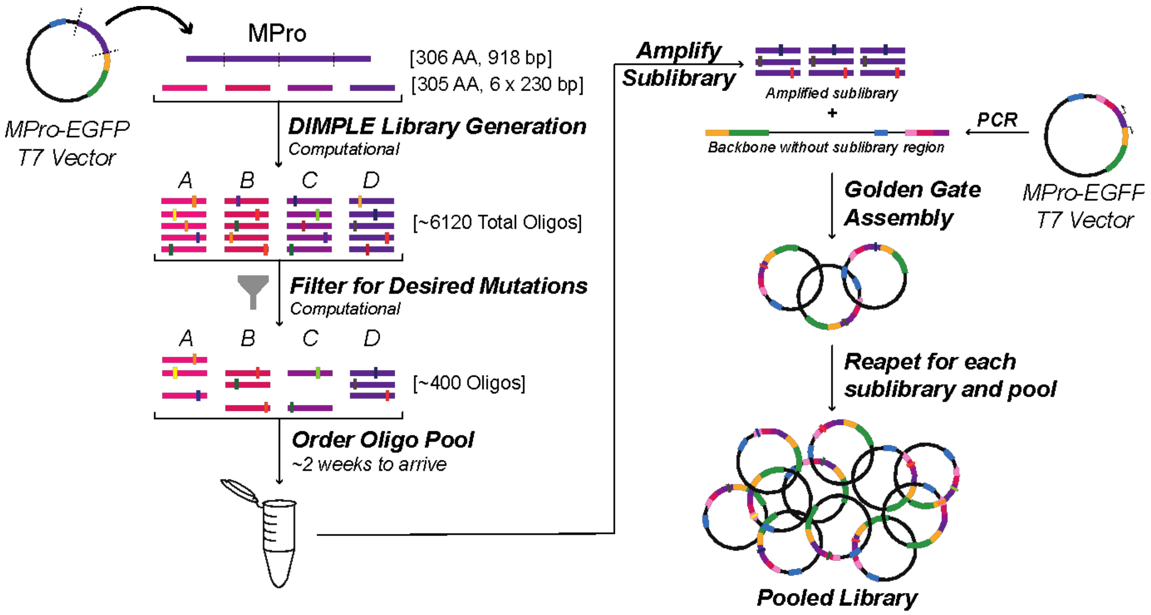


**Figure S15. Cloning strategy for pooled M$^{pro}$ variant libraries.**

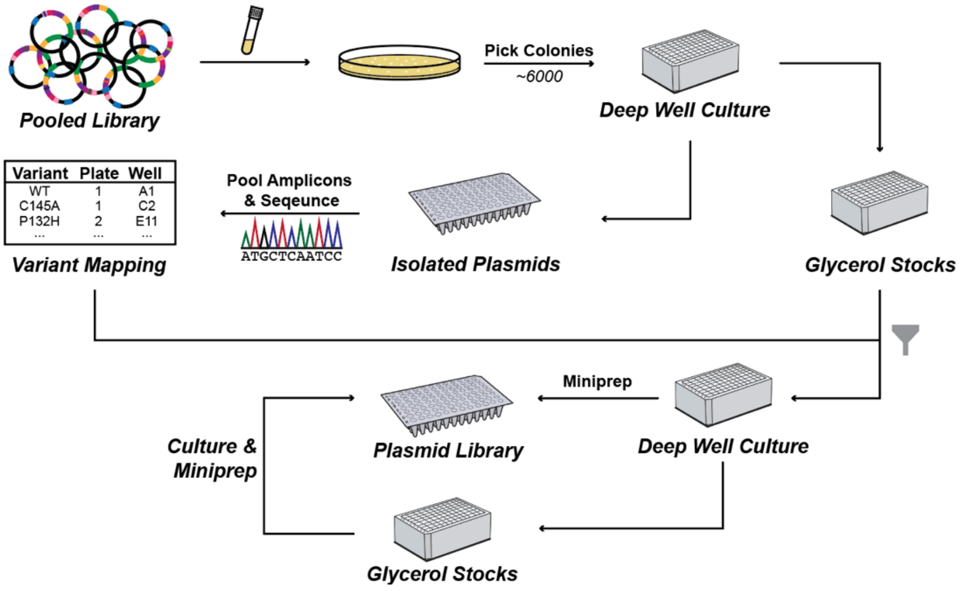


**Figure S16. Pipeline to the isolation and sequence identification of single plasmids from pooled M$^{pro}$ variant libraries.**

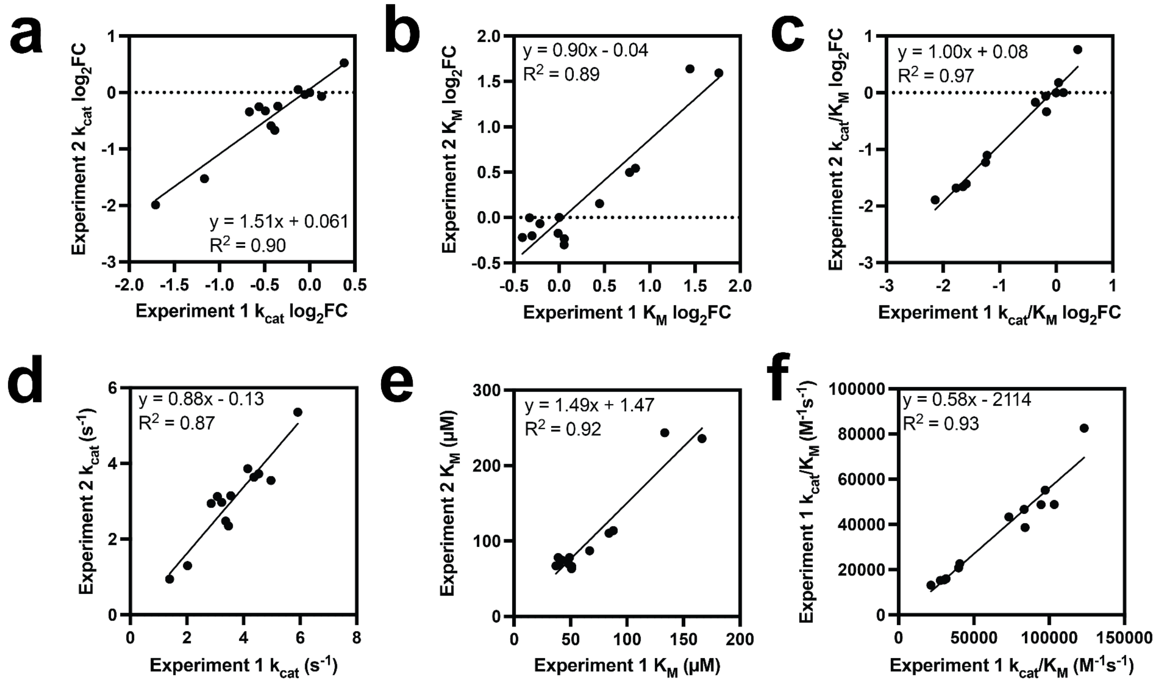


**Figure S17. Comparison between log2FC values and numeric values for $k_{cat}$, $K_M$, and $k_{cat}/K_M$ for WT and twelve variant $M^{pro}$ proteases.**
The common variants between the two experiments were: A173V, H164Q, K088R, L089I, L141I, L205V, M165I, P132T, Q189P, S144A, T169S, and Y126F.

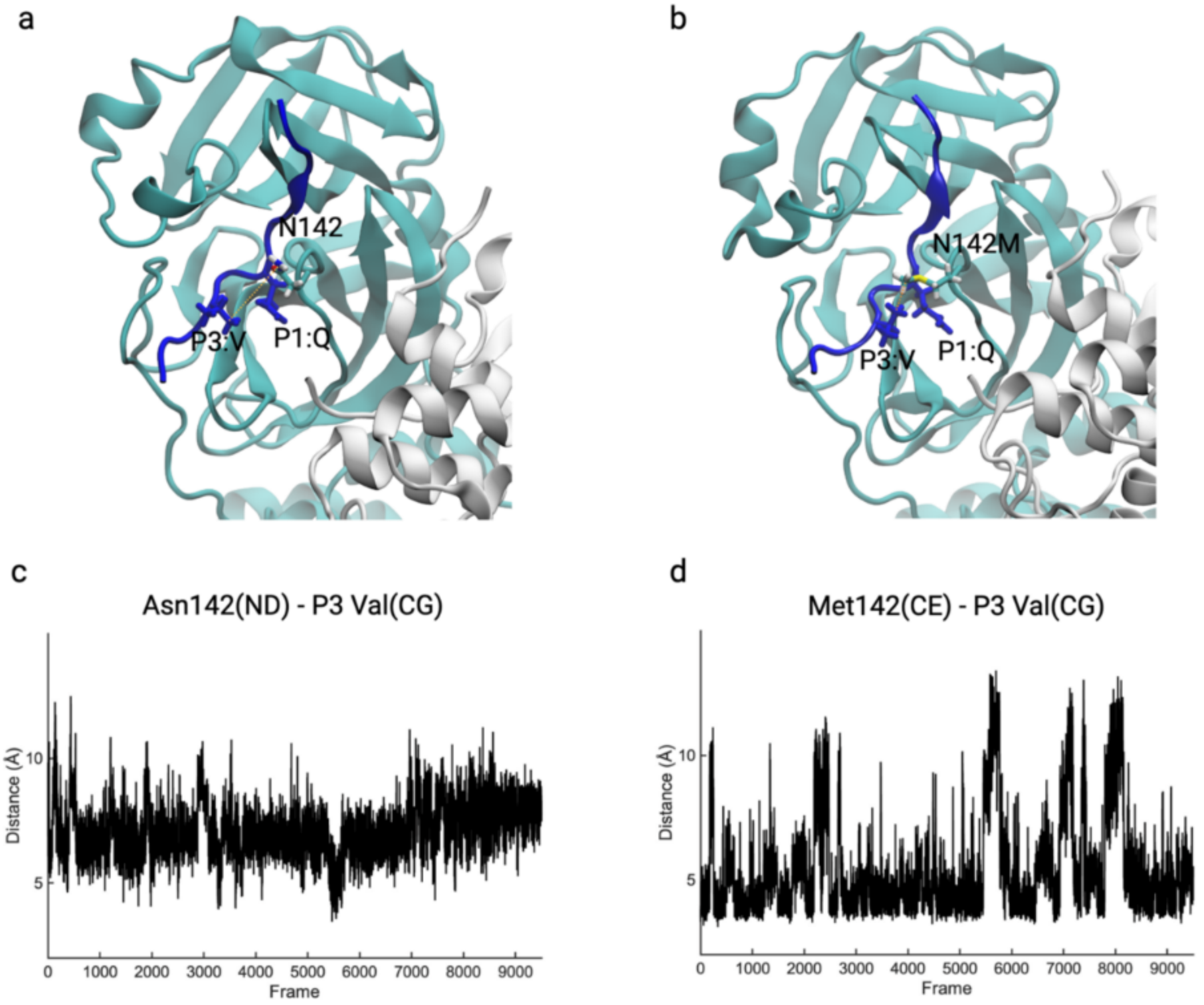


**Figure S18. MD simulations reveal enhanced contacts between residue 142 and the P3 Val at the NSP4-5 cleavage site in N142M SARS-CoV-2 $M^{pro}$.**

**(a, b)** Representative MD snapshots for **(a)** wild type (Asn142) and **(b)** N142M (Met142) $M^{pro}$ bound to the NSP4-5 substrate, illustrating closer packing of Met142 against the P3 Val side chain in NSP4-5. **(c, d)** Distance measurements over 100-ns MD trajectories between **(c)** Asn142 (ND)-P3 Val (CG) and **(d)** Met142(CE)-P3 Val(CG). The N142M mutant exhibits consistently short Met142-P3 Val distances across the majority of simulation frames.

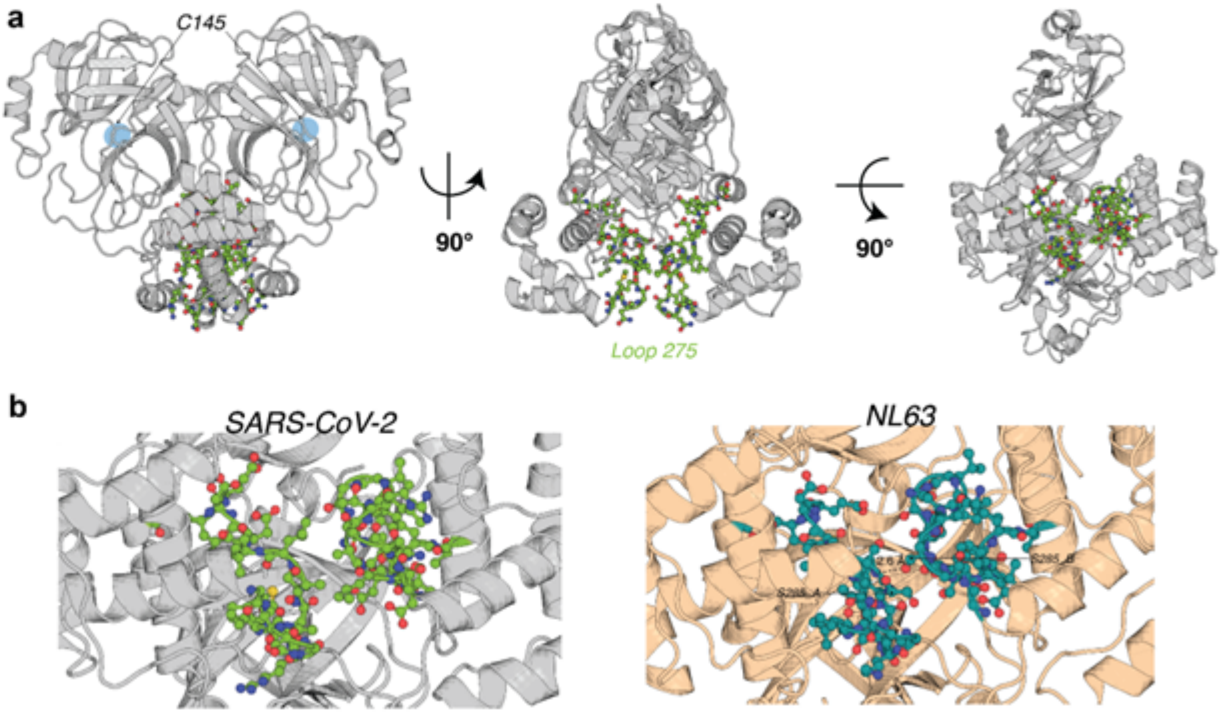


**Figure S19. Loop 275–292 (“Loop 275”) in SARS-CoV-2 M$^{pro}$ (PDB 7CAM) and NL63 Mpro (PDB 5GWY).**

**(a)** Views from multiple orientations showing Loop 275 (green) relative to the catalytic Cys145 (C145). **(b)** Close-up of the dimer interface highlighting Loop 275 contacts. In NL63 Mpro, Ser285 forms a backbone hydrogen bond to the partner protomer (dashed line); the corresponding interaction is absent in SARS-CoV-2 M$^{pro}$.

### NL63 Loop 275 Modifications

CoV2:GMNGRTILGSALLEDEFT

NL63:GFGGKNILGYSSLCDEFT

| | kcat | | KM | | kcat/KM | |
|---|---|---|---|---|---|---|
| Mutation | log2FC | log10(p-value) | log2FC | log10(p-value) | log2FC | log10(p-value) |
| SARS2-NL63-Loop275 | 0.03 | 0.80 | -0.88 | 3.16 | 0.89 | 4.48 |
| SARS2-M276F | -0.05 | 0.46 | -0.20 | 1.03 | 0.17 | 0.86 |
| SARS2-N277G | -0.14 | 1.13 | -0.20 | 0.93 | 0.20 | 0.39 |
| SARS2-R279K | -0.11 | 0.56 | -0.20 | 0.88 | 0.07 | 0.47 |
| SARS2-T280N | -0.47 | 1.29 | -0.08 | 0.39 | -0.40 | 1.66 |
| SARS2-S284Y | -0.73 | 2.16 | -0.04 | 0.46 | -0.74 | 3.15 |
| SARS2-A285S | -0.39 | 1.14 | 0.06 | 0.70 | -0.48 | 1.37 |
| SARS2-L286S | 0.22 | 0.84 | 0.03 | 0.37 | 0.21 | 0.79 |
| SARS2-E288C | -2.30 | 4.81 | -0.03 | 0.36 | -2.14 | 4.23 |

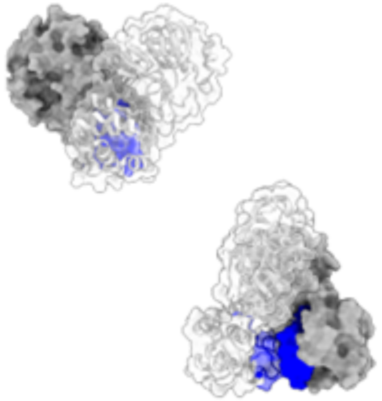

**Figure S20. Effects of individual residue swaps within loop 275–292 between NL63 and SARS-CoV-2 M$^{pro}$ on $k_{cat}$, $K_M$, and $k_{cat}/K_M$.**

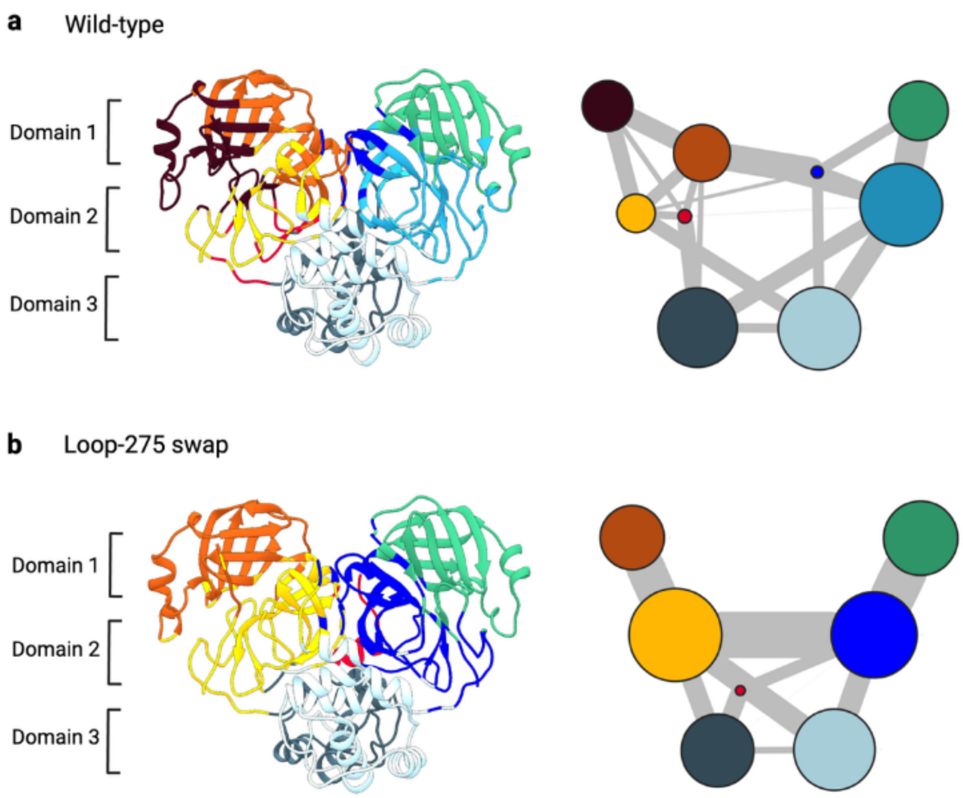


**Figure S21. Community network analysis of –SARS-CoV-2 $M^{pro}$ wild-type and loop-275 swap.** (a) Community analysis of 100-ns MD trajectories of the wild-type $M^{pro}$. Each community within the $M^{pro}$ dimer is uniquely colored and mapped onto the corresponding structural regions. Line thickness between communities reflects the strength of dynamic communication (edge betweenness) during the simulation (See methods). (b) Community analysis of 100-ns MD trajectories of the Loop -275 swap (NL63 residues 275–292 inserted in domain III at the dimer interface). The Loop-275 swap caused several communities in domain I and domain II to merge into larger communities (orange, yellow, green and blue). These results suggest that the loop-swap increases communication across the dimer interface, resulting in enhanced inter-protomer coupling and more coordinated motions between the two protomers compared to the wild-type $M^{pro}$.

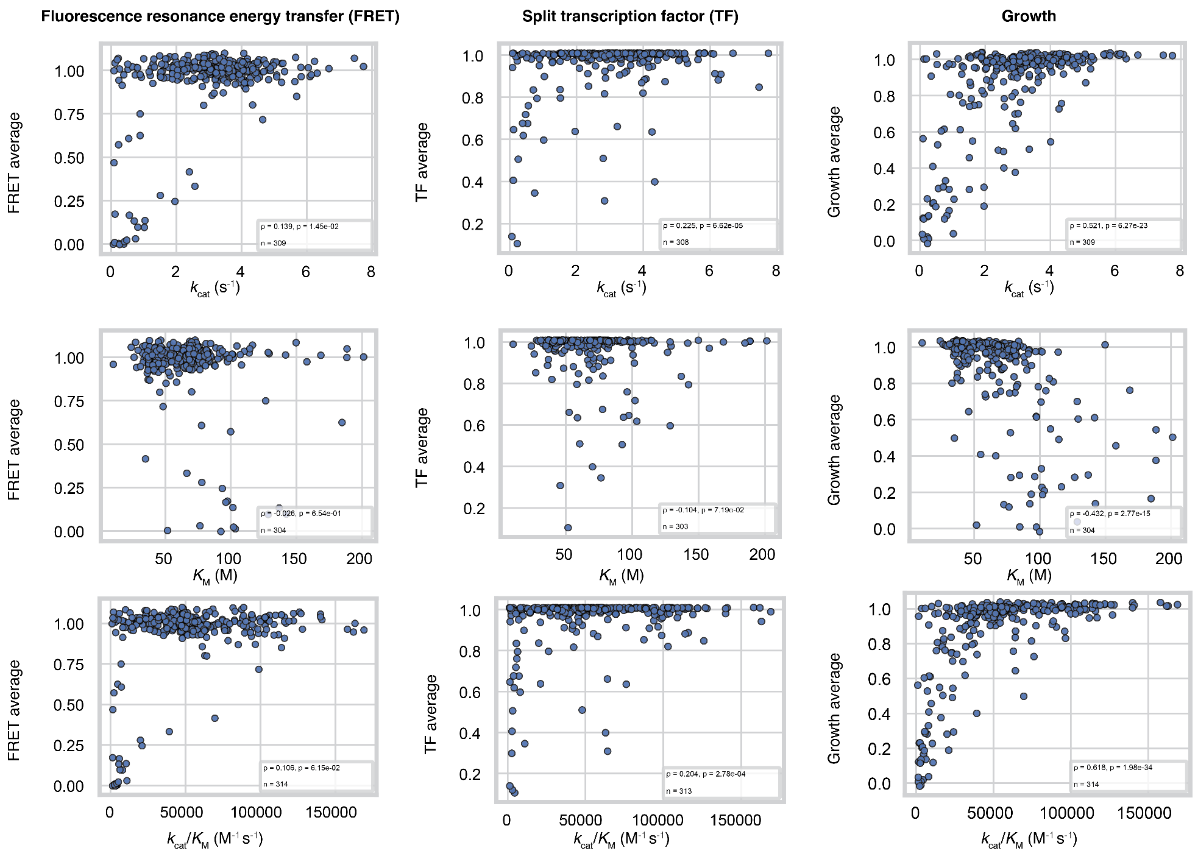


**Figure S22. Comparisons of HT-MEK$^{pro}$ catalytic measurements with yeast-based high throughput screening.**

Scatterplots comparing catalytic turnover ($k_{cat}$), Michaelis constant ($K_M$), and catalytic efficiency ($k_{cat}/K_M$) measured by HT-MEK$^{pro}$ with functional scores from yeast-based deep mutational scanning (DMS) assays. Each point represents a single M$^{pro}$ variant for which both biochemical and DMS measurements were available. Panels correspond to the three independent DMS readouts: intracellular FRET-based NSP4–5 cleavage reporter (left), split transcription factor reporter (middle), and growth (right). For each comparison, Spearman's rank correlation coefficient (ρ) and associated p-value are reported in the panel.

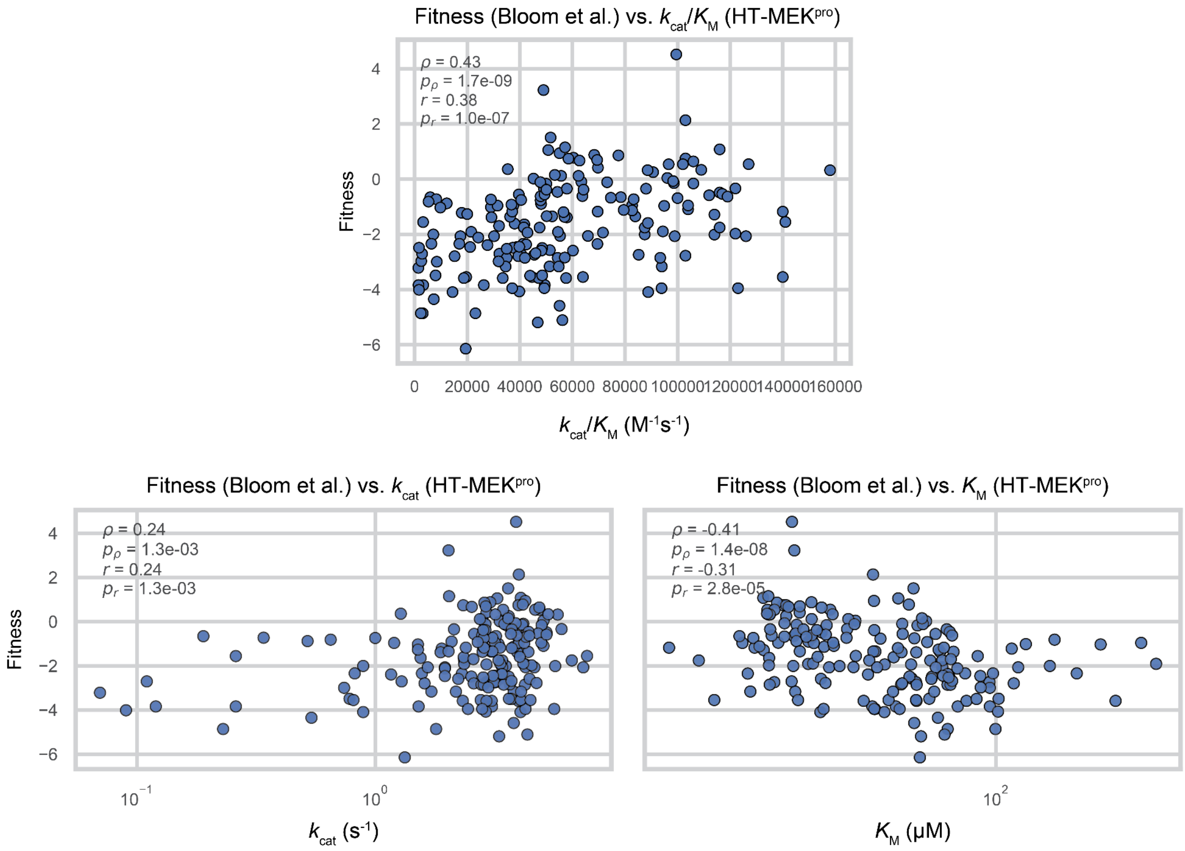


**Figure S23. Comparisons of HT-MEK$^{pro}$ catalytic measurements with fitness effects estimated from SRAs-CoV-2 genome mutations.**

Scatterplot comparing catalytic turnover ($k_{cat}$; bottom left), Michaelis constant ($K_M$; bottom right), and catalytic efficiency ($k_{cat}/K_M$; top middle) measured by HT-MEK$^{pro}$ with fitness effects inferred from coronavirus phylogenetic analyses (Bloom et al.). Each point represents a single M$^{pro}$ variant for which both biochemical measurements and phylogeny-based fitness estimates were available. Spearman's rank correlation coefficient (ρ) and associated p-value (p) are reported in each panel.

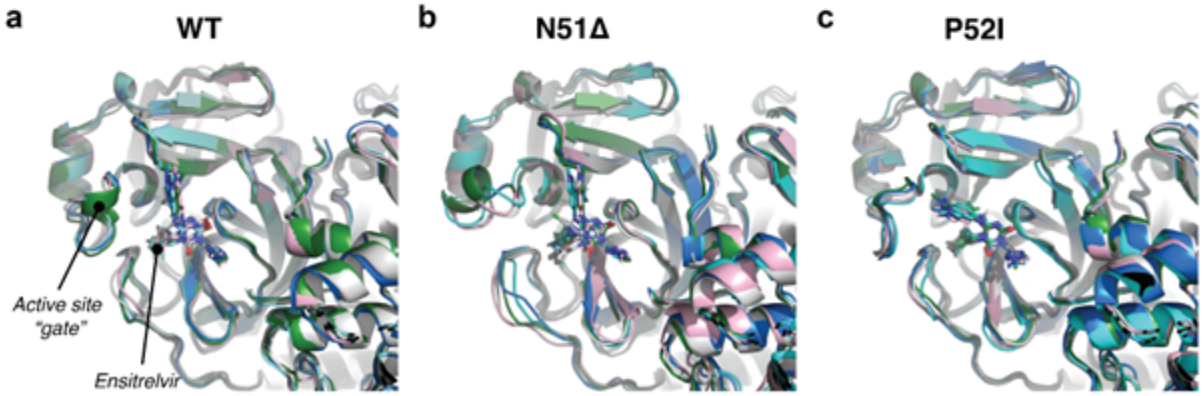


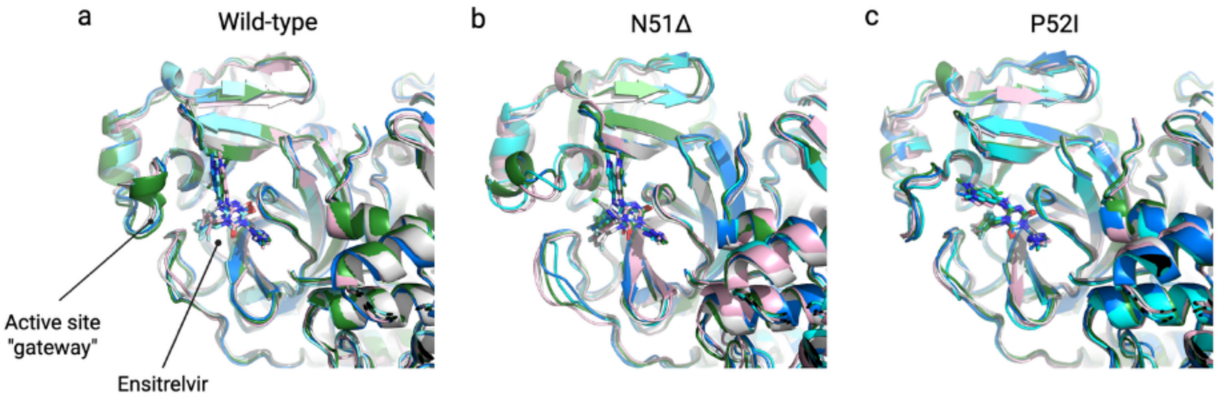


**Figure S24. MD simulations of ensitrelvir-resistant M$^{pro}$ mutants at the active site "gateway".**
**(a–c)** Representative conformations of **(a)** wild-type Mpro, **(b)** N51Δ, and **(c)** P52I highlighting the 45–53 "gateway" region. In the wild type, this loop adopts a well-defined conformation that partially occludes access to the active-site cleft. In N51Δ, the loop exhibits increased flexibility, as reflected by greater conformational heterogeneity. In P52I, the loop preferentially samples a more open state, widening the gateway and increasing accessibility to the active site.

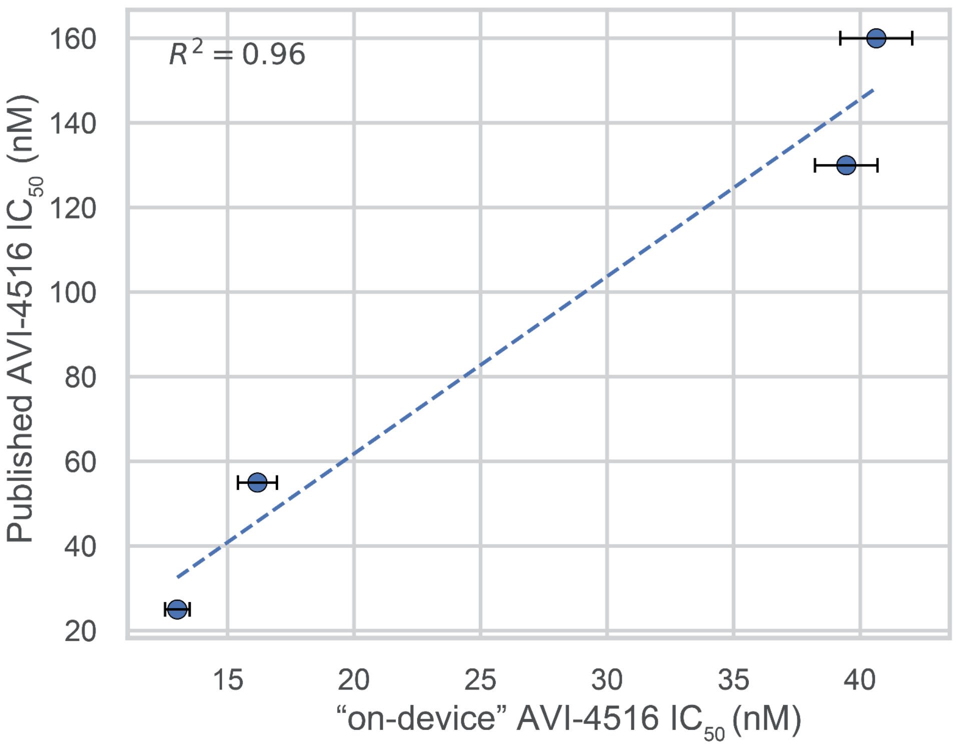


**Figure S25. Benchmarking HT-MEK$^{pro}$ $IC_{50}$ measurements for AVI-4516.**
Scatter plot comparing AVI-4516 $IC_{50}$ values measured on-device by HT-MEK$^{pro}$ with previously reported off-device measurements for four SARS-CoV-2 M$^{pro}$ variants by Detomasi et. al. (2025). Each point represents a single variant; the solid line indicates the best-fit linear regression ($R^2$ = 0.96), demonstrating close agreement between on-device and published $IC_{50}$ values.

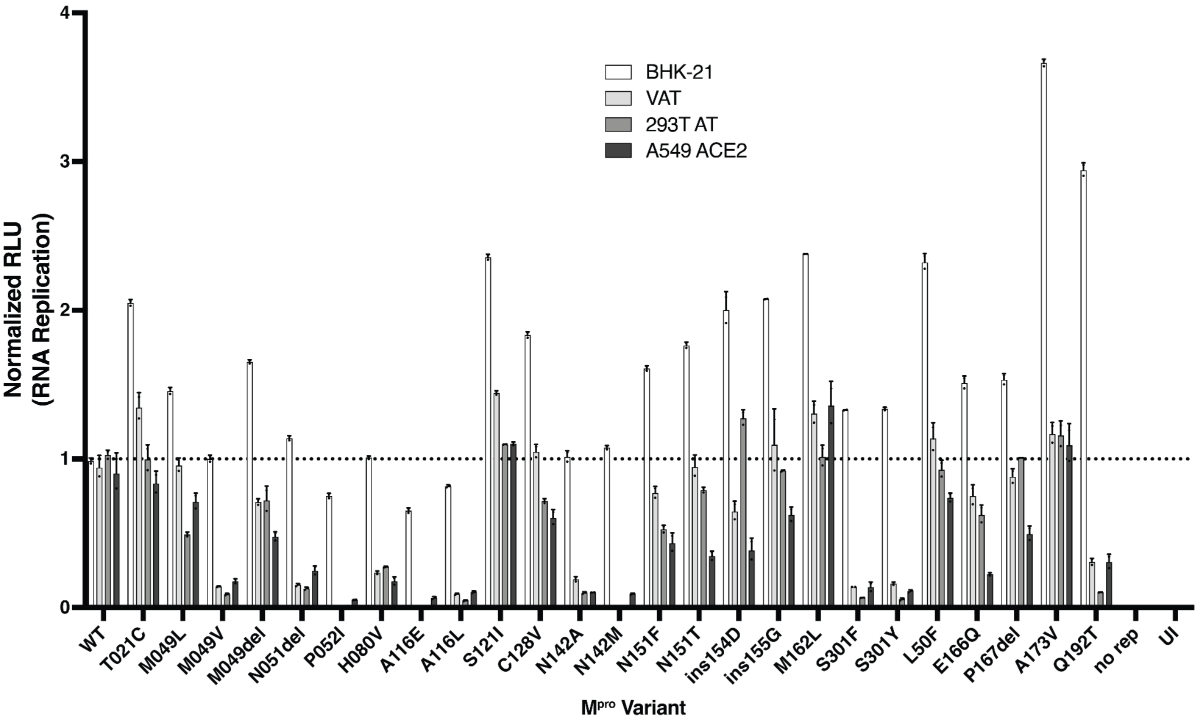


**Figure S26. SARS-CoV-2 $M^{pro}$ variant-dependent RNA replication across cell types.** Bar plots showing normalized SARS-CoV-2 replicon RNA replication for $M^{pro}$ variants in four cell types: BHK-21, VAT, 293T-AT, and A549-ACE2. Variants were selected based on HT-MEK$^{pro}$ measurements (hyperactive or ensitrelvir-resistant with near–WT catalytic efficiency). Replication was quantified using a luciferase-based reporter and normalized to WT within BHK-21. Bars and error bars represent the mean and standard deviation of two replicates, respectively.

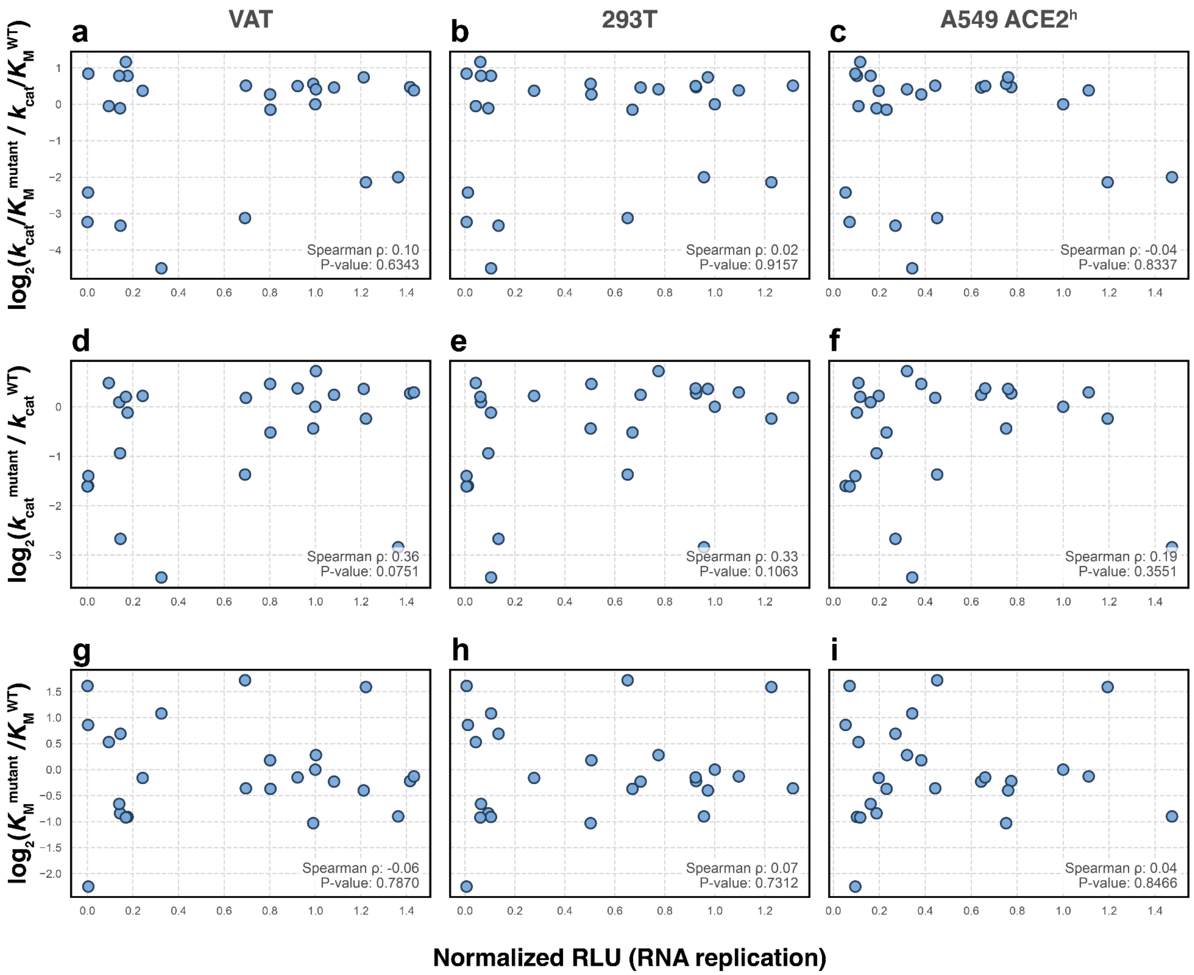


**Figure S27. Correlation of SARS-CoV-2 M$^{pro}$ catalytic parameters with viral RNA replication.** Correlation between HT-MEK$^{pro}$–measured catalytic parameters and viral RNA replication in three infection-based assays. Each point represents an individual M$^{pro}$ variant tested in VAT, 293T-AT, or A549-ACE2h cells. **(a–c)** $\log_2$ fold change in catalytic efficiency ($k_{cat}/K_M$) versus normalized RNA replication for VAT (a), 293T-AT (b), and A549-ACE2$^h$ (c). Panel (c) reproduces the data shown in Fig. 5c. **(d–f)** $\log_2$ fold change in turnover number ($k_{cat}$) versus RNA replication in VAT (d), 293T-AT (e), and A549-ACE2h (f). **(g–i)** $\log_2$ fold change in Michaelis constant ($K_M$) versus RNA replication in VAT (g), 293T-AT (h), and A549-ACE2h (i). Across all nine comparisons, replication showed weak or no correlation with any single biochemical parameter (all Spearman $\rho < 0.4$; no significant p-values).

| | P6 | P5 | P4 | P3 | P2 | P1 | P1’ | P2’ | P3’ | P4’ | P5’ | P6’ |
|---|---|---|---|---|---|---|---|---|---|---|---|---|
| **NSP4 \| NSP5** | T | S | A | V | L | Q | S | G | F | R | K | M |
| **NSP5 \| NSP6** | S | G | V | T | F | Q | S | A | V | K | R | T |
| **NSP6 \| NSP7** | K | V | A | T | V | Q | S | K | M | S | D | V |
| **NSP7 \| NSP8** | N | R | A | T | L | Q | A | I | A | S | E | F |
| **NSP8 \| NSP9** | S | A | V | K | L | Q | N | N | E | L | S | P |
| **NSP9 \| NSP10** | A | T | V | R | L | Q | A | G | N | A | T | E |
| **NSP10 \| NSP11** | R | E | P | M | L | Q | S | A | D | A | Q | S |
| **NSP12 \| NSP13** | P | H | T | V | L | Q | A | V | G | A | C | V |
| **NSP13 \| NSP14** | N | V | A | T | L | Q | A | E | N | V | T | G |
| **NSP14 \| NSP15** | T | F | T | R | L | Q | S | L | E | N | V | A |
| **NSP15 \| NSP16** | T | Y | P | K | L | Q | S | S | Q | A | W | Q |

**Figure S28. Junction peptide sequences for the 11 M$^{pro}$ cutsites in the viral polyproteins pp1a and pp1ab.**

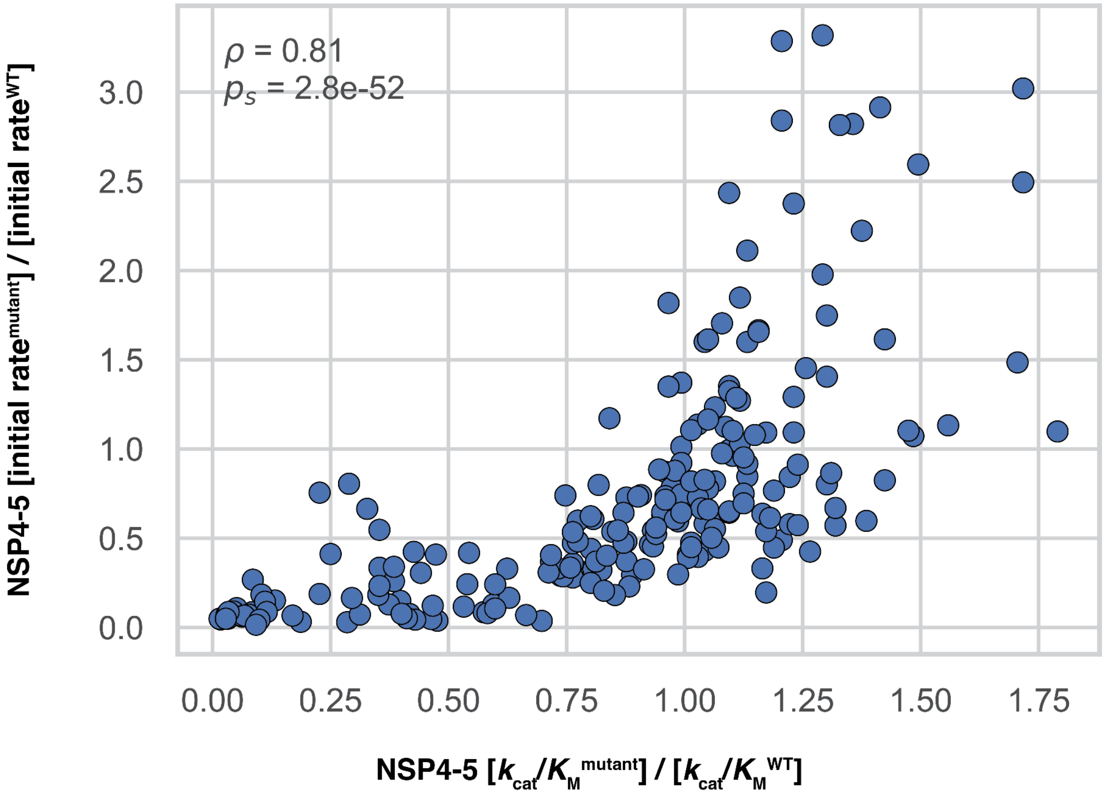


**Figure S29. Relationship between NSP4–5 initial rates and catalytic efficiency.**
Scatter plot comparing initial rates for the NSP4–5 substrate relative to WT measured at 15 μM on the HT-MEK$^{pro}$ platform (y-axis) with $k_{cat}/K_M$ values relative to WT from full Michaelis–Menten fits from an independent experiment (x-axis). Each point represents a differnt variant. The Spearman correlation coefficient (ρ = 0.81) and p-value (p = 2.8 × 10$^{-52}$) are indicated on the plot.

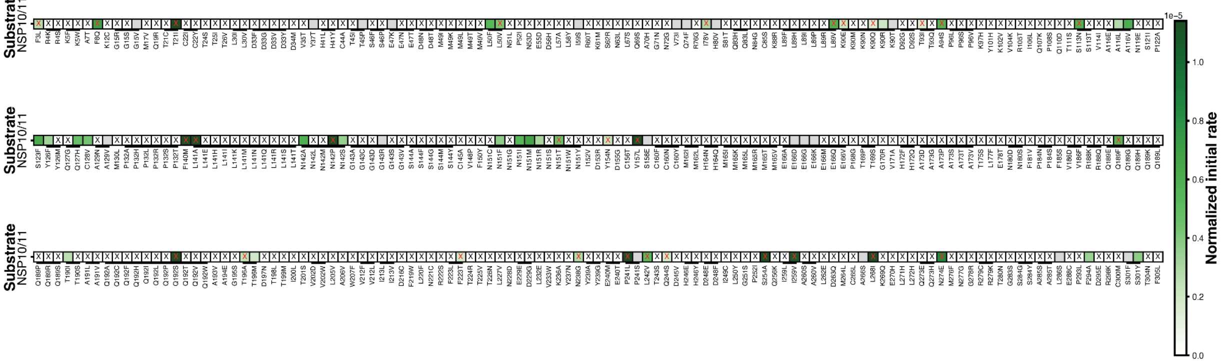


**Figure S30. Initial rates for NSP10–11 peptides for SARS-CoV-2 M$^{pro}$ variants.**
Heatmap analogous to Fig. 6a showing initial rates for NSP10–11 peptides measured at 15 μM substrate. Color scale (green) indicates the average initial rate per chamber. Black X's mark conditions where either WT or the mutant had undetectable initial rates. Red X's mark conditions where only a single biological replicate passed quality filters for that variant–run combination.

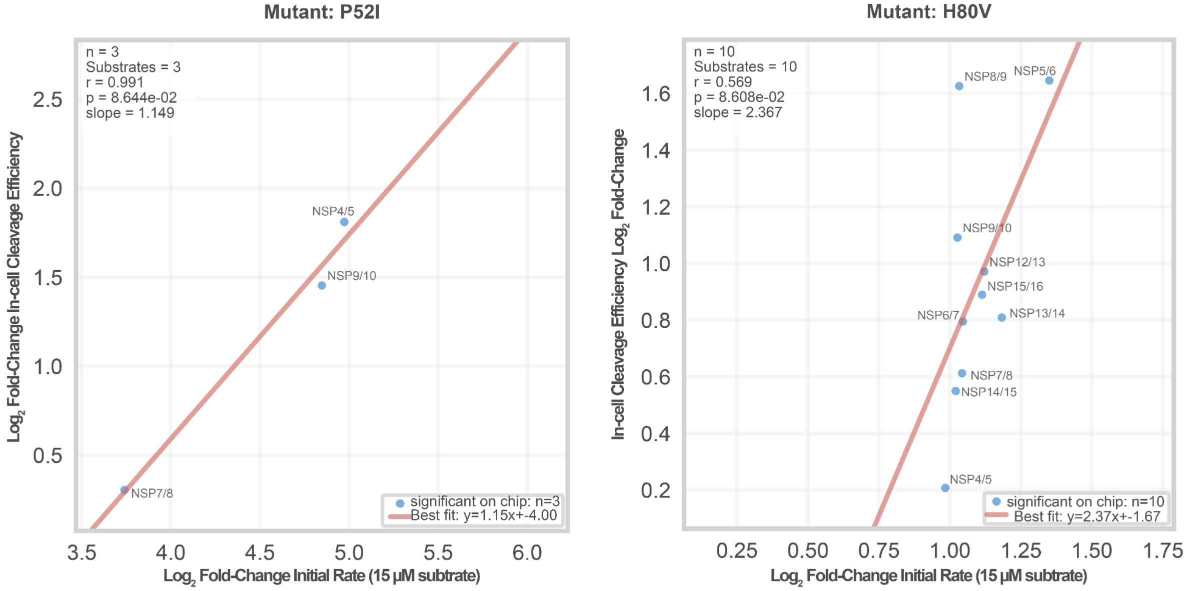


**Figure S31. Concordance between *in vitro* substrate specificity measured by HT-MEK$^{pro}$ and in-cell luciferase cleavage assays for highly perturbed variants P52I and H80V.** Scatter plots comparing the absolute value of the log2 fold changes between *in vitro* and in-cell cleavage efficiency for two highly perturbed mutants.

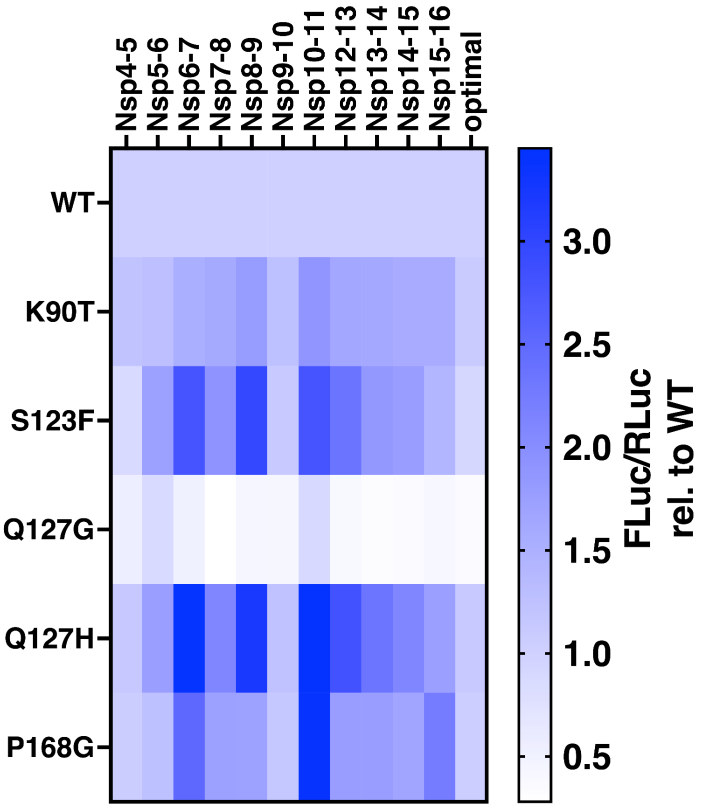


**Figure S32. In-cell cleavage profiles of M$^{pro}$ variants prioritized by *in vitro* specificity measurements.**

Five SARS-CoV-2 M$^{pro}$ variants were selected from HT-MEK$^{pro}$ assays based on divergent *in vitro* specificity profiles and analyzed using a split-luciferase reporter assay measuring cleavage across all eleven native polyprotein junctions plus a non-native optimized sequence (see Methods).

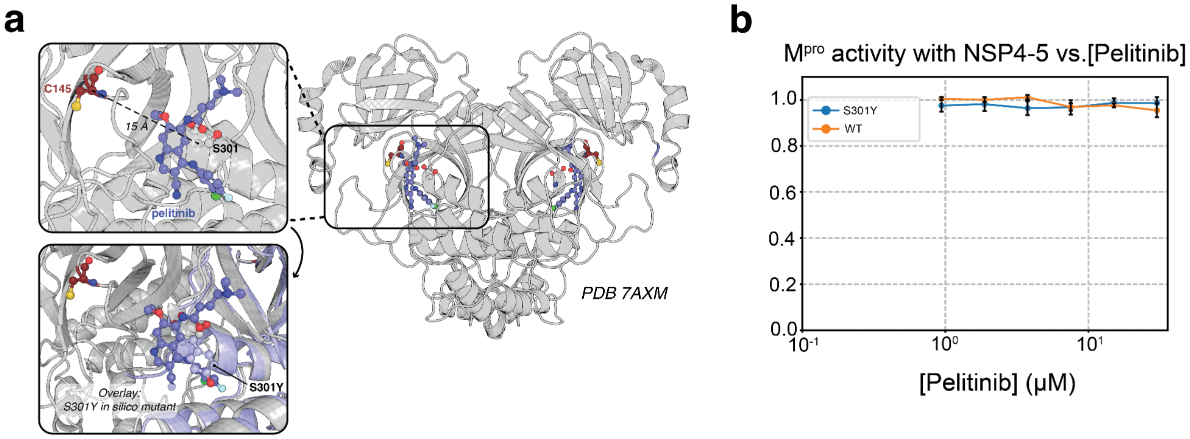


**Figure S33. Allosteric engagement of M$^{pro}$ by pelitinib does not directly inhibit catalysis.** **(a)** Structure of SARS-CoV-2 M$^{pro}$ illustrating the allosteric pocket between domains 2 and 3 that contains residue S301 and the bound kinase inhibitor pelitinib. An overlay of the predicted S301Y structure (mutation made *in silico*; see Methods) is shown in light blue. The allosteric site is ~15 Å from the catalytic cysteine C145. **(b)** Effect of pelitinib on M$^{pro}$ catalytic activity *in vitro*. Normalized initial rates of NSP4–5 cleavage by WT and S301Y M$^{pro}$ as a function of pelitinib concentration across a range spanning the reported antiviral $EC_{50}$ (1.25 μM). Points show mean values from 3 replicate measurements; error bars represent standard deviation.

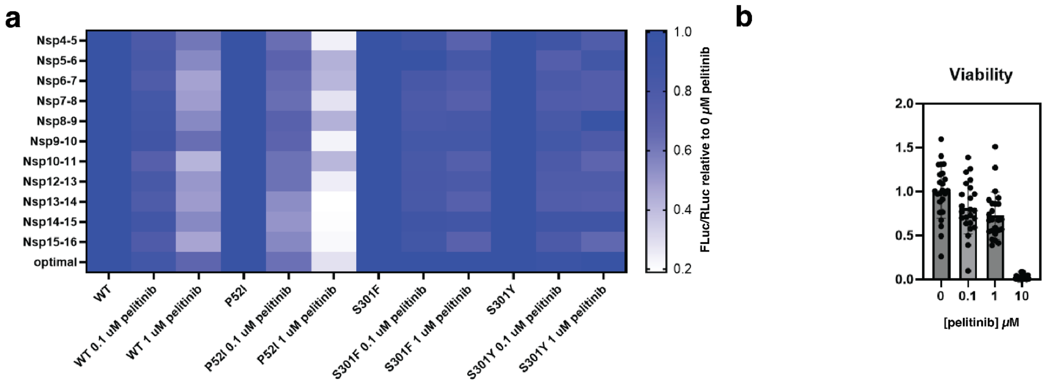


**Figure S34. Effects of pelitinib on SARS-CoV-2 M$^{pro}$–mediated NSP cleavage in cells.** **(a)** Split-luciferase–based cleavage efficiencies for the 11 NSP junctions plus the optimal sequence were measured in cells across a range of pelitinib concentrations for wild-type (WT) M$^{pro}$ and the P52I, S301Y, and S301F variants. For each NSP junction, the ratio of firefly luciferase to renilla luciferase values at each pelitinib concentration were normalized to the corresponding value at 0 μM pelitinib. **(b)** Pelitinib concentrations were kept below the cell viability limit, as determined by cell viability assays described in the Methods.

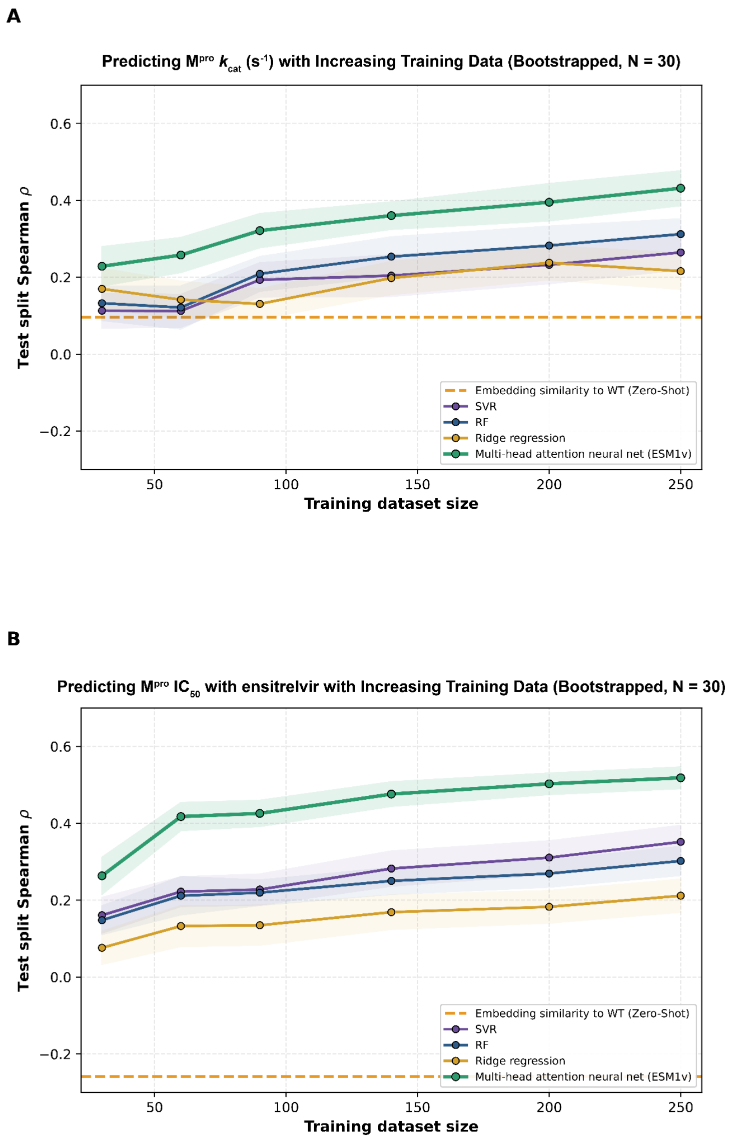


**Figure S35. $M^{pro}$ kcat and $IC_{50}$ prediction improves with supervised learning on HT-MEK$^{pro}$ data.**

(a–b) The mean test Spearman correlation (ρ) for different models across 30 bootstraps is plotted against training dataset size for (a) $k_{cat}$ and (b) ensitrelvir $IC_{50}$. Models include random forest (RF), support vector regression (SVR), ridge regression, and a multi-head attention neural net trained on sequence embeddings (ESM-1v), alongside a zero-shot baseline (cosine similarity). Points and lines denote the mean across bootstrapped train/test splits; shaded regions indicate 95% confidence intervals.

**Supplementary Tables:**

**Table S1. DNA coding sequence and translation of FLAG-TEV-Mpro-SG25-eGFP.**

| **Coding Sequence** |
|---|
| ATGGATTACAAAGATGACGATGACAAGGAAAACCTGTATTTTCAGTCGGGGTTTCGCAAA<br>ATGGCGTTTCCGTCCGGCAAAGTGGAAGGTTGTATGGTACAGGTGACATGCGGCACCA<br>CAACCCTGAATGGGTTGTGGTTAGACGACGTAGTCTATTGCCCTCGTCATGTCATCTGCA<br>CCTCTGAGGACATGCTCAATCCGAATTACGAAGATCTCCTGATCCGCAAATCCAACCACA<br>ACTTCCTCGTTCAAGCTGGCAATGTTCAACTGCGCGTTATTGGTCATAGCATGCAGAATT<br>GCGTCCTTAAACTGAAAGTTGATACCGCCAACCCGAAAACCCCGAAGTACAAGTTTGTG<br>CGTATTCAGCCTGGTCAGACCTTTTCAGTGTTAGCGTGCTATAACGGCAGTCCCTCTGGT<br>GTGTATCAGTGTGCTATGCGTCCAAACTTCACGATCAAAGGCAGCTTCCTTAATGGCAGC<br>TGTGGTTCGGTGGGCTTTAACATCGACTACGATTGCGTTAGCTTCTGCTATATGCACCAC<br>ATGGAATTGCCGACTGGTGTACATGCCGGGACGGACTTAGAAGGCAACTTTTATGGTCC<br>GTTTGTCGATCGCCAAACAGCCCAAGCCGCAGGAACCGATACGACCATTACTGTGAATG<br>TTCTGGCATGGCTTTATGCGGCGGTAATCAACGGAGATCGGTGGTTTCTGAACCGCTTT<br>ACCACCACCTTGAACGATTTCAACCTGGTCGCGATGAAGTACAACTACGAGCCACTGAC<br>GCAGGATCATGTTGACATTCTGGGACCGTTGTCAGCACAGACGGGCATTGCTGTGCTGG<br>ATATGTGTGCCAGTCTGAAAGAACTGCTGCAGAATGGGATGAATGGCCGCACTATTCTG<br>GGTAGTGCGTTACTGGAAGATGAGTTCACTCCCTTCGATGTGGTGCGTCAGTGTAGCGG<br>TGTCACGTTTGCGGGTGGCGGCTCCGGTGGTGGCGGATCTGGTGGCGGTGGATCGGG<br>TGGGGGCGGTTCAGGTGGTGGTGGAAGCGGAATGGTGAGCAAGGGCGAGGAGCTGTT<br>CACCGGGGTGGTGCCCATCCTGGTCGAGCTGGACGGCGACGTAAACGGCCACAAGTTC<br>AGCGTGTCCGGCGAGGGCGAGGGCGATGCCACCTACGGCAAGCTGACCCTGAAGTTCA<br>TCTGCACCACCGGCAAGCTGCCCGTGCCCTGGCCCACCCTCGTGACCACCCTGACCTA<br>CGGCGTGCAGTGCTTCAGCCGCTACCCCGACCACATGAAGCAGCACGACTTCTTCAAGT<br>CCGCCATGCCCGAAGGCTACGTCCAGGAGCGCACCATCTTCTTCAAGGACGACGGCAA<br>CTACAAGACCCGCGCCGAGGTGAAGTTCGAGGGCGACACCCTGGTGAACCGCATCGAG<br>CTGAAGGGCATCGACTTCAAGGAGGACGGCAACATCCTGGGGCACAAGCTGGAGTACA<br>ACTACAACAGCCACAACGTCTATATCATGGCCGACAAGCAGAAGAACGGCATCAAGGTG<br>AACTTCAAGATCCGCCACAACATCGAGGACGGCAGCGTGCAGCTCGCCGACCACTACC<br>AGCAGAACACCCCCATCGGCGACGGCCCCGTGCTGCTGCCCGACAACCACTACCTGAG<br>CACCCAGTCCGCCCTGAGCAAAGACCCCAACGAGAAGCGCGATCACATGGTCCTGCTG<br>GAGTTCGTGACCGCCGCCGGGATCACTCTCGGCATGGACGAGCTGTACAAATAA |
| **Translation** |
| MDYKDDDDKENLYFQSGFRKMAFPSGKVEGCMVQVTCGTTTLNGLWLDDVVYCPRHVICT<br>SEDMLNPNYEDLLIRKSNHNFLVQAGNVQLRVIGHSMQNCVLKLKVDTANPKTPKYKFVRIQ<br>PGQTFSVLACYNGSPSGVYQCAMRPNFTIKGSFLNGSCGSVGFNIDYDCVSFCYMHHMELP<br>TGVHAGTDLEGNFYGPFVDRQTAQAAGTDTTITVNVLAWLYAAVINGDRWFLNRFTTTLNDF<br>NLVAMKYNYEPLTQDHVDILGPLSAQTGIAVLDMCASLKELLQNGMNGRTILGSALLEDEFTP<br>FDVVRQCSGVTFAGGGSGGGGSGGGGSGGGGSGGGGSGMVSKGEELFTGVVPILVELDG<br>DVNGHKFSVSGEGEGDATYGKLTLKFICTTGKLPVPWPTLVTTLTYGVQCFSRYPDHMKQH<br>DFFKSAMPEGYVQERTIFFKDDGNYKTRAEVKFEGDTLVNRIELKGIDFKEDGNILGHKLEYN<br>YNSHNVYIMMPROQKNGIKVNFKIRHNIEDGSVQLADHYQQNTPIGDGPVLLPDNHYLSTQS<br>ALSKDPNEKRDHMVLLEFVTAAGITLGMDELYK* |

**Table S2. Amino acid sequences of related coronavirus proteases**

| | |
|---|---|
| **CoV1** | SGFRKMAFPSGKVEGCMVQVTCGTTTLNGLWLDDTVYCPRHVVCTAEDMLNP<br>NYDDLLIRKSNHSFLVQAGNVQLRVIGHSMQNCLLRLKVDTSNPKTPKYKFVRIQ<br>PGQTFSVLACYNGSPSGVYQCAMRPNHTIKGSFLNGSCGSVGFNIDYDCVSFC<br>YMHHMELPTGVHAGTDLEGKFYGPFVDRQTAQAAGTDTTITLNVLAWLYAAVIN<br>GDRWFLNRFTTTLNDFNLVAMKYNYEPLTQDHVDILGPLSAQTGIAVLDMCAAL<br>KELLQNGMNGRTILGSTILEDEFTPFDVVRQCSGVTFQ |
| **MERS** | SGLVKMSHPSGDVEACMVQVTCGSMTLNGLWLDNTVWCPRHVMCPADQLSD<br>PNYDALLISMTNHSFSVQKHIGAPANLRVVGHAMQGTLLKLTVDVANPSTPAYT<br>FTTVKPGAAFSVLACYNGRPTGTFTVVMRPNYTIKGSFLCGSCGSVGYTKEGS<br>VINFCYMHQMELANGTHTGSAFDGTMYGAFMDKQVHQVQLTDKYCSVNVVAW<br>LYAAILNGCAWFVKPNRTSVVSFNEWALANQFTEFVGTQSVDMLAVKTGVAIEQ<br>LLYAIQQLYTGFQGKQILGSTMLEDEFTPEDVNMQIMGVVMQ |
| **NL63** | SGLKKMAQPSGCVERCVVRVCYGSTVLNGVWLGDTVTCPRHVIAPSTTVLIDY<br>DHAYSTMRLHNFSVSHNGVFLGVVGVTMHGSVLRIKVSQSNVHTPKHVFKTLK<br>PGDSFNILACYEGIASGVFGVNLRTNFTIKGSFINGACGSPGYNVRNDGTVEFCY<br>LHQIELGSGAHVGSDFTGSVYGNFDDQPSLQVESANLMLSDNVVAFLYAALLN<br>GCRWWLCSTRVNVDGFNEWAMANGYTSVSSVECYSILAAKTGVSVEQLLASIQ<br>HLHEGFGGKNILGYSSLCDEFTLAEVVKQMYGVNLQ |
| **OC43** | SGIVKMVNPTSKVEPCVVSVTYGNMTLNGLWLDDKVYCPRHVICSASDMTNPD<br>YTNLLCRVTSSDFTVLFDRLSLTVMSYQMRGCMLVLTVTLQNSRTPKYTFGVVK<br>PGETFTVLAAYNGKPQGAFHVTMRSSYTIKGSFLCGSCGSVGYVIMGDCVKFV<br>YMHQLELSTGCHTGTDFNGDFYGPYKDAQVVQLPIQDYIQSVNFLAWLYAAILN<br>NCNWFIQSDKCSVEDFNVWALSNGFSQVKSDLVIDALASMTGVSLETLLAAIKR<br>LKNGFQGRQIMGSCSFEDELTPSDVYQQLAGIKLA |
| **229E** | AGLRKMAQPSGFVEKCVVRVCYGNTVLNGLWLGDIVYCPRHVIASNTTSAIDYD<br>HEYSIMRLHNFSIISGTAFLGVVGATMHGVTLKIKVSQTNMHTPRHSFRTLKSGE<br>GFNILACYDGCAQGVFGVNMRTNWTIRGSFINGACGSPGYNLKNGEVEFVYMH<br>QIELGSGSHVGSSFDGVMYGGFEDQPNLQVESANQMLTVNVVAFLYAAILNGC<br>TWWLKGEKLFVEHYNEWAQANGFTAMNGEDAFSILAAKTGVCVERLLHAIQVL<br>NNGFGGKQILGYSSLNDEFSINEVVKQMFGVNLQ |
| **HKU1** | SGIVKMVSPTSKIEPCIVSVTYGSMTLNGLWLDDKVYCPRHVICLSSNMNEPDY<br>SALLCRVTLGDFTIMSGRMSLTVVSYQMQGCQLVLTVSLQNPYTPKYTFGVVKP<br>GETFTVLAAYNGRPQGAFHVTMRSSYTIKGSFLCGSCGSVGYVLTGDSVKFVY<br>MHQLELSTGCHTGTDFTGNFYGPYRDAQVVQLPVKDYVQTVNVIAWLYAAILN<br>NCAWFVQNDVCSIEDFNVWAMTNGFSQVKADLVLDALASMTGVSIETLLAAIKR<br>LYMGFQGRQILGSCTFEDELAPSDVYQQLAGVKLQ |
| **HKU4** | SGLVKMSAPSGAVENCIVQVTCGSMTLNGLWLDNTVWCPRHIMCPADQLTDP<br>NYDALLISKTNHSFIVQKHIGAQANLRVVAHSMVGVLLKLTVDVANPSTPAYTFS<br>TVKPGASFSVLACYNGKPTGVFTVNLRHNSTIKGSFLCGSCGSVGYTENGGVIN<br>FVYMHQMELSNGTHTGSSFDGVMYGAFEDKQTHQLQLTDKYCTINVVAWLYA<br>AVLNGCKWFVKPTRVGIVTYNEWALSNQFTEFVGTQSIDMLAHRTGVSVEQML<br>AAIQSLHAGFQGKTILGQSTLEDEFTPDDVNMQVMGVVMQ |
| **HKU5** | SGLVKMAAPSGVVENCMVQVTCGSMTLNGLWLDNYVWCPRHVMCPADQLSD<br>PNYDALLVSKTNLSFIVQKNVGAPANLRVVGHTMVGTLLKLTVESANPQTPAYT<br>FTTVKPGASFSVLACYNGRPTGVFMVNMRQNSTIKGSFLCGSCGSVGYTQEGN<br>VINFCYMHQMELSNGTHTGCAFDGVMYGAFEDRQVHQVQLSDKYCTINIVAWL<br>YAAILNGCNWFVKPNKTGIATFNEWAMSNQFTEFIGTQSVDMLAHKTGVSVEQL<br>LYAIQTLHKGFQGKTILGNSMLEDEFTPDDVNMQVMGVVMQ |

| | |
|---|---|
| **HKU9** | AGLTRMAHPSGLVEPCLVKVNYGSMTLNGIWLDNFVICPRHVMCSRDELANPD<br>YPRLSMRAANYDFHVSQNGHNIRVIGHTMEGSLLKLTVDVNNPKTPAYSFIRVS<br>TGQAMSLLACYDGLPTGVYTCTLRSNGTMRASFLCGSCGSPGFVMNGKEVQF<br>CYLHQLELPNGTHTGTDFSGVFYGPFEDKQVPQLAAPDCTITVNVLAWLYAAVL<br>SGENWFLTKSSISPAEFNNCAVKYMCQSVTSESLQVLQPLAAKTGISVERMLSA<br>LKVLLSAGFCGRTIMGSCSLEDEHTPYDIGRQMLGVKLQ |
| **HKU15** | AGIKILLHPSGVVERCMVSVVYNGSALNGIWLKNVVYCPRHVIGKFRGDQWTH<br>MVSIADCRDFIVKCPIQGIQLNVQSVKMVGALLQLTVHTNNTATPDYKFERLQPG<br>SSMTIACAYDGIVRHVYHVVLQLNNLIYASFLNGACGSVGYTLKGKTLYLHYMHH<br>IEFNNKTHSGTDLEGNFYGPYVDEEVIQQQTAFQYYTDNVVAQLYAHLLTVDAR<br>PKWLAQSQISIEDFNSWAANNSFANFPCEQTNMSYIMGLSQTARVPVERILNTII<br>QLTTNRDGACIMGSYDFECDWTPEMVYNQAPISLQ |
| **MHV** | SGIVKMVSPTSKVEPCIVSVTYGNMTLNGLWLDDKVYCPRHVICSSADMTDPDY<br>PNLLCRVTSSDFCVMSGRMSLTVMSYQMQGCQLVLTVTLQNPNTPKYSFGVV<br>KPGETFTVLAAYNGRPQGAFHVTLRSSHTIKGSFLCGSCGSVGYVLTGDSVRFV<br>YMHQLELSTGCHTGTDFSGNFYGPYRDAQVVQLPVQDYTQTVNVVAWLYAAIF<br>NRCNWFVQSDSCSLEEFNVWAMTNGFSSIKADLVLDALASMTGVTVEQVLAAI<br>KRLHSGFQGKQILGSCVLEDETPSDVYQQLAGVKLQ |
| **FIPV** | SGLRKMAQPSGVVEPCIVRVAYGNNVLNGLWLGDEVICPRHVIASDTSRVINYE<br>NELSSVRLHNFSIAKNNAFLGVVSAKYKGVNLVLKVNQVNPNTPEHKFKSVRPG<br>ESFNILACYEGCPGSVYGVNMRSQGTIKGSFIAGTCGSVGYVLENGTLYFVYMH<br>HLELGNGSHVGSNLEGEMYGGYEDQPSMQLEGTNVMSSDNVVAFLYAALING<br>ERWFVTNTSMTLESYNAWAKTNSFTEIVSTDAFNMLAAKTGYSVEKLLECIVRL<br>NKGFGGRTILSYGSLCDEFTPTEVIRQMYGVNLQ |
| **IBV** | SGFKKLVSPSSAVEKCIVSVSYRGNNLNGLWLGDTIYCPRHVLGKFSGDQWND<br>VLNLANNHEFEVTTQHGVTLNVVSRRLKGAVLILQTAVANAETPKYKFIKANCGD<br>SFTIACAYGGTVVGLYPVTMRSNGTIRASFLAGACGSVGFNIEKGVVNFFYMHH<br>LELPNALHTGTDLMGEFYGGYVDEEVAQRVPPDNLVTNNIVAWLYAAIISVKESS<br>FSLPKWLESTTVSVDDYNKWAGDNGFTPFSTSTAITKLSAITGVDVCKLLRTIMV<br>KNSQWGGDPILGQYNFEDELTPESVFNQIGGVRLQ |
| **PEDV** | AGLRKMAQPSGVVEKCIVRVCYGNMALNGLWLGDIVMCPRHVIASSTTSTIDYD<br>YALSVLRLHNFSISSGNVFLGVVSATMRGALLQIKVNQNNVHTPKYTYRTVRPG<br>ESFNILACYDGAAAGVYGVNMRSNYTIRGSFINGACGSPGYNINNGTVEFCYLH<br>QLELGSGCHVGSDLDGVMYGGYEDQPTLQVEGASSLFTENVLAFLYAALINGS<br>TWWLSSSRIAVDRFNEWAVHNGMTTVGNTDCFSILAAKTGVDVQRLLASIQSLH<br>KNFGGKQILGHTSLTDEFTTGEVVRQMYGVNLQ |

**Table S3. $M^{pro}$ Ortholog Michaelis-Menten, Ensitrelvir, and AVI-4516 Inhibition Values.**

| Mutant ID | Measurement | Value | SD | log2FC | -log10 (q-value) |
|---|---|---|---|---|---|
| HKU4-WT | kcat (s-1) | 2.45 | 0.56 | -0.61 | 1.67 |
| | KM (μM) | 45.18 | 9.62 | -0.79 | 2.81 |
| | kcat/KM (M-1s-1) | 5.56E+04 | 1.47E+04 | 0.19 | 0.89 |
| | $IC_{50}$ (M) | 5.26E-09 | 8.85E-10 | 2.30 | 6.25 |
| FIPV-WT | kcat (s-1) | 6.50 | 1.12 | 0.80 | 3.38 |
| | KM (μM) | 38.80 | 8.18 | -1.01 | 2.93 |
| | kcat/KM (M-1s-1) | 1.70E+05 | 1.85E+04 | 1.80 | 5.83 |
| | $IC_{50}$ (M) | 6.70E-07 | 7.04E-08 | 9.29 | 7.89 |
| IBV-WT | kcat (s-1) | 11.74 | 2.09 | 1.66 | 4.71 |
| | KM (μM) | 112.82 | 26.21 | 0.53 | 1.80 |
| | kcat/KM (M-1s-1) | 1.06E+05 | 1.16E+04 | 1.12 | 5.41 |
| | $IC_{50}$ (M) | 2.53E-05 | 7.91E-06 | 14.53 | 4.50 |
| MHV-WT | kcat (s-1) | Not Active | -- | -- | -- |
| | KM (μM) | Not Active | -- | -- | -- |
| | kcat/KM (M-1s-1) | Not Active | -- | -- | -- |
| | $IC_{50}$ (M) | Not Active | -- | -- | -- |
| PEDV-WT | kcat (s-1) | 10.57 | 1.49 | 1.50 | 4.49 |
| | KM (μM) | 31.40 | 4.18 | -1.32 | 2.69 |
| | kcat/KM (M-1s-1) | 3.41E+05 | 5.94E+04 | 2.81 | 4.33 |
| | $IC_{50}$ (M) | 2.38E-07 | 1.93E-08 | 7.80 | 7.54 |
| SARS1-WT | kcat (s-1) | 3.36 | 0.87 | -0.15 | 0.33 |
| | KM (μM) | 81.73 | 14.79 | 0.06 | 0.62 |
| | kcat/KM (M-1s-1) | 4.09E+04 | 6.73E+03 | -0.25 | 1.08 |
| | $IC_{50}$ (M) | 7.69E-10 | 3.35E-10 | -0.47 | 1.27 |
| MERS-WT | kcat (s-1) | 1.13 | 0.21 | -1.72 | 3.66 |
| | KM (μM) | 70.37 | 21.71 | -0.15 | 0.54 |
| | kcat/KM (M-1s-1) | 1.68E+04 | 3.52E+03 | -1.53 | 3.78 |
| | $IC_{50}$ (M) | 7.39E-09 | 7.75E-10 | 2.79 | 6.99 |
| NL63-WT | kcat (s-1) | 3.19 | 1.05 | -0.23 | 0.64 |
| | KM (μM) | 21.92 | 3.33 | -1.84 | 4.24 |
| | kcat/KM (M-1s-1) | 1.52E+05 | 5.43E+04 | 1.64 | 2.98 |
| | $IC_{50}$ (M) | 1.12E-05 | 2.21E-06 | 13.36 | 6.02 |
| OC43-WT | kcat (s-1) | 0.65 | 0.26 | -2.52 | 5.04 |
| | KM (μM) | 70.23 | 6.69 | -0.16 | 0.57 |
| | kcat/KM (M-1s-1) | 9.39E+03 | 4.16E+03 | -2.37 | 4.31 |

| | | | | | |
|---|---|---|---|---|---|
| | $IC_{50}$ (M) | 9.89E-09 | 3.85E-09 | 3.21 | 3.27 |
| HKU1-WT | kcat (s-1) | 0.08 | 0.02 | -5.47 | 2.15 |
| | KM (μM) | 45.65 | 12.03 | -0.78 | 1.16 |
| | kcat/KM (M-1s-1) | 1.89E+03 | 3.74E+02 | -4.69 | 2.48 |
| | $IC_{50}$ (M) | 2.87E-08 | 5.87E-08 | 4.75 | 0.75 |
| 2293E-WT | kcat (s-1) | 0.39 | 0.25 | -3.25 | 4.36 |
| | KM (μM) | 15.18 | 2.99 | -2.37 | 4.06 |
| | kcat/KM (M-1s-1) | 2.55E+04 | 1.70E+04 | -0.93 | 1.67 |
| | $IC_{50}$ (M) | 1.69E-06 | 3.50E-07 | 10.63 | 5.14 |
| HKU5-WT | kcat (s-1) | 1.20 | 0.48 | -1.63 | 3.46 |
| | KM (μM) | 63.28 | 19.71 | -0.31 | 0.73 |
| | kcat/KM (M-1s-1) | 1.83E+04 | 5.07E+03 | -1.41 | 3.23 |
| | $IC_{50}$ (M) | 6.03E-10 | 1.64E-10 | -0.82 | 1.61 |
| HKU9-WT | kcat (s-1) | Not Active | -- | -- | -- |
| | KM (μM) | Not Active | -- | -- | -- |
| | kcat/KM (M-1s-1) | Not Active | -- | -- | -- |
| | $IC_{50}$ (M) | Not Active | -- | -- | -- |
| HKU15-WT | kcat (s-1) | 7.56 | 0.83 | 1.02 | 3.94 |
| | KM (μM) | 114.00 | 9.16 | 0.54 | 2.58 |
| | kcat/KM (M-1s-1) | 6.63E+04 | 4.02E+03 | 0.44 | 2.61 |
| | $IC_{50}$ (M) | 1.55E-04 | 7.81E-05 | 17.15 | 1.72 |

307

**Table S4. Michaelis-Menten parameters for functional and clinically focused sublibrary**

| Mutant ID | Measurement | Value | SD | log2FC | -log10 (q-value) |
|---|---|---|---|---|---|
| SARS2-WT | kcat (s-1) | 4.54 | 1.05 | 0.00 | N/A |
| | KM (μM) | 49.04 | 14.04 | 0.00 | N/A |
| | kcat/KM (M-1s-1) | 9.46E+04 | 1.04E+04 | 0.00 | N/A |
| SARS2-C145A | kcat (s-1) | Not Active | -- | -- | -- |
| | KM (μM) | Not Active | -- | -- | -- |
| | kcat/KM (M-1s-1) | Not Active | -- | -- | -- |
| SARS2-TQA-WT | kcat (s-1) | 1.11 | 0.33 | -2.03 | 4.55 |
| | KM (μM) | 54.79 | 12.92 | 0.16 | 0.69 |
| | kcat/KM (M-1s-1) | 2.04E+04 | 5.09E+03 | -2.21 | 6.46 |
| SARS2-TQA-C145A | kcat (s-1) | Not Active | -- | -- | -- |
| | KM (μM) | Not Active | -- | -- | -- |
| | kcat/KM (M-1s-1) | Not Active | -- | -- | -- |
| SARS2-R004S | kcat (s-1) | Not Active | -- | -- | -- |
| | KM (μM) | Not Active | -- | -- | -- |
| | kcat/KM (M-1s-1) | Not Active | -- | -- | -- |
| SARS2-K005F | kcat (s-1) | 1.90 | 0.37 | -1.25 | 3.86 |
| | KM (μM) | 36.42 | 10.29 | -0.43 | 1.59 |
| | kcat/KM (M-1s-1) | 5.44E+04 | 1.19E+04 | -0.80 | 4.21 |
| SARS2-K005W | kcat (s-1) | 1.88 | 0.26 | -1.27 | 3.90 |
| | KM (μM) | 41.15 | 3.17 | -0.25 | 1.18 |
| | kcat/KM (M-1s-1) | 4.59E+04 | 7.19E+03 | -1.04 | 6.22 |
| SARS2-A007T | kcat (s-1) | 3.63 | 0.71 | -0.32 | 1.77 |
| | KM (μM) | 40.03 | 6.34 | -0.29 | 1.23 |
| | kcat/KM (M-1s-1) | 9.07E+04 | 1.27E+04 | -0.06 | 1.53 |
| SARS2-G015S | kcat (s-1) | 5.08 | 0.72 | 0.16 | 0.73 |
| | KM (μM) | 54.16 | 12.21 | 0.14 | 0.57 |
| | kcat/KM (M-1s-1) | 9.61E+04 | 1.34E+04 | 0.02 | 0.39 |
| SARS2-G015V | kcat (s-1) | 2.31 | 0.42 | -0.97 | 3.14 |
| | KM (μM) | 33.85 | 8.22 | -0.54 | 1.41 |
| | kcat/KM (M-1s-1) | 6.95E+04 | 8.22E+03 | -0.44 | 3.03 |

| | | | | | |
|---|---|---|---|---|---|
| SARS2-T021I | kcat (s-1) | 4.08 | 1.00 | -0.15 | 0.62 |
| | KM (µM) | 36.23 | 9.99 | -0.44 | 1.62 |
| | kcat/KM (M-1s-1) | 1.16E+05 | 2.51E+04 | 0.29 | 2.09 |
| SARS2-C022I | kcat (s-1) | 0.12 | 0.04 | -5.29 | 5.46 |
| | KM (µM) | 38.33 | 25.98 | -0.36 | 0.67 |
| | kcat/KM (M-1s-1) | 3.61E+03 | 1.27E+03 | -4.71 | 7.35 |
| SARS2-T025I | kcat (s-1) | 2.27 | 0.47 | -1.00 | 3.13 |
| | KM (µM) | 68.19 | 13.86 | 0.48 | 1.49 |
| | kcat/KM (M-1s-1) | 3.34E+04 | 2.15E+03 | -1.50 | 6.24 |
| SARS2-L030I | kcat (s-1) | 3.21 | 0.47 | -0.50 | 2.87 |
| | KM (µM) | 41.37 | 8.22 | -0.25 | 1.48 |
| | kcat/KM (M-1s-1) | 7.84E+04 | 6.50E+03 | -0.27 | 2.80 |
| SARS2-D033F | kcat (s-1) | 3.10 | 0.75 | -0.55 | 1.75 |
| | KM (µM) | 37.94 | 6.26 | -0.37 | 1.48 |
| | kcat/KM (M-1s-1) | 8.17E+04 | 1.41E+04 | -0.21 | 2.21 |
| SARS2-D033V | kcat (s-1) | 3.53 | 0.72 | -0.36 | 2.19 |
| | KM (µM) | 41.29 | 11.99 | -0.25 | 1.20 |
| | kcat/KM (M-1s-1) | 8.75E+04 | 8.96E+03 | -0.11 | 1.03 |
| SARS2-D033Y | kcat (s-1) | Not Active | -- | -- | -- |
| | KM (µM) | Not Active | -- | -- | -- |
| | kcat/KM (M-1s-1) | Not Active | -- | -- | -- |
| SARS2-D034M | kcat (s-1) | 4.13 | 0.51 | -0.13 | 0.87 |
| | KM (µM) | 53.71 | 9.95 | 0.13 | 0.59 |
| | kcat/KM (M-1s-1) | 7.84E+04 | 1.11E+04 | -0.27 | 1.55 |
| SARS2-H041L | kcat (s-1) | Not Active | -- | -- | -- |
| | KM (µM) | Not Active | -- | -- | -- |
| | kcat/KM (M-1s-1) | Not Active | -- | -- | -- |
| SARS2-H041Y | kcat (s-1) | Not Active | -- | -- | -- |
| | KM (µM) | Not Active | -- | -- | -- |
| | kcat/KM (M-1s-1) | Not Active | -- | -- | -- |
| SARS2-T045I | kcat (s-1) | 2.09 | 0.39 | -1.12 | 4.60 |
| | KM (µM) | 44.12 | 9.29 | -0.15 | 0.61 |
| | kcat/KM (M-1s-1) | 4.81E+04 | 6.91E+03 | -0.98 | 5.01 |

| | | | | | |
|---|---|---|---|---|---|
| SARS2-S046F | kcat (s-1) | 5.23 | 0.63 | 0.20 | 1.09 |
| | KM (µM) | 46.75 | 10.44 | -0.07 | 0.42 |
| | kcat/KM (M-1s-1) | 1.16E+05 | 2.72E+04 | 0.30 | 1.40 |
| SARS2-S046P | kcat (s-1) | 4.99 | 0.72 | 0.14 | 0.97 |
| | KM (µM) | 47.77 | 9.41 | -0.04 | 0.38 |
| | kcat/KM (M-1s-1) | 1.06E+05 | 8.96E+03 | 0.16 | 1.86 |
| SARS2-E047K | kcat (s-1) | 4.02 | 0.86 | -0.18 | 1.31 |
| | KM (µM) | 36.29 | 10.70 | -0.43 | 2.17 |
| | kcat/KM (M-1s-1) | 1.14E+05 | 1.70E+04 | 0.27 | 2.05 |
| SARS2-E047N | kcat (s-1) | 4.17 | 0.78 | -0.12 | 0.57 |
| | KM (µM) | 39.22 | 9.07 | -0.32 | 0.99 |
| | kcat/KM (M-1s-1) | 1.10E+05 | 2.68E+04 | 0.22 | 0.99 |
| SARS2-D048N | kcat (s-1) | 3.18 | 0.38 | -0.51 | 1.87 |
| | KM (µM) | 38.67 | 5.00 | -0.34 | 1.15 |
| | kcat/KM (M-1s-1) | 8.27E+04 | 7.08E+03 | -0.19 | 2.42 |
| SARS2-M049I | kcat (s-1) | 1.20 | 0.14 | -1.92 | 4.40 |
| | KM (µM) | 33.07 | 7.71 | -0.57 | 1.67 |
| | kcat/KM (M-1s-1) | 3.75E+04 | 7.27E+03 | -1.33 | 6.46 |
| SARS2-M049K | kcat (s-1) | 1.55 | 0.27 | -1.55 | 4.56 |
| | KM (µM) | 37.16 | 4.04 | -0.40 | 1.65 |
| | kcat/KM (M-1s-1) | 4.17E+04 | 4.05E+03 | -1.18 | 5.42 |
| SARS2-M049L | kcat (s-1) | 3.34 | 0.50 | -0.44 | 2.46 |
| | KM (µM) | 24.01 | 3.87 | -1.03 | 3.03 |
| | kcat/KM (M-1s-1) | 1.40E+05 | 1.31E+04 | 0.56 | 3.58 |
| SARS2-M049T | kcat (s-1) | 1.72 | 0.31 | -1.40 | 4.17 |
| | KM (µM) | 34.24 | 7.54 | -0.52 | 2.33 |
| | kcat/KM (M-1s-1) | 5.13E+04 | 8.49E+03 | -0.88 | 5.49 |
| SARS2-M049V | kcat (s-1) | 2.36 | 0.34 | -0.94 | 3.16 |
| | KM (µM) | 27.34 | 5.41 | -0.84 | 2.32 |
| | kcat/KM (M-1s-1) | 8.75E+04 | 7.14E+03 | -0.11 | 0.88 |
| SARS2-L050F | kcat (s-1) | 5.83 | 0.60 | 0.36 | 1.91 |
| | KM (µM) | 37.11 | 4.66 | -0.40 | 1.53 |
| | kcat/KM (M-1s-1) | 1.58E+05 | 6.57E+03 | 0.74 | 6.31 |

| | | | | | |
|---|---|---|---|---|---|
| SARS2-V073I | kcat (s-1) | Not Active | -- | -- | -- |
| | KM (μM) | Not Active | -- | -- | -- |
| | kcat/KM (M-1s-1) | Not Active | -- | -- | -- |
| SARS2-Q083L | kcat (s-1) | 4.14 | 0.45 | -0.13 | 0.77 |
| | KM (μM) | 41.91 | 3.39 | -0.23 | 0.94 |
| | kcat/KM (M-1s-1) | 9.86E+04 | 5.53E+03 | 0.06 | 0.68 |
| SARS2-K088R | kcat (s-1) | 4.37 | 1.20 | -0.05 | 0.40 |
| | KM (μM) | 42.36 | 10.70 | -0.21 | 0.81 |
| | kcat/KM (M-1s-1) | 1.03E+05 | 8.94E+03 | 0.13 | 2.24 |
| SARS2-L089F | kcat (s-1) | 2.81 | 0.68 | -0.69 | 2.54 |
| | KM (μM) | 52.50 | 18.40 | 0.10 | 0.49 |
| | kcat/KM (M-1s-1) | 5.57E+04 | 8.37E+03 | -0.76 | 5.18 |
| SARS2-L089H | kcat (s-1) | Not Active | -- | -- | -- |
| | KM (μM) | Not Active | -- | -- | -- |
| | kcat/KM (M-1s-1) | Not Active | -- | -- | -- |
| SARS2-L089I | kcat (s-1) | 3.23 | 0.55 | -0.49 | 2.48 |
| | KM (μM) | 39.11 | 10.06 | -0.33 | 1.20 |
| | kcat/KM (M-1s-1) | 8.39E+04 | 7.22E+03 | -0.17 | 1.34 |
| SARS2-L089P | kcat (s-1) | Not Active | -- | -- | -- |
| | KM (μM) | Not Active | -- | -- | -- |
| | kcat/KM (M-1s-1) | Not Active | -- | -- | -- |
| SARS2-L089R | kcat (s-1) | Not Active | -- | -- | -- |
| | KM (μM) | Not Active | -- | -- | -- |
| | kcat/KM (M-1s-1) | Not Active | -- | -- | -- |
| SARS2-L089V | kcat (s-1) | 2.36 | 0.56 | -0.94 | 3.17 |
| | KM (μM) | 44.07 | 9.72 | -0.15 | 0.67 |
| | kcat/KM (M-1s-1) | 5.45E+04 | 1.16E+04 | -0.80 | 4.37 |
| SARS2-K090E | kcat (s-1) | 4.07 | 0.50 | -0.16 | 1.54 |
| | KM (μM) | 46.48 | 8.87 | -0.08 | 0.73 |
| | kcat/KM (M-1s-1) | 8.88E+04 | 9.58E+03 | -0.09 | 0.69 |
| SARS2-K090M | kcat (s-1) | 4.27 | 0.87 | -0.09 | 0.49 |
| | KM (μM) | 47.14 | 14.07 | -0.06 | 0.40 |
| | kcat/KM (M-1s-1) | 9.39E+04 | 2.10E+04 | -0.01 | 0.34 |

| | | | | | |
|---|---|---|---|---|---|
| SARS2-K090N | kcat (s-1) | 3.82 | 0.55 | -0.25 | 1.13 |
| | KM (μM) | 47.24 | 11.47 | -0.05 | 0.55 |
| | kcat/KM (M-1s-1) | 8.27E+04 | 1.07E+04 | -0.19 | 1.44 |
| SARS2-K090Q | kcat (s-1) | 3.81 | 1.00 | -0.25 | 0.90 |
| | KM (μM) | 41.64 | 14.10 | -0.24 | 0.75 |
| | kcat/KM (M-1s-1) | 9.40E+04 | 1.17E+04 | -0.01 | 0.34 |
| SARS2-K090T | kcat (s-1) | 4.65 | 1.08 | 0.04 | 0.32 |
| | KM (μM) | 48.29 | 13.82 | -0.02 | 0.49 |
| | kcat/KM (M-1s-1) | 9.89E+04 | 1.50E+04 | 0.06 | 0.91 |
| SARS2-K090R | kcat (s-1) | 3.90 | 0.69 | -0.22 | 1.28 |
| | KM (μM) | 41.06 | 14.19 | -0.26 | 1.00 |
| | kcat/KM (M-1s-1) | 9.95E+04 | 1.59E+04 | 0.07 | 0.65 |
| SARS2-D092G | kcat (s-1) | 3.99 | 0.75 | -0.19 | 1.09 |
| | KM (μM) | 39.47 | 10.87 | -0.31 | 1.18 |
| | kcat/KM (M-1s-1) | 1.03E+05 | 1.18E+04 | 0.13 | 1.09 |
| SARS2-T093I | kcat (s-1) | 4.88 | 0.77 | 0.11 | 0.53 |
| | KM (μM) | 48.52 | 10.35 | -0.02 | 0.32 |
| | kcat/KM (M-1s-1) | 1.02E+05 | 1.05E+04 | 0.11 | 1.27 |
| SARS2-P096L | kcat (s-1) | 3.88 | 0.48 | -0.22 | 0.90 |
| | KM (μM) | 39.74 | 6.22 | -0.30 | 0.95 |
| | kcat/KM (M-1s-1) | 9.83E+04 | 5.44E+03 | 0.06 | 0.66 |
| SARS2-P096S | kcat (s-1) | Not Active | -- | -- | -- |
| | KM (μM) | Not Active | -- | -- | -- |
| | kcat/KM (M-1s-1) | Not Active | -- | -- | -- |
| SARS2-P108S | kcat (s-1) | 3.04 | 0.89 | -0.58 | 1.94 |
| | KM (μM) | 44.70 | 10.73 | -0.13 | 0.71 |
| | kcat/KM (M-1s-1) | 6.83E+04 | 1.23E+04 | -0.47 | 3.82 |
| SARS2-S113T | kcat (s-1) | 4.53 | 0.56 | 0.00 | 0.31 |
| | KM (μM) | 37.13 | 4.42 | -0.40 | 1.57 |
| | kcat/KM (M-1s-1) | 1.22E+05 | 7.60E+03 | 0.37 | 3.46 |
| SARS2-A116E | kcat (s-1) | 1.49 | 0.29 | -1.61 | 4.52 |
| | KM (μM) | 149.71 | 29.62 | 1.61 | 4.98 |
| | kcat/KM (M-1s-1) | 1.01E+04 | 1.66E+03 | -3.23 | 7.14 |

| | | | | | |
|---|---|---|---|---|---|
| SARS2-A116L | kcat (s-1) | 6.32 | 0.91 | 0.48 | 4.27 |
| | KM (µM) | 70.65 | 14.18 | 0.53 | 3.35 |
| | kcat/KM (M-1s-1) | 9.10E+04 | 1.34E+04 | -0.05 | 0.58 |
| SARS2-A116V | kcat (s-1) | 6.05 | 0.79 | 0.41 | 2.32 |
| | KM (µM) | 52.23 | 10.74 | 0.09 | 0.57 |
| | kcat/KM (M-1s-1) | 1.22E+05 | 4.24E+04 | 0.37 | 1.20 |
| SARS2-S123F | kcat (s-1) | 5.69 | 0.58 | 0.33 | 1.52 |
| | KM (µM) | 50.53 | 8.16 | 0.04 | 0.38 |
| | kcat/KM (M-1s-1) | 1.14E+05 | 8.99E+03 | 0.27 | 2.20 |
| SARS2-Y126F | kcat (s-1) | 5.91 | 0.76 | 0.38 | 1.72 |
| | KM (µM) | 48.65 | 9.60 | -0.01 | 0.32 |
| | kcat/KM (M-1s-1) | 1.23E+05 | 1.07E+04 | 0.38 | 3.49 |
| SARS2-Y126M | kcat (s-1) | 2.40 | 0.37 | -0.92 | 3.00 |
| | KM (µM) | 32.54 | 6.44 | -0.59 | 1.66 |
| | kcat/KM (M-1s-1) | 7.46E+04 | 7.17E+03 | -0.34 | 2.49 |
| SARS2-Q127H | kcat (s-1) | 5.08 | 0.61 | 0.16 | 0.80 |
| | KM (µM) | 49.30 | 7.38 | 0.01 | 0.33 |
| | kcat/KM (M-1s-1) | 1.04E+05 | 5.88E+03 | 0.13 | 1.42 |
| SARS2-A129V | kcat (s-1) | 5.32 | 1.19 | 0.23 | 1.01 |
| | KM (µM) | 49.02 | 12.46 | 0.00 | 0.30 |
| | kcat/KM (M-1s-1) | 1.09E+05 | 6.80E+03 | 0.21 | 2.82 |
| SARS2-P132A | kcat (s-1) | 2.91 | 0.36 | -0.64 | 2.34 |
| | KM (µM) | 39.72 | 8.35 | -0.30 | 0.97 |
| | kcat/KM (M-1s-1) | 7.46E+04 | 8.94E+03 | -0.34 | 3.00 |
| SARS2-P132H | kcat (s-1) | 2.87 | 0.72 | -0.66 | 2.36 |
| | KM (µM) | 38.45 | 12.26 | -0.35 | 1.29 |
| | kcat/KM (M-1s-1) | 7.75E+04 | 1.50E+04 | -0.29 | 1.74 |
| SARS2-P132L | kcat (s-1) | 2.93 | 0.39 | -0.63 | 2.31 |
| | KM (µM) | 42.46 | 5.73 | -0.21 | 0.84 |
| | kcat/KM (M-1s-1) | 6.97E+04 | 9.37E+03 | -0.44 | 2.94 |
| SARS2-P132R | kcat (s-1) | 2.40 | 0.30 | -0.92 | 3.00 |
| | KM (µM) | 34.88 | 6.65 | -0.49 | 1.51 |
| | kcat/KM (M-1s-1) | 6.96E+04 | 4.85E+03 | -0.44 | 3.72 |

| | | | | | |
|---|---|---|---|---|---|
| SARS2-P132S | kcat (s-1) | 3.44 | 0.58 | -0.40 | 1.91 |
| | KM (μM) | 36.89 | 9.74 | -0.41 | 1.28 |
| | kcat/KM (M-1s-1) | 9.66E+04 | 1.69E+04 | 0.03 | 0.32 |
| SARS2-P132T | kcat (s-1) | 3.07 | 0.36 | -0.56 | 2.45 |
| | KM (μM) | 37.02 | 3.97 | -0.41 | 1.62 |
| | kcat/KM (M-1s-1) | 8.33E+04 | 8.42E+03 | -0.18 | 1.67 |
| SARS2-F140M | kcat (s-1) | 5.06 | 0.84 | 0.16 | 0.81 |
| | KM (μM) | 54.25 | 12.05 | 0.15 | 0.69 |
| | kcat/KM (M-1s-1) | 9.44E+04 | 9.52E+03 | 0.00 | 0.32 |
| SARS2-L141A | kcat (s-1) | 2.01 | 0.66 | -1.18 | 3.10 |
| | KM (μM) | 78.08 | 12.98 | 0.67 | 3.36 |
| | kcat/KM (M-1s-1) | 2.60E+04 | 9.12E+03 | -1.86 | 5.72 |
| SARS2-L141E | kcat (s-1) | 1.53 | 0.69 | -1.57 | 3.69 |
| | KM (μM) | 50.33 | 11.23 | 0.04 | 0.36 |
| | kcat/KM (M-1s-1) | 2.97E+04 | 6.60E+03 | -1.67 | 5.71 |
| SARS2-L141H | kcat (s-1) | 2.33 | 0.83 | -0.96 | 2.25 |
| | KM (μM) | 70.71 | 14.95 | 0.53 | 1.73 |
| | kcat/KM (M-1s-1) | 3.35E+04 | 1.02E+04 | -1.50 | 7.49 |
| SARS2-L141I | kcat (s-1) | 1.39 | 0.20 | -1.71 | 4.12 |
| | KM (μM) | 50.99 | 9.80 | 0.06 | 0.41 |
| | kcat/KM (M-1s-1) | 2.77E+04 | 4.45E+03 | -1.77 | 7.82 |
| SARS2-L141K | kcat (s-1) | 2.97 | 0.66 | -0.61 | 1.87 |
| | KM (μM) | 96.83 | 28.70 | 0.98 | 2.24 |
| | kcat/KM (M-1s-1) | 3.14E+04 | 3.46E+03 | -1.59 | 6.09 |
| SARS2-L141M | kcat (s-1) | 3.87 | 0.85 | -0.23 | 1.69 |
| | KM (μM) | 51.72 | 16.53 | 0.08 | 0.35 |
| | kcat/KM (M-1s-1) | 7.75E+04 | 1.15E+04 | -0.29 | 2.13 |
| SARS2-L141N | kcat (s-1) | 1.92 | 0.31 | -1.24 | 3.86 |
| | KM (μM) | 71.20 | 10.08 | 0.54 | 2.81 |
| | kcat/KM (M-1s-1) | 2.74E+04 | 5.05E+03 | -1.79 | 6.67 |
| SARS2-L141Q | kcat (s-1) | 2.16 | 0.65 | -1.07 | 3.24 |
| | KM (μM) | 81.64 | 19.50 | 0.74 | 2.74 |
| | kcat/KM (M-1s-1) | 2.69E+04 | 5.64E+03 | -1.82 | 6.51 |

| | | | | | |
|---|---|---|---|---|---|
| SARS2-L141R | kcat (s-1) | 3.41 | 0.38 | -0.41 | 2.37 |
| | KM (μM) | 53.71 | 8.81 | 0.13 | 0.67 |
| | kcat/KM (M-1s-1) | 6.59E+04 | 1.74E+04 | -0.52 | 2.18 |
| SARS2-L141S | kcat (s-1) | 1.67 | 0.28 | -1.44 | 4.02 |
| | KM (μM) | 74.50 | 17.12 | 0.60 | 1.90 |
| | kcat/KM (M-1s-1) | 2.39E+04 | 7.37E+03 | -1.98 | 5.57 |
| SARS2-L141T | kcat (s-1) | 1.99 | 0.29 | -1.19 | 3.99 |
| | KM (μM) | 59.78 | 11.07 | 0.29 | 1.57 |
| | kcat/KM (M-1s-1) | 3.36E+04 | 2.94E+03 | -1.49 | 6.81 |
| SARS2-N142A | kcat (s-1) | 4.16 | 0.90 | -0.12 | 0.85 |
| | KM (μM) | 26.18 | 7.62 | -0.91 | 2.85 |
| | kcat/KM (M-1s-1) | 1.63E+05 | 2.60E+04 | 0.78 | 4.74 |
| SARS2-N142L | kcat (s-1) | 2.48 | 0.40 | -0.87 | 3.59 |
| | KM (μM) | 28.06 | 5.09 | -0.81 | 2.66 |
| | kcat/KM (M-1s-1) | 8.95E+04 | 1.13E+04 | -0.08 | 0.77 |
| SARS2-N142M | kcat (s-1) | 1.72 | 0.17 | -1.40 | 4.13 |
| | KM (μM) | 10.32 | 1.77 | -2.25 | 4.20 |
| | kcat/KM (M-1s-1) | 1.69E+05 | 2.20E+04 | 0.84 | 5.38 |
| SARS2-N142P | kcat (s-1) | 3.59 | 0.42 | -0.34 | 1.54 |
| | KM (μM) | 61.42 | 9.54 | 0.32 | 1.58 |
| | kcat/KM (M-1s-1) | 5.89E+04 | 5.85E+03 | -0.68 | 4.83 |
| SARS2-N142S | kcat (s-1) | 4.82 | 0.52 | 0.09 | 0.60 |
| | KM (μM) | 41.73 | 8.46 | -0.23 | 0.98 |
| | kcat/KM (M-1s-1) | 1.17E+05 | 1.21E+04 | 0.31 | 2.82 |
| SARS2-G143A | kcat (s-1) | 0.26 | 0.11 | -4.14 | 5.19 |
| | KM (μM) | 84.67 | 34.94 | 0.79 | 1.54 |
| | kcat/KM (M-1s-1) | 3.25E+03 | 1.33E+03 | -4.86 | 7.48 |
| SARS2-G143C | kcat (s-1) | Not Active | -- | -- | -- |
| | KM (μM) | Not Active | -- | -- | -- |
| | kcat/KM (M-1s-1) | Not Active | -- | -- | -- |
| SARS2-G143D | kcat (s-1) | 0.44 | 1.05 | -3.37 | 3.62 |
| | KM (μM) | 296.72 | 834.32 | 2.60 | 0.74 |
| | kcat/KM (M-1s-1) | 3.08E+03 | 1.21E+03 | -4.94 | 7.66 |

| | | | | | |
|---|---|---|---|---|---|
| SARS2-G143R | kcat (s-1) | 0.12 | 0.02 | -5.29 | 2.87 |
| | KM (μM) | 97.40 | 45.76 | 0.99 | 1.55 |
| | kcat/KM (M-1s-1) | 1.48E+03 | 8.56E+02 | -5.99 | 3.65 |
| SARS2-G143S | kcat (s-1) | 0.23 | 0.08 | -4.32 | 5.44 |
| | KM (μM) | 99.82 | 41.86 | 1.03 | 2.15 |
| | kcat/KM (M-1s-1) | 2.34E+03 | 2.89E+02 | -5.34 | 7.71 |
| SARS2-G143V | kcat (s-1) | Not Active | -- | -- | -- |
| | KM (μM) | Not Active | -- | -- | -- |
| | kcat/KM (M-1s-1) | Not Active | -- | -- | -- |
| SARS2-S144A | kcat (s-1) | 4.15 | 0.71 | -0.13 | 1.18 |
| | KM (μM) | 133.44 | 29.92 | 1.44 | 5.01 |
| | kcat/KM (M-1s-1) | 3.15E+04 | 2.47E+03 | -1.59 | 6.55 |
| SARS2-S144F | kcat (s-1) | 0.24 | 0.07 | -4.23 | 5.44 |
| | KM (μM) | 72.59 | 32.13 | 0.57 | 1.61 |
| | kcat/KM (M-1s-1) | 3.49E+03 | 5.95E+02 | -4.76 | 7.75 |
| SARS2-S144G | kcat (s-1) | 2.92 | 0.93 | -0.63 | 2.44 |
| | KM (μM) | 188.46 | 77.07 | 1.94 | 3.00 |
| | kcat/KM (M-1s-1) | 1.61E+04 | 1.84E+03 | -2.56 | 7.02 |
| SARS2-S144M | kcat (s-1) | 0.89 | 0.15 | -2.36 | 5.02 |
| | KM (μM) | 184.76 | 39.20 | 1.91 | 4.91 |
| | kcat/KM (M-1s-1) | 4.84E+03 | 4.56E+02 | -4.29 | 7.55 |
| SARS2-S144Y | kcat (s-1) | 0.22 | 0.06 | -4.39 | 5.49 |
| | KM (μM) | 116.47 | 50.38 | 1.25 | 2.40 |
| | kcat/KM (M-1s-1) | 2.07E+03 | 6.72E+02 | -5.51 | 7.78 |
| SARS2-N151C | kcat (s-1) | 5.27 | 1.05 | 0.22 | 0.83 |
| | KM (μM) | 51.02 | 10.82 | 0.06 | 0.39 |
| | kcat/KM (M-1s-1) | 1.04E+05 | 6.71E+03 | 0.13 | 1.38 |
| SARS2-N151F | kcat (s-1) | 6.24 | 0.76 | 0.46 | 2.03 |
| | KM (μM) | 55.67 | 11.07 | 0.18 | 0.63 |
| | kcat/KM (M-1s-1) | 1.14E+05 | 1.03E+04 | 0.27 | 2.10 |
| SARS2-N151G | kcat (s-1) | 3.94 | 1.48 | -0.20 | 0.59 |
| | KM (μM) | 54.32 | 11.71 | 0.15 | 0.48 |
| | kcat/KM (M-1s-1) | 7.19E+04 | 1.56E+04 | -0.40 | 1.97 |

| | | | | | |
|---|---|---|---|---|---|
| SARS2-N151I | kcat (s-1) | 6.68 | 1.77 | 0.56 | 1.91 |
| | KM (μM) | 57.55 | 14.05 | 0.23 | 0.98 |
| | kcat/KM (M-1s-1) | 1.16E+05 | 1.17E+04 | 0.30 | 3.39 |
| SARS2-N151M | kcat (s-1) | 6.10 | 1.60 | 0.43 | 1.73 |
| | KM (μM) | 48.29 | 9.12 | -0.02 | 0.34 |
| | kcat/KM (M-1s-1) | 1.28E+05 | 3.28E+04 | 0.44 | 1.72 |
| SARS2-N151R | kcat (s-1) | 4.89 | 1.27 | 0.11 | 0.32 |
| | KM (μM) | 56.58 | 15.25 | 0.21 | 0.45 |
| | kcat/KM (M-1s-1) | 8.67E+04 | 3.87E+03 | -0.13 | 1.64 |
| SARS2-N151S | kcat (s-1) | 4.18 | 0.52 | -0.12 | 0.94 |
| | KM (μM) | 45.01 | 8.32 | -0.12 | 0.66 |
| | kcat/KM (M-1s-1) | 9.44E+04 | 1.13E+04 | 0.00 | 0.31 |
| SARS2-N151T | kcat (s-1) | 7.46 | 1.04 | 0.72 | 3.15 |
| | KM (μM) | 59.54 | 9.74 | 0.28 | 1.00 |
| | kcat/KM (M-1s-1) | 1.26E+05 | 1.10E+04 | 0.41 | 4.00 |
| SARS2-N151W | kcat (s-1) | 5.82 | 0.62 | 0.36 | 1.69 |
| | KM (μM) | 56.13 | 11.73 | 0.19 | 0.56 |
| | kcat/KM (M-1s-1) | 1.05E+05 | 1.14E+04 | 0.16 | 1.71 |
| SARS2-N151Y | kcat (s-1) | 7.74 | 2.54 | 0.77 | 1.67 |
| | KM (μM) | 54.26 | 11.50 | 0.15 | 0.50 |
| | kcat/KM (M-1s-1) | 1.41E+05 | 2.37E+04 | 0.58 | 3.62 |
| SARS2-V157L | kcat (s-1) | 2.03 | 0.40 | -1.16 | 3.83 |
| | KM (μM) | 41.49 | 6.64 | -0.24 | 0.96 |
| | kcat/KM (M-1s-1) | 4.90E+04 | 5.98E+03 | -0.95 | 4.50 |
| SARS2-C160F | kcat (s-1) | 0.19 | 0.09 | -4.55 | 5.37 |
| | KM (μM) | 32.65 | 4.09 | -0.59 | 2.13 |
| | kcat/KM (M-1s-1) | 5.93E+03 | 2.54E+03 | -3.99 | 7.49 |
| SARS2-C160N | kcat (s-1) | 1.77 | 0.22 | -1.35 | 3.96 |
| | KM (μM) | 48.96 | 7.32 | 0.00 | 0.31 |
| | kcat/KM (M-1s-1) | 3.64E+04 | 2.78E+03 | -1.38 | 6.41 |
| SARS2-C160Y | kcat (s-1) | 0.34 | 0.09 | -3.75 | 5.34 |
| | KM (μM) | 44.73 | 16.02 | -0.13 | 0.52 |
| | kcat/KM (M-1s-1) | 8.21E+03 | 2.40E+03 | -3.53 | 7.19 |

| | | | | | |
|---|---|---|---|---|---|
| SARS2-M162I | kcat (s-1) | 1.00 | 0.24 | -2.19 | 5.28 |
| | KM (µM) | 34.68 | 8.84 | -0.50 | 2.27 |
| | kcat/KM (M-1s-1) | 2.90E+04 | 3.37E+03 | -1.71 | 6.50 |
| SARS2-H164N | kcat (s-1) | 0.78 | 0.12 | -2.53 | 5.08 |
| | KM (µM) | 101.20 | 25.80 | 1.05 | 3.79 |
| | kcat/KM (M-1s-1) | 7.95E+03 | 9.81E+02 | -3.57 | 7.62 |
| SARS2-H164Q | kcat (s-1) | 3.47 | 0.99 | -0.39 | 1.15 |
| | KM (µM) | 87.91 | 23.13 | 0.84 | 2.05 |
| | kcat/KM (M-1s-1) | 3.98E+04 | 6.10E+03 | -1.25 | 5.12 |
| SARS2-M165I | kcat (s-1) | 3.37 | 0.63 | -0.43 | 1.92 |
| | KM (µM) | 84.15 | 19.23 | 0.78 | 2.58 |
| | kcat/KM (M-1s-1) | 4.06E+04 | 3.53E+03 | -1.22 | 5.39 |
| SARS2-M165K | kcat (s-1) | Not Active | -- | -- | -- |
| | KM (µM) | Not Active | -- | -- | -- |
| | kcat/KM (M-1s-1) | Not Active | -- | -- | -- |
| SARS2-M165L | kcat (s-1) | 1.29 | 0.29 | -1.81 | 3.97 |
| | KM (µM) | 40.99 | 8.10 | -0.26 | 0.91 |
| | kcat/KM (M-1s-1) | 3.22E+04 | 7.19E+03 | -1.55 | 8.01 |
| SARS2-M165R | kcat (s-1) | Not Active | -- | -- | -- |
| | KM (µM) | Not Active | -- | -- | -- |
| | kcat/KM (M-1s-1) | Not Active | -- | -- | -- |
| SARS2-M165T | kcat (s-1) | 0.09 | 0.02 | -5.62 | 1.52 |
| | KM (µM) | 62.95 | 37.99 | 0.36 | 0.41 |
| | kcat/KM (M-1s-1) | 1.65E+03 | 5.97E+02 | -5.84 | 3.34 |
| SARS2-M165V | kcat (s-1) | 2.59 | 0.46 | -0.81 | 3.17 |
| | KM (µM) | 66.41 | 13.89 | 0.44 | 1.68 |
| | kcat/KM (M-1s-1) | 3.94E+04 | 3.46E+03 | -1.26 | 5.85 |
| SARS2-E166A | kcat (s-1) | 1.90 | 0.62 | -1.26 | 4.09 |
| | KM (µM) | 107.13 | 32.80 | 1.13 | 2.92 |
| | kcat/KM (M-1s-1) | 1.79E+04 | 2.90E+03 | -2.40 | 7.01 |
| SARS2-E166D | kcat (s-1) | 0.07 | 0.04 | -5.98 | 1.32 |
| | KM (µM) | 52.35 | 33.36 | 0.09 | 0.36 |
| | kcat/KM (M-1s-1) | 1.41E+03 | 1.07E+02 | -6.07 | 1.58 |

| | | | | | |
|---|---|---|---|---|---|
| SARS2-E166G | kcat (s-1) | 0.74 | 0.22 | -2.62 | 5.23 |
| | KM (μM) | 97.38 | 45.67 | 0.99 | 1.82 |
| | kcat/KM (M-1s-1) | 8.42E+03 | 2.47E+03 | -3.49 | 7.36 |
| SARS2-E166K | kcat (s-1) | 0.54 | 0.15 | -3.07 | 5.08 |
| | KM (μM) | 77.62 | 27.59 | 0.66 | 1.43 |
| | kcat/KM (M-1s-1) | 7.29E+03 | 1.43E+03 | -3.70 | 7.35 |
| SARS2-E166M | kcat (s-1) | 1.31 | 0.54 | -1.79 | 3.98 |
| | KM (μM) | 104.60 | 53.24 | 1.09 | 1.80 |
| | kcat/KM (M-1s-1) | 1.31E+04 | 2.29E+03 | -2.85 | 7.67 |
| SARS2-E166Q | kcat (s-1) | 3.16 | 0.95 | -0.52 | 2.00 |
| | KM (μM) | 38.01 | 14.49 | -0.37 | 1.10 |
| | kcat/KM (M-1s-1) | 8.52E+04 | 9.32E+03 | -0.15 | 1.27 |
| SARS2-E166V | kcat (s-1) | 0.65 | 0.23 | -2.80 | 4.89 |
| | KM (μM) | 129.19 | 52.43 | 1.40 | 2.28 |
| | kcat/KM (M-1s-1) | 5.27E+03 | 8.81E+02 | -4.17 | 7.42 |
| SARS2-L50F/E166V | kcat (s-1) | 1.35 | 0.33 | -1.75 | 4.15 |
| | KM (μM) | 109.08 | 29.66 | 1.15 | 2.96 |
| | kcat/KM (M-1s-1) | 1.25E+04 | 1.05E+03 | -2.92 | 7.45 |
| SARS2-T21I/E166V | kcat (s-1) | 0.31 | 0.07 | -3.86 | 5.40 |
| | KM (μM) | 107.39 | 94.72 | 1.13 | 1.71 |
| | kcat/KM (M-1s-1) | 3.74E+03 | 1.13E+03 | -4.66 | 7.67 |
| SARS2-P168Δ | kcat (s-1) | 0.66 | 0.15 | -2.77 | 5.11 |
| | KM (μM) | 46.41 | 10.46 | -0.08 | 0.47 |
| | kcat/KM (M-1s-1) | 1.47E+04 | 3.29E+03 | -2.69 | 7.42 |
| SARS2-T169S | kcat (s-1) | 4.97 | 1.17 | 0.13 | 0.79 |
| | KM (μM) | 51.01 | 11.09 | 0.06 | 0.46 |
| | kcat/KM (M-1s-1) | 9.73E+04 | 4.23E+03 | 0.04 | 0.65 |
| SARS2-T169P | kcat (s-1) | 4.22 | 0.77 | -0.10 | 0.65 |
| | KM (μM) | 45.18 | 10.94 | -0.12 | 0.68 |
| | kcat/KM (M-1s-1) | 9.49E+04 | 8.42E+03 | 0.01 | 0.33 |
| SARS2-G170R | kcat (s-1) | 3.46 | 0.48 | -0.39 | 2.22 |
| | KM (μM) | 37.26 | 6.35 | -0.40 | 1.66 |
| | kcat/KM (M-1s-1) | 9.34E+04 | 8.13E+03 | -0.02 | 0.37 |

| | | | | | |
|---|---|---|---|---|---|
| SARS2-H172F | kcat (s-1) | 0.09 | 0.04 | -5.65 | 3.18 |
| | KM (μM) | 97.09 | 62.42 | 0.99 | 0.90 |
| | kcat/KM (M-1s-1) | 1.12E+03 | 4.22E+02 | -6.39 | 3.72 |
| SARS2-H172Q | kcat (s-1) | 0.37 | 0.32 | -3.61 | 5.39 |
| | KM (μM) | 167.50 | 151.70 | 1.77 | 1.54 |
| | kcat/KM (M-1s-1) | 2.50E+03 | 7.19E+02 | -5.24 | 7.59 |
| SARS2-A173D | kcat (s-1) | Not Active | -- | -- | -- |
| | KM (μM) | Not Active | -- | -- | -- |
| | kcat/KM (M-1s-1) | Not Active | -- | -- | -- |
| SARS2-A173G | kcat (s-1) | 0.81 | 0.26 | -2.49 | 4.74 |
| | KM (μM) | 42.16 | 12.88 | -0.22 | 0.76 |
| | kcat/KM (M-1s-1) | 1.96E+04 | 5.29E+03 | -2.27 | 6.18 |
| SARS2-A173P | kcat (s-1) | Not Active | -- | -- | -- |
| | KM (μM) | Not Active | -- | -- | -- |
| | kcat/KM (M-1s-1) | Not Active | -- | -- | -- |
| SARS2-A173S | kcat (s-1) | 2.23 | 0.45 | -1.03 | 3.32 |
| | KM (μM) | 46.60 | 9.77 | -0.07 | 0.33 |
| | kcat/KM (M-1s-1) | 4.82E+04 | 6.41E+03 | -0.97 | 5.63 |
| SARS2-A173T | kcat (s-1) | 2.82 | 0.61 | -0.68 | 2.58 |
| | KM (μM) | 83.11 | 31.94 | 0.76 | 1.97 |
| | kcat/KM (M-1s-1) | 3.63E+04 | 7.85E+03 | -1.38 | 5.51 |
| SARS2-A173V | kcat (s-1) | 3.55 | 0.97 | -0.35 | 1.20 |
| | KM (μM) | 166.50 | 45.74 | 1.76 | 3.78 |
| | kcat/KM (M-1s-1) | 2.15E+04 | 1.51E+03 | -2.14 | 6.71 |
| SARS2-N180D | kcat (s-1) | 2.81 | 0.51 | -0.69 | 2.74 |
| | KM (μM) | 44.78 | 11.25 | -0.13 | 0.55 |
| | kcat/KM (M-1s-1) | 6.38E+04 | 6.85E+03 | -0.57 | 3.38 |
| SARS2-P184S | kcat (s-1) | 3.74 | 0.92 | -0.28 | 1.53 |
| | KM (μM) | 43.16 | 11.76 | -0.18 | 1.09 |
| | kcat/KM (M-1s-1) | 8.80E+04 | 9.62E+03 | -0.10 | 0.64 |
| SARS2-F185S | kcat (s-1) | Not Active | -- | -- | -- |
| | KM (μM) | Not Active | -- | -- | -- |
| | kcat/KM (M-1s-1) | Not Active | -- | -- | -- |

| | | | | | |
|---|---|---|---|---|---|
| SARS2-V186F | kcat (s-1) | 4.29 | 0.73 | -0.08 | 0.49 |
| | KM (μM) | 43.73 | 11.59 | -0.17 | 0.59 |
| | kcat/KM (M-1s-1) | 1.00E+05 | 9.55E+03 | 0.08 | 0.74 |
| SARS2-R188K | kcat (s-1) | 4.98 | 0.57 | 0.14 | 0.75 |
| | KM (μM) | 49.15 | 10.13 | 0.00 | 0.40 |
| | kcat/KM (M-1s-1) | 1.04E+05 | 1.70E+04 | 0.14 | 1.30 |
| SARS2-Q189Δ | kcat (s-1) | Not Active | -- | -- | -- |
| | KM (μM) | Not Active | -- | -- | -- |
| | kcat/KM (M-1s-1) | Not Active | -- | -- | -- |
| SARS2-Q189E | kcat (s-1) | 4.08 | 0.61 | -0.15 | 1.05 |
| | KM (μM) | 29.25 | 4.42 | -0.75 | 2.92 |
| | kcat/KM (M-1s-1) | 1.40E+05 | 1.32E+04 | 0.57 | 4.11 |
| SARS2-Q189F | kcat (s-1) | 1.02 | 0.28 | -2.15 | 4.96 |
| | KM (μM) | 128.48 | 39.50 | 1.39 | 3.40 |
| | kcat/KM (M-1s-1) | 8.06E+03 | 7.71E+02 | -3.55 | 7.49 |
| SARS2-Q189G | kcat (s-1) | 2.56 | 0.83 | -0.83 | 2.67 |
| | KM (μM) | 114.47 | 48.88 | 1.22 | 2.26 |
| | kcat/KM (M-1s-1) | 2.35E+04 | 3.62E+03 | -2.01 | 6.37 |
| SARS2-Q189H | kcat (s-1) | 1.96 | 0.31 | -1.21 | 3.78 |
| | KM (μM) | 84.50 | 22.66 | 0.78 | 2.35 |
| | kcat/KM (M-1s-1) | 2.42E+04 | 4.78E+03 | -1.97 | 6.34 |
| SARS2-Q189K | kcat (s-1) | 1.51 | 0.60 | -1.59 | 3.89 |
| | KM (μM) | 157.94 | 63.89 | 1.69 | 2.79 |
| | kcat/KM (M-1s-1) | 9.78E+03 | 1.71E+03 | -3.27 | 7.04 |
| SARS2-Q189L | kcat (s-1) | 0.89 | 0.15 | -2.36 | 5.04 |
| | KM (μM) | 126.55 | 28.60 | 1.37 | 5.55 |
| | kcat/KM (M-1s-1) | 7.10E+03 | 7.75E+02 | -3.73 | 7.60 |
| SARS2-Q189P | kcat (s-1) | 2.02 | 0.44 | -1.17 | 3.79 |
| | KM (μM) | 66.86 | 12.91 | 0.45 | 1.80 |
| | kcat/KM (M-1s-1) | 3.02E+04 | 2.45E+03 | -1.65 | 6.10 |
| SARS2-Q189R | kcat (s-1) | 1.61 | 0.39 | -1.49 | 4.57 |
| | KM (μM) | 108.11 | 34.52 | 1.14 | 3.18 |
| | kcat/KM (M-1s-1) | 1.52E+04 | 1.56E+03 | -2.63 | 7.18 |

| | | | | | |
|---|---|---|---|---|---|
| SARS2-Q189S | kcat (s-1) | 3.01 | 0.56 | -0.59 | 1.99 |
| | KM (μM) | 128.43 | 29.15 | 1.39 | 3.50 |
| | kcat/KM (M-1s-1) | 2.36E+04 | 2.01E+03 | -2.00 | 6.77 |
| SARS2-T190I | kcat (s-1) | 5.23 | 1.04 | 0.21 | 0.73 |
| | KM (μM) | 47.59 | 10.69 | -0.04 | 0.50 |
| | kcat/KM (M-1s-1) | 1.12E+05 | 1.88E+04 | 0.24 | 2.44 |
| SARS2-A191V | kcat (s-1) | 4.69 | 1.01 | 0.05 | 0.31 |
| | KM (μM) | 38.04 | 11.11 | -0.37 | 1.31 |
| | kcat/KM (M-1s-1) | 1.27E+05 | 2.01E+04 | 0.43 | 2.60 |
| SARS2-Q192T | kcat (s-1) | 0.41 | 0.15 | -3.45 | 5.42 |
| | KM (μM) | 103.37 | 48.02 | 1.08 | 2.32 |
| | kcat/KM (M-1s-1) | 4.17E+03 | 4.71E+02 | -4.50 | 7.67 |
| SARS2-Q192A | kcat (s-1) | 0.39 | 0.17 | -3.56 | 5.02 |
| | KM (μM) | 123.85 | 103.35 | 1.34 | 1.29 |
| | kcat/KM (M-1s-1) | 3.80E+03 | 9.93E+02 | -4.64 | 7.49 |
| SARS2-Q192C | kcat (s-1) | 0.56 | 0.21 | -3.03 | 5.32 |
| | KM (μM) | 96.12 | 32.94 | 0.97 | 3.05 |
| | kcat/KM (M-1s-1) | 5.91E+03 | 1.29E+03 | -4.00 | 7.55 |
| SARS2-Q192F | kcat (s-1) | 1.04 | 0.36 | -2.13 | 4.59 |
| | KM (μM) | 101.55 | 41.44 | 1.05 | 2.09 |
| | kcat/KM (M-1s-1) | 1.05E+04 | 1.10E+03 | -3.17 | 7.22 |
| SARS2-Q192H | kcat (s-1) | 0.11 | 0.03 | -5.31 | 5.51 |
| | KM (μM) | 51.46 | 40.49 | 0.07 | 0.37 |
| | kcat/KM (M-1s-1) | 2.83E+03 | 9.33E+02 | -5.06 | 7.69 |
| SARS2-Q192I | kcat (s-1) | Not Active | -- | -- | -- |
| | KM (μM) | Not Active | -- | -- | -- |
| | kcat/KM (M-1s-1) | Not Active | -- | -- | -- |
| SARS2-Q192L | kcat (s-1) | 0.82 | 0.34 | -2.47 | 4.89 |
| | KM (μM) | 142.32 | 73.67 | 1.54 | 2.19 |
| | kcat/KM (M-1s-1) | 6.37E+03 | 1.55E+03 | -3.89 | 7.34 |
| SARS2-Q192P | kcat (s-1) | 0.26 | 0.12 | -4.11 | 5.57 |
| | KM (μM) | 92.37 | 50.56 | 0.91 | 1.87 |
| | kcat/KM (M-1s-1) | 3.20E+03 | 1.03E+03 | -4.88 | 7.85 |

| | | | | | |
|---|---|---|---|---|---|
| SARS2-Q192S | kcat (s-1) | 0.15 | 0.12 | -4.93 | 5.33 |
| | KM (μM) | 58.31 | 50.10 | 0.25 | 0.56 |
| | kcat/KM (M-1s-1) | 2.66E+03 | 6.05E+02 | -5.15 | 7.62 |
| SARS2-Q192V | kcat (s-1) | 0.71 | 0.27 | -2.68 | 5.14 |
| | KM (μM) | 136.84 | 70.39 | 1.48 | 3.00 |
| | kcat/KM (M-1s-1) | 5.59E+03 | 1.16E+03 | -4.08 | 7.63 |
| SARS2-Q192W | kcat (s-1) | 0.48 | 0.16 | -3.24 | 5.27 |
| | KM (μM) | 102.07 | 48.82 | 1.06 | 2.21 |
| | kcat/KM (M-1s-1) | 5.28E+03 | 1.62E+03 | -4.16 | 7.51 |
| SARS2-T196M | kcat (s-1) | 4.43 | 1.13 | -0.03 | 0.36 |
| | KM (μM) | 37.59 | 10.38 | -0.38 | 1.17 |
| | kcat/KM (M-1s-1) | 1.19E+05 | 1.16E+04 | 0.33 | 2.49 |
| SARS2-V202W | kcat (s-1) | 1.88 | 0.31 | -1.27 | 3.97 |
| | KM (μM) | 44.19 | 11.99 | -0.15 | 0.63 |
| | kcat/KM (M-1s-1) | 4.34E+04 | 4.70E+03 | -1.12 | 6.06 |
| SARS2-L205V | kcat (s-1) | 2.86 | 0.58 | -0.67 | 2.28 |
| | KM (μM) | 39.88 | 10.80 | -0.30 | 0.95 |
| | kcat/KM (M-1s-1) | 7.32E+04 | 9.62E+03 | -0.37 | 3.08 |
| SARS2-A206V | kcat (s-1) | Not Active | -- | -- | -- |
| | KM (μM) | Not Active | -- | -- | -- |
| | kcat/KM (M-1s-1) | Not Active | -- | -- | -- |
| SARS2-V212F | kcat (s-1) | 2.83 | 0.19 | -0.68 | 2.95 |
| | KM (μM) | 35.62 | 2.21 | -0.46 | 1.86 |
| | kcat/KM (M-1s-1) | 7.95E+04 | 4.53E+03 | -0.25 | 2.82 |
| SARS2-I213V | kcat (s-1) | 2.91 | 0.75 | -0.64 | 3.71 |
| | KM (μM) | 42.48 | 13.20 | -0.21 | 1.21 |
| | kcat/KM (M-1s-1) | 6.94E+04 | 5.03E+03 | -0.45 | 4.38 |
| SARS2-L220F | kcat (s-1) | 2.14 | 0.56 | -1.09 | 3.41 |
| | KM (μM) | 37.33 | 8.86 | -0.39 | 1.50 |
| | kcat/KM (M-1s-1) | 5.79E+04 | 1.11E+04 | -0.71 | 3.41 |
| SARS2-F223L | kcat (s-1) | 4.88 | 0.93 | 0.11 | 0.57 |
| | KM (μM) | 46.73 | 12.86 | -0.07 | 0.42 |
| | kcat/KM (M-1s-1) | 1.06E+05 | 8.78E+03 | 0.17 | 1.64 |

| | | | | | |
|---|---|---|---|---|---|
| SARS2-Y239A | kcat (s-1) | 3.96 | 1.23 | -0.20 | 0.70 |
| | KM (µM) | 42.02 | 8.58 | -0.22 | 0.81 |
| | kcat/KM (M-1s-1) | 9.42E+04 | 1.70E+04 | -0.01 | 0.32 |
| SARS2-Y239G | kcat (s-1) | 0.51 | 0.22 | -3.15 | 4.73 |
| | KM (µM) | 36.62 | 14.00 | -0.42 | 1.08 |
| | kcat/KM (M-1s-1) | 1.44E+04 | 3.63E+03 | -2.71 | 7.24 |
| SARS2-E240M | kcat (s-1) | 4.52 | 1.83 | -0.01 | 0.31 |
| | KM (µM) | 43.29 | 12.17 | -0.18 | 0.60 |
| | kcat/KM (M-1s-1) | 1.03E+05 | 1.32E+04 | 0.12 | 2.79 |
| SARS2-P241L | kcat (s-1) | 3.19 | 0.55 | -0.51 | 2.10 |
| | KM (µM) | 37.27 | 10.42 | -0.40 | 1.44 |
| | kcat/KM (M-1s-1) | 8.86E+04 | 1.38E+04 | -0.09 | 0.73 |
| SARS2-H246Y | kcat (s-1) | 1.28 | 0.17 | -1.83 | 4.69 |
| | KM (µM) | 36.83 | 7.31 | -0.41 | 1.42 |
| | kcat/KM (M-1s-1) | 3.54E+04 | 5.62E+03 | -1.42 | 5.38 |
| SARS2-D248F | kcat (s-1) | 3.06 | 0.70 | -0.57 | 1.83 |
| | KM (µM) | 35.02 | 12.01 | -0.49 | 1.24 |
| | kcat/KM (M-1s-1) | 9.02E+04 | 1.09E+04 | -0.07 | 0.61 |
| SARS2-I259L | kcat (s-1) | 2.04 | 0.47 | -1.15 | 3.98 |
| | KM (µM) | 37.03 | 11.23 | -0.41 | 1.38 |
| | kcat/KM (M-1s-1) | 5.72E+04 | 1.12E+04 | -0.72 | 4.01 |
| SARS2-A260V | kcat (s-1) | 2.34 | 0.27 | -0.95 | 3.47 |
| | KM (µM) | 40.03 | 3.87 | -0.29 | 1.07 |
| | kcat/KM (M-1s-1) | 5.85E+04 | 4.53E+03 | -0.69 | 4.30 |
| SARS2-A266S | kcat (s-1) | 4.75 | 1.18 | 0.07 | 0.40 |
| | KM (µM) | 47.52 | 13.38 | -0.05 | 0.56 |
| | kcat/KM (M-1s-1) | 1.03E+05 | 2.02E+04 | 0.12 | 1.45 |
| SARS2-G278R | kcat (s-1) | 2.03 | 0.20 | -1.16 | 3.81 |
| | KM (µM) | 42.49 | 12.08 | -0.21 | 0.90 |
| | kcat/KM (M-1s-1) | 5.02E+04 | 1.06E+04 | -0.91 | 4.92 |
| SARS2-R279C | kcat (s-1) | 2.84 | 0.61 | -0.68 | 2.84 |
| | KM (µM) | 45.77 | 13.04 | -0.10 | 0.37 |
| | kcat/KM (M-1s-1) | 6.42E+04 | 1.32E+04 | -0.56 | 3.81 |

| | | | | | |
|---|---|---|---|---|---|
| SARS2-G283S | kcat (s-1) | 2.75 | 0.55 | -0.72 | 3.37 |
| | KM (μM) | 44.74 | 10.49 | -0.13 | 0.72 |
| | kcat/KM (M-1s-1) | 6.23E+04 | 6.87E+03 | -0.60 | 4.12 |
| SARS2-S284G | kcat (s-1) | 2.54 | 0.29 | -0.84 | 3.13 |
| | KM (μM) | 40.44 | 2.91 | -0.28 | 1.22 |
| | kcat/KM (M-1s-1) | 6.27E+04 | 4.71E+03 | -0.59 | 5.90 |
| SARS2-A285T | kcat (s-1) | 4.04 | 0.58 | -0.17 | 1.61 |
| | KM (μM) | 45.93 | 7.88 | -0.09 | 0.78 |
| | kcat/KM (M-1s-1) | 8.87E+04 | 8.37E+03 | -0.09 | 0.77 |
| SARS2-S301F | kcat (s-1) | 4.83 | 0.68 | 0.09 | 0.58 |
| | KM (μM) | 31.10 | 9.73 | -0.66 | 1.84 |
| | kcat/KM (M-1s-1) | 1.62E+05 | 2.93E+04 | 0.78 | 3.38 |

**Table S5. Michaelis-Menten parameters for the evolution-focused NL63 fusion sublibrary**

| MutantID | Measurement | ValueM | SD | log2FC | -log10(q-value) |
|---|---|---|---|---|---|
| SARS2-WT | kcat (s-1) | 3.73 | 1.12 | 0.00 | N/A |
| | KM (µM) | 78.23 | 20.39 | 0.00 | N/A |
| | kcat/KM (M-1s-1) | 4.87E+04 | 1.11E+04 | 0.00 | N/A |
| SARS2-C145A | kcat (s-1) | Not Active | -- | -- | -- |
| | KM (µM) | Not Active | -- | -- | -- |
| | kcat/KM (M-1s-1) | Not Active | -- | -- | -- |
| SARS2-TQA-WT | kcat (s-1) | 0.95 | 0.28 | -1.96 | 4.05 |
| | KM (µM) | 76.79 | 20.08 | -0.03 | 0.48 |
| | kcat/KM (M-1s-1) | 1.28E+04 | 3.18E+03 | -1.93 | 4.70 |
| SARS2-TQA-C145A | kcat (s-1) | Not Active | -- | -- | -- |
| | KM (µM) | Not Active | -- | -- | -- |
| | kcat/KM (M-1s-1) | Not Active | -- | -- | -- |
| SARS2-F003L | kcat (s-1) | 3.35 | 1.17 | -0.15 | 0.35 |
| | KM (µM) | 69.25 | 20.03 | -0.18 | 0.49 |
| | kcat/KM (M-1s-1) | 5.14E+04 | 1.98E+04 | 0.08 | 0.53 |
| SARS2-R004K | kcat (s-1) | 4.18 | 1.59 | 0.17 | 0.58 |
| | KM (µM) | 66.44 | 11.61 | -0.24 | 1.03 |
| | kcat/KM (M-1s-1) | 6.34E+04 | 2.15E+04 | 0.38 | 1.61 |
| SARS2-F008Q | kcat (s-1) | 3.35 | 0.62 | -0.15 | 0.96 |
| | KM (µM) | 88.57 | 18.97 | 0.18 | 0.81 |
| | kcat/KM (M-1s-1) | 3.88E+04 | 8.68E+03 | -0.33 | 1.28 |
| SARS2-K012C | kcat (s-1) | 2.57 | 0.28 | -0.54 | 1.84 |
| | KM (µM) | 66.41 | 11.28 | -0.24 | 0.89 |
| | kcat/KM (M-1s-1) | 3.92E+04 | 4.74E+03 | -0.31 | 1.54 |
| SARS2-G015R | kcat (s-1) | 3.09 | 1.12 | -0.27 | 1.51 |
| | KM (µM) | 64.60 | 13.04 | -0.28 | 1.23 |
| | kcat/KM (M-1s-1) | 4.95E+04 | 2.08E+04 | 0.02 | 0.30 |
| SARS2-M017V | kcat (s-1) | 3.25 | 0.74 | -0.20 | 0.74 |
| | KM (µM) | 65.29 | 9.94 | -0.26 | 1.27 |
| | kcat/KM (M-1s-1) | 5.08E+04 | 1.30E+04 | 0.06 | 0.44 |
| SARS2-Q019R | kcat (s-1) | 3.29 | 0.57 | -0.18 | 0.41 |
| | KM (µM) | 113.96 | 20.03 | 0.54 | 2.17 |
| | kcat/KM (M-1s-1) | 2.90E+04 | 2.55E+03 | -0.75 | 2.49 |
| SARS2-T021C | kcat (s-1) | 4.48 | 1.23 | 0.27 | 2.56 |
| | KM (µM) | 67.06 | 18.78 | -0.22 | 1.38 |

| | | | | | |
|---|---|---|---|---|---|
| | kcat/KM (M-1s-1) | 6.74E+04 | 5.78E+03 | 0.47 | 3.19 |
| SARS2-C022Y | kcat (s-1) | 1.80 | 0.61 | -1.05 | 2.53 |
| | KM (µM) | 81.01 | 23.01 | 0.05 | 0.48 |
| | kcat/KM (M-1s-1) | 2.31E+04 | 7.93E+03 | -1.07 | 2.53 |
| SARS2-T024S | kcat (s-1) | 3.46 | 0.99 | -0.11 | 0.52 |
| | KM (µM) | 87.44 | 19.72 | 0.16 | 0.75 |
| | kcat/KM (M-1s-1) | 4.13E+04 | 1.47E+04 | -0.24 | 0.71 |
| SARS2-T026V | kcat (s-1) | 3.66 | 0.76 | -0.02 | 0.35 |
| | KM (µM) | 62.01 | 10.65 | -0.34 | 1.72 |
| | kcat/KM (M-1s-1) | 5.97E+04 | 1.22E+04 | 0.29 | 1.78 |
| SARS2-L030V | kcat (s-1) | 3.06 | 0.78 | -0.28 | 0.93 |
| | KM (µM) | 59.09 | 8.69 | -0.40 | 1.71 |
| | kcat/KM (M-1s-1) | 5.23E+04 | 1.24E+04 | 0.10 | 0.52 |
| SARS2-D033G | kcat (s-1) | 3.33 | 0.84 | -0.16 | 0.87 |
| | KM (µM) | 73.73 | 15.30 | -0.09 | 0.63 |
| | kcat/KM (M-1s-1) | 4.52E+04 | 5.88E+03 | -0.11 | 0.90 |
| SARS2-V035T | kcat (s-1) | 3.19 | 0.67 | -0.23 | 0.65 |
| | KM (µM) | 68.58 | 15.29 | -0.19 | 0.53 |
| | kcat/KM (M-1s-1) | 4.86E+04 | 1.35E+04 | 0.00 | 0.36 |
| SARS2-Y037T | kcat (s-1) | 5.66 | 1.28 | 0.60 | 2.34 |
| | KM (µM) | 72.02 | 13.19 | -0.12 | 0.40 |
| | kcat/KM (M-1s-1) | 8.30E+04 | 3.46E+04 | 0.77 | 1.85 |
| SARS2-C044A | kcat (s-1) | 2.56 | 0.81 | -0.54 | 1.75 |
| | KM (µM) | 70.78 | 18.94 | -0.14 | 0.67 |
| | kcat/KM (M-1s-1) | 3.72E+04 | 1.03E+04 | -0.39 | 2.10 |
| SARS2-T045P | kcat (s-1) | 1.96 | 0.98 | -0.92 | 1.74 |
| | KM (µM) | 110.22 | 22.71 | 0.49 | 3.34 |
| | kcat/KM (M-1s-1) | 1.74E+04 | 6.35E+03 | -1.49 | 2.72 |
| SARS2-E047T | kcat (s-1) | 3.19 | 0.73 | -0.23 | 0.95 |
| | KM (µM) | 64.29 | 9.76 | -0.28 | 1.64 |
| | kcat/KM (M-1s-1) | 5.03E+04 | 1.13E+04 | 0.05 | 0.39 |
| SARS2-D048T | kcat (s-1) | 2.15 | 0.41 | -0.79 | 2.50 |
| | KM (µM) | 71.53 | 13.45 | -0.13 | 0.40 |
| | kcat/KM (M-1s-1) | 3.05E+04 | 5.12E+03 | -0.67 | 2.30 |
| SARS2-M049Δ | kcat (s-1) | 1.44 | 0.48 | -1.37 | 4.15 |
| | KM (µM) | 256.95 | 66.36 | 1.72 | 4.19 |
| | kcat/KM (M-1s-1) | 5.60E+03 | 9.90E+02 | -3.12 | 5.26 |
| SARS2-L050V | kcat (s-1) | 4.27 | 1.50 | 0.20 | 0.41 |
| | KM (µM) | 61.55 | 14.24 | -0.35 | 1.04 |

| | | | | | |
|---|---|---|---|---|---|
| | kcat/KM (M-1s-1) | 7.16E+04 | 2.58E+04 | 0.56 | 1.28 |
| SARS2-N051L | kcat (s-1) | 2.04 | 0.29 | -0.87 | 3.03 |
| | KM (μM) | 72.67 | 12.08 | -0.11 | 0.69 |
| | kcat/KM (M-1s-1) | 2.84E+04 | 4.24E+03 | -0.78 | 4.12 |
| SARS2-N051Δ | kcat (s-1) | 0.58 | 0.16 | -2.67 | 4.82 |
| | KM (μM) | 126.63 | 28.41 | 0.69 | 2.37 |
| | kcat/KM (M-1s-1) | 4.83E+03 | 1.58E+03 | -3.33 | 4.39 |
| SARS2-P052I | kcat (s-1) | 1.23 | 0.69 | -1.60 | 4.34 |
| | KM (μM) | 141.71 | 38.45 | 0.86 | 2.37 |
| | kcat/KM (M-1s-1) | 9.09E+03 | 4.78E+03 | -2.42 | 4.34 |
| SARS2-N053D | kcat (s-1) | 3.41 | 0.61 | -0.13 | 0.55 |
| | KM (μM) | 75.11 | 18.60 | -0.06 | 0.41 |
| | kcat/KM (M-1s-1) | 4.75E+04 | 1.11E+04 | -0.04 | 0.36 |
| SARS2-E055D | kcat (s-1) | 3.57 | 0.72 | -0.06 | 0.44 |
| | KM (μM) | 71.03 | 5.78 | -0.14 | 0.73 |
| | kcat/KM (M-1s-1) | 5.01E+04 | 7.87E+03 | 0.04 | 0.39 |
| SARS2-D056H | kcat (s-1) | 3.20 | 0.93 | -0.22 | 1.61 |
| | KM (μM) | 70.56 | 15.81 | -0.15 | 0.73 |
| | kcat/KM (M-1s-1) | 4.55E+04 | 7.69E+03 | -0.10 | 0.73 |
| SARS2-L057A | kcat (s-1) | 1.85 | 1.16 | -1.01 | 2.35 |
| | KM (μM) | 80.34 | 21.79 | 0.04 | 0.39 |
| | kcat/KM (M-1s-1) | 2.26E+04 | 1.02E+04 | -1.11 | 3.30 |
| SARS2-L058Y | kcat (s-1) | 2.09 | 0.43 | -0.84 | 2.72 |
| | KM (μM) | 100.55 | 16.85 | 0.36 | 1.87 |
| | kcat/KM (M-1s-1) | 2.10E+04 | 3.93E+03 | -1.22 | 3.77 |
| SARS2-I059S | kcat (s-1) | 4.27 | 1.42 | 0.20 | 1.05 |
| | KM (μM) | 79.12 | 28.49 | 0.02 | 0.41 |
| | kcat/KM (M-1s-1) | 5.53E+04 | 1.07E+04 | 0.18 | 0.84 |
| SARS2-R060T | kcat (s-1) | 3.50 | 0.70 | -0.09 | 0.41 |
| | KM (μM) | 75.76 | 18.93 | -0.05 | 0.31 |
| | kcat/KM (M-1s-1) | 4.80E+04 | 1.32E+04 | -0.02 | 0.33 |
| SARS2-K061M | kcat (s-1) | 4.25 | 0.96 | 0.19 | 0.69 |
| | KM (μM) | 80.96 | 22.81 | 0.05 | 0.38 |
| | kcat/KM (M-1s-1) | 5.44E+04 | 1.23E+04 | 0.16 | 0.59 |
| SARS2-S062R | kcat (s-1) | 3.91 | 1.14 | 0.07 | 0.58 |
| | KM (μM) | 69.31 | 9.01 | -0.17 | 0.86 |
| | kcat/KM (M-1s-1) | 5.81E+04 | 2.01E+04 | 0.25 | 1.25 |
| SARS2-N063L | kcat (s-1) | 2.34 | 0.50 | -0.67 | 1.93 |
| | KM (μM) | 66.55 | 11.18 | -0.23 | 0.65 |

| | | | | | |
|---|---|---|---|---|---|
| | kcat/KM (M-1s-1) | 3.63E+04 | 9.80E+03 | -0.43 | 1.15 |
| SARS2-L067S | kcat (s-1) | 3.91 | 0.86 | 0.07 | 0.50 |
| | KM (µM) | 80.70 | 19.43 | 0.04 | 0.41 |
| | kcat/KM (M-1s-1) | 4.92E+04 | 6.60E+03 | 0.01 | 0.34 |
| SARS2-Q069S | kcat (s-1) | 3.94 | 0.52 | 0.08 | 1.85 |
| | KM (µM) | 101.67 | 20.08 | 0.38 | 1.72 |
| | kcat/KM (M-1s-1) | 3.93E+04 | 4.56E+03 | -0.31 | 0.90 |
| SARS2-A070H | kcat (s-1) | 4.03 | 0.83 | 0.11 | 0.63 |
| | KM (µM) | 74.28 | 19.64 | -0.07 | 0.45 |
| | kcat/KM (M-1s-1) | 5.51E+04 | 5.01E+03 | 0.18 | 1.02 |
| SARS2-G071N | kcat (s-1) | 3.19 | 0.34 | -0.22 | 1.00 |
| | KM (µM) | 69.15 | 7.88 | -0.18 | 1.02 |
| | kcat/KM (M-1s-1) | 4.65E+04 | 5.99E+03 | -0.07 | 0.60 |
| SARS2-N072G | kcat (s-1) | 2.87 | 0.59 | -0.37 | 0.94 |
| | KM (µM) | 67.57 | 11.98 | -0.21 | 0.64 |
| | kcat/KM (M-1s-1) | 4.41E+04 | 1.29E+04 | -0.14 | 0.52 |
| SARS2-Q074F | kcat (s-1) | 3.25 | 1.06 | -0.20 | 0.44 |
| | KM (µM) | 51.45 | 12.64 | -0.60 | 1.77 |
| | kcat/KM (M-1s-1) | 6.95E+04 | 3.38E+04 | 0.51 | 1.25 |
| SARS2-R076G | kcat (s-1) | 3.31 | 0.38 | -0.17 | 0.85 |
| | KM (µM) | 72.11 | 10.54 | -0.12 | 0.65 |
| | kcat/KM (M-1s-1) | 4.68E+04 | 8.41E+03 | -0.06 | 0.43 |
| SARS2-I078V | kcat (s-1) | 3.61 | 0.67 | -0.04 | 1.13 |
| | KM (µM) | 69.78 | 3.56 | -0.16 | 0.55 |
| | kcat/KM (M-1s-1) | 5.17E+04 | 8.90E+03 | 0.09 | 0.64 |
| SARS2-H080V | kcat (s-1) | 4.34 | 0.80 | 0.22 | 1.99 |
| | KM (µM) | 70.08 | 16.71 | -0.16 | 0.44 |
| | kcat/KM (M-1s-1) | 6.29E+04 | 7.58E+03 | 0.37 | 2.07 |
| SARS2-S081T | kcat (s-1) | 3.95 | 1.29 | 0.08 | 0.42 |
| | KM (µM) | 70.64 | 18.93 | -0.15 | 0.53 |
| | kcat/KM (M-1s-1) | 5.79E+04 | 1.98E+04 | 0.25 | 0.46 |
| SARS2-Q083H | kcat (s-1) | 3.48 | 0.85 | -0.10 | 0.48 |
| | KM (µM) | 72.74 | 13.29 | -0.11 | 0.52 |
| | kcat/KM (M-1s-1) | 4.78E+04 | 7.75E+03 | -0.03 | 0.36 |
| SARS2-N084G | kcat (s-1) | 3.14 | 0.46 | -0.25 | 0.63 |
| | KM (µM) | 66.07 | 17.00 | -0.24 | 0.69 |
| | kcat/KM (M-1s-1) | 4.91E+04 | 1.02E+04 | 0.01 | 0.45 |
| SARS2-C085S | kcat (s-1) | 3.19 | 1.16 | -0.22 | 0.50 |
| | KM (µM) | 81.50 | 22.13 | 0.06 | 0.53 |

| | | | | | |
|---|---|---|---|---|---|
| | kcat/KM (M-1s-1) | 3.90E+04 | 6.15E+03 | -0.32 | 1.11 |
| SARS2-K088R | kcat (s-1) | 3.64 | 1.37 | -0.03 | 0.44 |
| | KM (μM) | 74.63 | 10.90 | -0.07 | 0.32 |
| | kcat/KM (M-1s-1) | 4.88E+04 | 1.49E+04 | 0.00 | 0.38 |
| SARS2-L089I | kcat (s-1) | 2.97 | 0.89 | -0.33 | 0.66 |
| | KM (μM) | 78.02 | 11.33 | 0.00 | 0.67 |
| | kcat/KM (M-1s-1) | 3.86E+04 | 1.25E+04 | -0.33 | 0.98 |
| SARS2-D092S | kcat (s-1) | 3.57 | 0.43 | -0.06 | 0.62 |
| | KM (μM) | 69.12 | 7.05 | -0.18 | 0.69 |
| | kcat/KM (M-1s-1) | 5.19E+04 | 5.76E+03 | 0.09 | 0.91 |
| SARS2-T093Q | kcat (s-1) | 3.53 | 0.45 | -0.08 | 0.48 |
| | KM (μM) | 69.46 | 8.65 | -0.17 | 0.78 |
| | kcat/KM (M-1s-1) | 5.13E+04 | 7.02E+03 | 0.07 | 0.48 |
| SARS2-A094S | kcat (s-1) | 3.63 | 0.48 | -0.04 | 0.39 |
| | KM (μM) | 75.65 | 11.72 | -0.05 | 0.42 |
| | kcat/KM (M-1s-1) | 4.85E+04 | 6.96E+03 | -0.01 | 0.32 |
| SARS2-P096V | kcat (s-1) | 3.83 | 1.14 | 0.04 | 0.65 |
| | KM (μM) | 71.97 | 11.90 | -0.12 | 0.38 |
| | kcat/KM (M-1s-1) | 5.33E+04 | 1.33E+04 | 0.13 | 0.82 |
| SARS2-K097H | kcat (s-1) | 3.62 | 1.12 | -0.04 | 0.47 |
| | KM (μM) | 77.00 | 13.64 | -0.02 | 0.44 |
| | kcat/KM (M-1s-1) | 4.88E+04 | 2.02E+04 | 0.00 | 0.37 |
| SARS2-Y101H | kcat (s-1) | 2.35 | 0.92 | -0.67 | 1.53 |
| | KM (μM) | 64.02 | 11.96 | -0.29 | 0.94 |
| | kcat/KM (M-1s-1) | 3.80E+04 | 1.67E+04 | -0.36 | 1.12 |
| SARS2-K102V | kcat (s-1) | 2.81 | 0.54 | -0.41 | 1.66 |
| | KM (μM) | 70.66 | 12.63 | -0.15 | 0.94 |
| | kcat/KM (M-1s-1) | 4.02E+04 | 6.64E+03 | -0.28 | 1.26 |
| SARS2-V104K | kcat (s-1) | 2.52 | 0.65 | -0.57 | 1.93 |
| | KM (μM) | 72.91 | 17.42 | -0.10 | 0.58 |
| | kcat/KM (M-1s-1) | 3.65E+04 | 1.26E+04 | -0.42 | 1.14 |
| SARS2-R105T | kcat (s-1) | 2.34 | 0.54 | -0.67 | 2.17 |
| | KM (μM) | 68.66 | 8.77 | -0.19 | 0.65 |
| | kcat/KM (M-1s-1) | 3.41E+04 | 6.43E+03 | -0.52 | 1.61 |
| SARS2-I106L | kcat (s-1) | 2.80 | 0.55 | -0.41 | 2.33 |
| | KM (μM) | 60.42 | 12.38 | -0.37 | 1.62 |
| | kcat/KM (M-1s-1) | 4.79E+04 | 1.13E+04 | -0.02 | 0.36 |
| SARS2-Q107K | kcat (s-1) | 3.81 | 0.62 | 0.03 | 0.86 |
| | KM (μM) | 70.00 | 14.31 | -0.16 | 0.55 |

| | | | | | |
|---|---|---|---|---|---|
| | kcat/KM (M-1s-1) | 5.51E+04 | 6.35E+03 | 0.18 | 1.15 |
| SARS2-Q110D | kcat (s-1) | 4.45 | 1.03 | 0.26 | 1.70 |
| | KM (μM) | 84.94 | 17.02 | 0.12 | 0.67 |
| | kcat/KM (M-1s-1) | 5.27E+04 | 8.57E+03 | 0.11 | 0.77 |
| SARS2-T111S | kcat (s-1) | 3.67 | 0.80 | -0.02 | 0.61 |
| | KM (μM) | 66.13 | 12.45 | -0.24 | 0.83 |
| | kcat/KM (M-1s-1) | 5.71E+04 | 1.77E+04 | 0.23 | 1.11 |
| SARS2-S113N | kcat (s-1) | Not Active | -- | -- | -- |
| | KM (μM) | Not Active | -- | -- | -- |
| | kcat/KM (M-1s-1) | Not Active | -- | -- | -- |
| SARS2-V114I | kcat (s-1) | 2.90 | 0.34 | -0.36 | 1.28 |
| | KM (μM) | 82.49 | 22.44 | 0.08 | 0.59 |
| | kcat/KM (M-1s-1) | 3.77E+04 | 1.12E+04 | -0.37 | 0.98 |
| SARS2-N119E | kcat (s-1) | 3.92 | 1.39 | 0.07 | 0.43 |
| | KM (μM) | 63.55 | 10.12 | -0.30 | 1.28 |
| | kcat/KM (M-1s-1) | 6.18E+04 | 2.01E+04 | 0.34 | 1.46 |
| SARS2-S121I | kcat (s-1) | 4.55 | 1.13 | 0.29 | 1.38 |
| | KM (μM) | 71.40 | 15.37 | -0.13 | 0.40 |
| | kcat/KM (M-1s-1) | 6.35E+04 | 5.07E+03 | 0.38 | 3.15 |
| SARS2-P122A | kcat (s-1) | 2.77 | 0.71 | -0.43 | 1.28 |
| | KM (μM) | 62.59 | 4.29 | -0.32 | 1.37 |
| | kcat/KM (M-1s-1) | 4.46E+04 | 1.29E+04 | -0.13 | 0.45 |
| SARS2-Y126F | kcat (s-1) | 5.36 | 1.76 | 0.52 | 2.43 |
| | KM (μM) | 69.31 | 16.41 | -0.17 | 0.76 |
| | kcat/KM (M-1s-1) | 8.26E+04 | 4.17E+04 | 0.76 | 1.58 |
| SARS2-Q127G | kcat (s-1) | 0.93 | 0.45 | -2.01 | 3.79 |
| | KM (μM) | 68.99 | 19.28 | -0.18 | 0.47 |
| | kcat/KM (M-1s-1) | 1.38E+04 | 5.54E+03 | -1.81 | 3.88 |
| SARS2-C128V | kcat (s-1) | 4.40 | 1.32 | 0.24 | 1.15 |
| | KM (μM) | 66.60 | 16.20 | -0.23 | 0.57 |
| | kcat/KM (M-1s-1) | 6.68E+04 | 1.29E+04 | 0.46 | 2.05 |
| SARS2-A129N | kcat (s-1) | 0.23 | 0.12 | -4.03 | 5.10 |
| | KM (μM) | 51.72 | 25.37 | -0.60 | 1.03 |
| | kcat/KM (M-1s-1) | 4.50E+03 | 1.33E+03 | -3.44 | 4.70 |
| SARS2-M130L | kcat (s-1) | 4.39 | 1.12 | 0.24 | 1.44 |
| | KM (μM) | 78.39 | 20.89 | 0.00 | 0.31 |
| | kcat/KM (M-1s-1) | 5.70E+04 | 1.02E+04 | 0.23 | 1.08 |
| SARS2-P132T | kcat (s-1) | 3.13 | 1.23 | -0.25 | 0.58 |
| | KM (μM) | 67.16 | 12.94 | -0.22 | 0.62 |

| | | | | | |
|---|---|---|---|---|---|
| | kcat/KM (M-1s-1) | 4.66E+04 | 1.57E+04 | -0.06 | 0.36 |
| SARS2-L141I | kcat (s-1) | 0.94 | 0.12 | -1.99 | 4.91 |
| | KM (µM) | 63.47 | 13.77 | -0.30 | 0.91 |
| | kcat/KM (M-1s-1) | 1.52E+04 | 2.89E+03 | -1.68 | 3.72 |
| SARS2-S144A | kcat (s-1) | 3.86 | 1.07 | 0.05 | 0.40 |
| | KM (µM) | 243.65 | 74.99 | 1.64 | 3.40 |
| | kcat/KM (M-1s-1) | 1.60E+04 | 1.59E+03 | -1.61 | 4.38 |
| SARS2-V148P | kcat (s-1) | 3.08 | 0.90 | -0.28 | 0.80 |
| | KM (µM) | 56.33 | 15.89 | -0.47 | 1.18 |
| | kcat/KM (M-1s-1) | 5.70E+04 | 1.95E+04 | 0.23 | 0.79 |
| SARS2-F150Y | kcat (s-1) | 2.53 | 0.50 | -0.56 | 1.41 |
| | KM (µM) | 68.59 | 8.86 | -0.19 | 0.62 |
| | kcat/KM (M-1s-1) | 3.73E+04 | 7.59E+03 | -0.39 | 0.83 |
| SARS2-I152V | kcat (s-1) | 4.40 | 0.49 | 0.24 | 2.53 |
| | KM (µM) | 81.75 | 12.02 | 0.06 | 0.71 |
| | kcat/KM (M-1s-1) | 5.43E+04 | 5.93E+03 | 0.16 | 1.45 |
| SARS2-D153R | kcat (s-1) | 2.68 | 0.69 | -0.47 | 1.20 |
| | KM (µM) | 68.43 | 13.40 | -0.19 | 0.51 |
| | kcat/KM (M-1s-1) | 3.90E+04 | 4.18E+03 | -0.32 | 1.56 |
| SARS2-Y154N | kcat (s-1) | 3.99 | 0.65 | 0.10 | 2.14 |
| | KM (µM) | 87.95 | 18.45 | 0.17 | 0.79 |
| | kcat/KM (M-1s-1) | 4.72E+04 | 1.22E+04 | -0.05 | 0.32 |
| SARS2-ins154D | kcat (s-1) | 4.22 | 0.58 | 0.18 | 1.12 |
| | KM (µM) | 60.85 | 5.74 | -0.36 | 1.75 |
| | kcat/KM (M-1s-1) | 6.94E+04 | 6.74E+03 | 0.51 | 3.55 |
| SARS2-D155G | kcat (s-1) | 2.60 | 0.47 | -0.52 | 2.00 |
| | KM (µM) | 66.56 | 10.77 | -0.23 | 0.99 |
| | kcat/KM (M-1s-1) | 3.96E+04 | 6.80E+03 | -0.30 | 1.23 |
| SARS2-ins155G | kcat (s-1) | 4.81 | 0.83 | 0.37 | 2.04 |
| | KM (µM) | 70.45 | 7.41 | -0.15 | 0.46 |
| | kcat/KM (M-1s-1) | 6.90E+04 | 1.48E+04 | 0.50 | 1.84 |
| SARS2-C156T | kcat (s-1) | 3.83 | 0.82 | 0.04 | 0.96 |
| | KM (µM) | 64.13 | 9.17 | -0.29 | 0.84 |
| | kcat/KM (M-1s-1) | 6.01E+04 | 1.09E+04 | 0.30 | 1.24 |
| SARS2-S158E | kcat (s-1) | 3.69 | 0.46 | -0.01 | 0.33 |
| | KM (µM) | 70.91 | 15.32 | -0.14 | 0.97 |
| | kcat/KM (M-1s-1) | 5.36E+04 | 9.92E+03 | 0.14 | 0.77 |
| SARS2-M162L | kcat (s-1) | 0.52 | 0.29 | -2.84 | 4.71 |
| | KM (µM) | 41.86 | 10.79 | -0.90 | 2.74 |

| | | | | | |
|---|---|---|---|---|---|
| | kcat/KM (M-1s-1) | 1.22E+04 | 4.88E+03 | -2.00 | 5.11 |
| SARS2-H164Q | kcat (s-1) | 2.35 | 0.37 | -0.67 | 2.35 |
| | KM (µM) | 113.94 | 19.51 | 0.54 | 4.57 |
| | kcat/KM (M-1s-1) | 2.08E+04 | 2.83E+03 | -1.23 | 3.69 |
| SARS2-M165I | kcat (s-1) | 2.48 | 0.50 | -0.59 | 2.13 |
| | KM (µM) | 110.33 | 22.50 | 0.50 | 2.41 |
| | kcat/KM (M-1s-1) | 2.27E+04 | 2.61E+03 | -1.10 | 3.74 |
| SARS2-P168G | kcat (s-1) | 6.14 | 0.52 | 0.72 | 5.39 |
| | KM (µM) | 83.73 | 14.72 | 0.10 | 0.66 |
| | kcat/KM (M-1s-1) | 7.60E+04 | 1.91E+04 | 0.64 | 1.90 |
| SARS2-T169S | kcat (s-1) | 3.55 | 0.54 | -0.07 | 0.52 |
| | KM (µM) | 66.45 | 16.38 | -0.24 | 1.05 |
| | kcat/KM (M-1s-1) | 5.51E+04 | 1.05E+04 | 0.18 | 0.75 |
| SARS2-V171A | kcat (s-1) | 3.73 | 1.07 | 0.00 | 0.58 |
| | KM (µM) | 91.88 | 35.19 | 0.23 | 0.75 |
| | kcat/KM (M-1s-1) | 4.39E+04 | 1.45E+04 | -0.15 | 0.53 |
| SARS2-A173V | kcat (s-1) | 3.15 | 1.70 | -0.24 | 0.71 |
| | KM (µM) | 235.95 | 88.61 | 1.59 | 3.08 |
| | kcat/KM (M-1s-1) | 1.31E+04 | 3.06E+03 | -1.89 | 4.76 |
| SARS2-T175S | kcat (s-1) | 1.57 | 0.42 | -1.24 | 3.48 |
| | KM (µM) | 98.73 | 27.83 | 0.34 | 1.24 |
| | kcat/KM (M-1s-1) | 1.70E+04 | 5.59E+03 | -1.51 | 4.72 |
| SARS2-L177F | kcat (s-1) | 3.29 | 1.54 | -0.18 | 0.56 |
| | KM (µM) | 79.12 | 11.99 | 0.02 | 0.31 |
| | kcat/KM (M-1s-1) | 4.15E+04 | 1.71E+04 | -0.23 | 0.83 |
| SARS2-E178T | kcat (s-1) | 2.03 | 0.64 | -0.88 | 2.07 |
| | KM (µM) | 64.26 | 10.29 | -0.28 | 1.37 |
| | kcat/KM (M-1s-1) | 3.23E+04 | 1.07E+04 | -0.59 | 1.68 |
| SARS2-N180S | kcat (s-1) | 1.88 | 0.50 | -0.99 | 2.59 |
| | KM (µM) | 66.21 | 14.00 | -0.24 | 1.23 |
| | kcat/KM (M-1s-1) | 2.92E+04 | 8.79E+03 | -0.74 | 2.92 |
| SARS2-F181V | kcat (s-1) | Not Active | -- | -- | -- |
| | KM (µM) | Not Active | -- | -- | -- |
| | kcat/KM (M-1s-1) | Not Active | -- | -- | -- |
| SARS2-P184N | kcat (s-1) | 3.22 | 1.26 | -0.21 | 1.34 |
| | KM (µM) | 61.56 | 15.98 | -0.35 | 1.28 |
| | kcat/KM (M-1s-1) | 5.17E+04 | 7.60E+03 | 0.09 | 0.58 |
| SARS2-V186D | kcat (s-1) | 3.08 | 0.73 | -0.27 | 0.62 |
| | KM (µM) | 168.59 | 39.18 | 1.11 | 3.37 |

| | | | | | |
|---|---|---|---|---|---|
| | kcat/KM (M-1s-1) | 1.87E+04 | 4.17E+03 | -1.38 | 4.11 |
| SARS2-R188Q | kcat (s-1) | 3.69 | 0.93 | -0.02 | 0.33 |
| | KM (µM) | 91.51 | 14.47 | 0.23 | 1.15 |
| | kcat/KM (M-1s-1) | 4.08E+04 | 1.09E+04 | -0.26 | 1.02 |
| SARS2-Q189P | kcat (s-1) | 1.30 | 0.17 | -1.52 | 4.34 |
| | KM (µM) | 87.01 | 18.83 | 0.15 | 0.87 |
| | kcat/KM (M-1s-1) | 1.54E+04 | 3.33E+03 | -1.66 | 3.45 |
| SARS2-T190S | kcat (s-1) | 2.79 | 0.41 | -0.42 | 1.14 |
| | KM (µM) | 82.01 | 16.32 | 0.07 | 0.64 |
| | kcat/KM (M-1s-1) | 3.50E+04 | 7.70E+03 | -0.48 | 1.27 |
| SARS2-A191L | kcat (s-1) | 1.53 | 0.34 | -1.28 | 3.28 |
| | KM (µM) | 90.05 | 18.19 | 0.20 | 0.78 |
| | kcat/KM (M-1s-1) | 1.72E+04 | 2.49E+03 | -1.50 | 4.41 |
| SARS2-A193V | kcat (s-1) | 3.93 | 1.17 | 0.08 | 1.09 |
| | KM (µM) | 82.59 | 18.06 | 0.08 | 0.71 |
| | kcat/KM (M-1s-1) | 4.79E+04 | 9.67E+03 | -0.03 | 0.34 |
| SARS2-A194E | kcat (s-1) | 2.88 | 0.87 | -0.37 | 2.13 |
| | KM (µM) | 80.57 | 19.92 | 0.04 | 0.42 |
| | kcat/KM (M-1s-1) | 3.56E+04 | 4.61E+03 | -0.45 | 1.95 |
| SARS2-G195S | kcat (s-1) | 3.74 | 1.94 | 0.01 | 0.31 |
| | KM (µM) | 65.21 | 14.75 | -0.26 | 1.36 |
| | kcat/KM (M-1s-1) | 5.66E+04 | 2.03E+04 | 0.22 | 0.68 |
| SARS2-T196A | kcat (s-1) | 3.71 | 0.69 | -0.01 | 0.31 |
| | KM (µM) | 76.83 | 15.02 | -0.03 | 0.39 |
| | kcat/KM (M-1s-1) | 4.94E+04 | 1.05E+04 | 0.02 | 0.36 |
| SARS2-D197N | kcat (s-1) | 1.51 | 0.72 | -1.30 | 3.51 |
| | KM (µM) | 77.87 | 24.25 | -0.01 | 0.50 |
| | kcat/KM (M-1s-1) | 2.00E+04 | 7.97E+03 | -1.28 | 4.30 |
| SARS2-T198L | kcat (s-1) | 4.68 | 1.70 | 0.33 | 1.25 |
| | KM (µM) | 77.63 | 11.72 | -0.01 | 0.45 |
| | kcat/KM (M-1s-1) | 5.95E+04 | 1.57E+04 | 0.29 | 1.37 |
| SARS2-T199M | kcat (s-1) | 1.55 | 0.50 | -1.27 | 3.12 |
| | KM (µM) | 77.71 | 13.78 | -0.01 | 0.41 |
| | kcat/KM (M-1s-1) | 1.95E+04 | 3.36E+03 | -1.32 | 3.72 |
| SARS2-I200L | kcat (s-1) | 2.18 | 0.54 | -0.77 | 2.15 |
| | KM (µM) | 63.82 | 4.51 | -0.29 | 1.24 |
| | kcat/KM (M-1s-1) | 3.46E+04 | 9.71E+03 | -0.49 | 1.51 |
| SARS2-T201S | kcat (s-1) | 2.45 | 0.74 | -0.61 | 1.37 |
| | KM (µM) | 65.68 | 8.38 | -0.25 | 0.70 |

| | | | | | |
|---|---|---|---|---|---|
| | kcat/KM (M-1s-1) | 3.71E+04 | 8.94E+03 | -0.39 | 1.66 |
| SARS2-V202D | kcat (s-1) | Not Active | -- | -- | -- |
| | KM (μM) | Not Active | -- | -- | -- |
| | kcat/KM (M-1s-1) | Not Active | -- | -- | -- |
| SARS2-L205V | kcat (s-1) | 2.94 | 0.80 | -0.34 | 0.95 |
| | KM (μM) | 68.01 | 11.67 | -0.20 | 0.60 |
| | kcat/KM (M-1s-1) | 4.33E+04 | 9.98E+03 | -0.17 | 0.69 |
| SARS2-W207F | kcat (s-1) | 2.91 | 0.23 | -0.36 | 1.03 |
| | KM (μM) | 77.62 | 17.96 | -0.01 | 0.40 |
| | kcat/KM (M-1s-1) | 3.90E+04 | 8.19E+03 | -0.32 | 0.87 |
| SARS2-V212L | kcat (s-1) | 1.52 | 0.59 | -1.29 | 2.69 |
| | KM (μM) | 58.30 | 11.86 | -0.42 | 1.07 |
| | kcat/KM (M-1s-1) | 2.63E+04 | 9.42E+03 | -0.89 | 2.24 |
| SARS2-I213L | kcat (s-1) | 3.40 | 1.17 | -0.13 | 1.42 |
| | KM (μM) | 81.03 | 15.66 | 0.05 | 0.53 |
| | kcat/KM (M-1s-1) | 4.24E+04 | 1.16E+04 | -0.20 | 1.32 |
| SARS2-D216C | kcat (s-1) | 3.22 | 0.39 | -0.21 | 0.90 |
| | KM (μM) | 52.49 | 12.45 | -0.58 | 1.88 |
| | kcat/KM (M-1s-1) | 6.41E+04 | 1.60E+04 | 0.40 | 1.23 |
| SARS2-F219W | kcat (s-1) | 3.14 | 0.91 | -0.25 | 0.71 |
| | KM (μM) | 73.49 | 15.65 | -0.09 | 0.53 |
| | kcat/KM (M-1s-1) | 4.28E+04 | 8.56E+03 | -0.19 | 0.87 |
| SARS2-N221C | kcat (s-1) | 3.52 | 1.14 | -0.08 | 0.43 |
| | KM (μM) | 66.75 | 21.88 | -0.23 | 1.03 |
| | kcat/KM (M-1s-1) | 5.80E+04 | 2.54E+04 | 0.25 | 0.72 |
| SARS2-R222S | kcat (s-1) | 3.89 | 1.07 | 0.06 | 0.68 |
| | KM (μM) | 79.01 | 18.40 | 0.01 | 0.49 |
| | kcat/KM (M-1s-1) | 5.12E+04 | 1.48E+04 | 0.07 | 0.52 |
| SARS2-F223T | kcat (s-1) | 4.21 | 0.70 | 0.18 | 0.68 |
| | KM (μM) | 79.00 | 15.62 | 0.01 | 0.34 |
| | kcat/KM (M-1s-1) | 5.42E+04 | 8.73E+03 | 0.15 | 0.81 |
| SARS2-T224R | kcat (s-1) | 3.56 | 1.70 | -0.06 | 1.10 |
| | KM (μM) | 70.28 | 14.59 | -0.15 | 0.60 |
| | kcat/KM (M-1s-1) | 5.15E+04 | 2.50E+04 | 0.08 | 0.89 |
| SARS2-T225V | kcat (s-1) | Not Active | -- | -- | -- |
| | KM (μM) | Not Active | -- | -- | -- |
| | kcat/KM (M-1s-1) | Not Active | -- | -- | -- |
| SARS2-T226N | kcat (s-1) | 2.90 | 0.52 | -0.36 | 0.98 |
| | KM (μM) | 58.81 | 13.01 | -0.41 | 1.35 |

| | | | | | |
|---|---|---|---|---|---|
| | kcat/KM (M-1s-1) | 5.01E+04 | 5.36E+03 | 0.04 | 0.55 |
| SARS2-L227V | kcat (s-1) | 2.93 | 1.02 | -0.35 | 1.01 |
| | KM (µM) | 69.06 | 17.95 | -0.18 | 0.89 |
| | kcat/KM (M-1s-1) | 4.29E+04 | 1.21E+04 | -0.18 | 0.65 |
| SARS2-N228D | kcat (s-1) | Not Active | -- | -- | -- |
| | KM (µM) | Not Active | -- | -- | -- |
| | kcat/KM (M-1s-1) | Not Active | -- | -- | -- |
| SARS2-D229G | kcat (s-1) | 0.89 | 0.31 | -2.07 | 4.04 |
| | KM (µM) | 61.45 | 12.57 | -0.35 | 1.72 |
| | kcat/KM (M-1s-1) | 1.44E+04 | 3.90E+03 | -1.76 | 4.01 |
| SARS2-L232E | kcat (s-1) | 4.34 | 1.39 | 0.22 | 0.70 |
| | KM (µM) | 77.76 | 22.59 | -0.01 | 0.31 |
| | kcat/KM (M-1s-1) | 5.60E+04 | 5.97E+03 | 0.20 | 1.21 |
| SARS2-V233W | kcat (s-1) | 2.84 | 0.62 | -0.39 | 1.37 |
| | KM (µM) | 67.30 | 9.63 | -0.22 | 0.59 |
| | kcat/KM (M-1s-1) | 4.24E+04 | 7.77E+03 | -0.20 | 0.71 |
| SARS2-K236A | kcat (s-1) | 3.69 | 1.50 | -0.01 | 0.32 |
| | KM (µM) | 79.92 | 17.25 | 0.03 | 0.42 |
| | kcat/KM (M-1s-1) | 4.67E+04 | 1.71E+04 | -0.06 | 0.41 |
| SARS2-Y237N | kcat (s-1) | 2.93 | 0.90 | -0.34 | 0.74 |
| | KM (µM) | 56.14 | 13.88 | -0.48 | 1.39 |
| | kcat/KM (M-1s-1) | 5.76E+04 | 3.26E+04 | 0.24 | 0.57 |
| SARS2-N238G | kcat (s-1) | 4.05 | 0.79 | 0.12 | 2.15 |
| | KM (µM) | 81.76 | 14.54 | 0.06 | 0.56 |
| | kcat/KM (M-1s-1) | 5.13E+04 | 1.39E+04 | 0.07 | 0.56 |
| SARS2-E240T | kcat (s-1) | Not Active | -- | -- | -- |
| | KM (µM) | Not Active | -- | -- | -- |
| | kcat/KM (M-1s-1) | Not Active | -- | -- | -- |
| SARS2-P241S | kcat (s-1) | 2.85 | 0.64 | -0.39 | 1.25 |
| | KM (µM) | 70.06 | 12.02 | -0.16 | 0.73 |
| | kcat/KM (M-1s-1) | 4.05E+04 | 4.48E+03 | -0.27 | 1.18 |
| SARS2-L242V | kcat (s-1) | 3.12 | 0.58 | -0.26 | 0.89 |
| | KM (µM) | 79.43 | 13.48 | 0.02 | 0.34 |
| | kcat/KM (M-1s-1) | 3.98E+04 | 7.78E+03 | -0.29 | 1.11 |
| SARS2-T243S | kcat (s-1) | 2.85 | 0.76 | -0.39 | 1.60 |
| | KM (µM) | 70.75 | 18.43 | -0.14 | 0.44 |
| | kcat/KM (M-1s-1) | 4.19E+04 | 1.04E+04 | -0.22 | 1.11 |
| SARS2-Q244S | kcat (s-1) | 3.99 | 1.52 | 0.10 | 0.71 |
| | KM (µM) | 81.72 | 17.96 | 0.06 | 0.63 |

| | | | | | |
|---|---|---|---|---|---|
| | kcat/KM (M-1s-1) | 5.03E+04 | 1.99E+04 | 0.05 | 0.46 |
| SARS2-D245V | kcat (s-1) | 2.81 | 0.43 | -0.41 | 1.55 |
| | KM (µM) | 58.79 | 13.55 | -0.41 | 1.21 |
| | kcat/KM (M-1s-1) | 4.93E+04 | 9.04E+03 | 0.02 | 0.46 |
| SARS2-D245Δ | kcat (s-1) | Not Active | -- | -- | -- |
| | KM (µM) | Not Active | -- | -- | -- |
| | kcat/KM (M-1s-1) | Not Active | -- | -- | -- |
| SARS2-H246E | kcat (s-1) | 3.03 | 0.66 | -0.30 | 1.07 |
| | KM (µM) | 68.64 | 13.69 | -0.19 | 0.72 |
| | kcat/KM (M-1s-1) | 4.57E+04 | 1.30E+04 | -0.09 | 0.50 |
| SARS2-H246Δ | kcat (s-1) | 1.71 | 0.24 | -1.13 | 2.35 |
| | KM (µM) | 76.44 | 14.25 | -0.03 | 0.41 |
| | kcat/KM (M-1s-1) | 2.31E+04 | 5.27E+03 | -1.08 | 2.87 |
| SARS2-V247Δ | kcat (s-1) | 4.28 | #DIV/0! | 0.20 | #DIV/0! |
| | KM (µM) | 76.85 | #DIV/0! | -0.03 | #DIV/0! |
| | kcat/KM (M-1s-1) | #VALUE! | #VALUE! | #VALUE! | #VALUE! |
| SARS2-D248Δ | kcat (s-1) | Not Active | -- | -- | -- |
| | KM (µM) | Not Active | -- | -- | -- |
| | kcat/KM (M-1s-1) | Not Active | -- | -- | -- |
| SARS2-D248E | kcat (s-1) | 3.73 | 1.50 | 0.00 | 0.65 |
| | KM (µM) | 82.60 | 9.63 | 0.08 | 0.71 |
| | kcat/KM (M-1s-1) | 4.60E+04 | 1.85E+04 | -0.08 | 0.35 |
| SARS2-I249C | kcat (s-1) | 3.73 | 0.85 | 0.00 | 0.60 |
| | KM (µM) | 69.75 | 7.84 | -0.17 | 0.63 |
| | kcat/KM (M-1s-1) | 5.46E+04 | 1.62E+04 | 0.17 | 0.88 |
| SARS2-L250Y | kcat (s-1) | Not Active | -- | -- | -- |
| | KM (µM) | Not Active | -- | -- | -- |
| | kcat/KM (M-1s-1) | Not Active | -- | -- | -- |
| SARS2-G251S | kcat (s-1) | 3.32 | 0.96 | -0.17 | 1.48 |
| | KM (µM) | 69.16 | 17.88 | -0.18 | 0.74 |
| | kcat/KM (M-1s-1) | 4.94E+04 | 1.18E+04 | 0.02 | 0.36 |
| SARS2-P252I | kcat (s-1) | 3.05 | 0.80 | -0.29 | 0.43 |
| | KM (µM) | 66.01 | 10.96 | -0.24 | 0.91 |
| | kcat/KM (M-1s-1) | 4.58E+04 | 4.54E+03 | -0.09 | 0.35 |
| SARS2-S254A | kcat (s-1) | 4.16 | 1.23 | 0.16 | 1.68 |
| | KM (µM) | 68.39 | 16.54 | -0.19 | 1.17 |
| | kcat/KM (M-1s-1) | 6.04E+04 | 4.98E+03 | 0.31 | 2.02 |
| SARS2-Q256K | kcat (s-1) | 3.64 | 1.06 | -0.03 | 0.61 |

| | | | | | |
|---|---|---|---|---|---|
| | KM (µM) | 74.76 | 9.62 | -0.07 | 0.32 |
| | kcat/KM (M-1s-1) | 4.85E+04 | 1.12E+04 | 0.00 | 0.37 |
| SARS2-I259V | kcat (s-1) | 3.61 | 0.92 | -0.05 | 0.35 |
| | KM (µM) | 72.41 | 15.20 | -0.11 | 0.49 |
| | kcat/KM (M-1s-1) | 5.33E+04 | 2.55E+04 | 0.13 | 0.52 |
| SARS2-A260S | kcat (s-1) | 4.44 | 1.55 | 0.25 | 1.16 |
| | KM (µM) | 75.46 | 27.24 | -0.05 | 0.33 |
| | kcat/KM (M-1s-1) | 6.02E+04 | 1.27E+04 | 0.31 | 1.51 |
| SARS2-L262E | kcat (s-1) | 2.85 | 0.73 | -0.39 | 1.62 |
| | KM (µM) | 78.49 | 16.67 | 0.00 | 0.55 |
| | kcat/KM (M-1s-1) | 3.77E+04 | 1.10E+04 | -0.37 | 1.05 |
| SARS2-D263Q | kcat (s-1) | Not Active | -- | -- | -- |
| | KM (µM) | Not Active | -- | -- | -- |
| | kcat/KM (M-1s-1) | Not Active | -- | -- | -- |
| SARS2-M264L | kcat (s-1) | 2.86 | 0.85 | -0.38 | 0.85 |
| | KM (µM) | 79.68 | 17.99 | 0.03 | 0.82 |
| | kcat/KM (M-1s-1) | 3.71E+04 | 1.33E+04 | -0.39 | 1.49 |
| SARS2-C265L | kcat (s-1) | 4.32 | 1.86 | 0.21 | 0.82 |
| | KM (µM) | 91.18 | 12.55 | 0.22 | 1.48 |
| | kcat/KM (M-1s-1) | 4.66E+04 | 1.57E+04 | -0.06 | 0.41 |
| SARS2-L268I | kcat (s-1) | 0.09 | 0.08 | -5.39 | 1.41 |
| | KM (µM) | 64.86 | 67.40 | -0.27 | 0.49 |
| | kcat/KM (M-1s-1) | 1.66E+03 | 5.45E+02 | -4.88 | 1.02 |
| SARS2-K269Q | kcat (s-1) | 4.09 | 0.94 | 0.13 | 0.63 |
| | KM (µM) | 76.38 | 19.31 | -0.03 | 0.37 |
| | kcat/KM (M-1s-1) | 5.48E+04 | 1.01E+04 | 0.17 | 0.75 |
| SARS2-E270Δ | kcat (s-1) | Not Active | -- | -- | -- |
| | KM (µM) | Not Active | -- | -- | -- |
| | kcat/KM (M-1s-1) | Not Active | -- | -- | -- |
| SARS2-E270H | kcat (s-1) | 3.62 | 1.21 | -0.04 | 0.50 |
| | KM (µM) | 60.11 | 11.05 | -0.38 | 1.36 |
| | kcat/KM (M-1s-1) | 6.43E+04 | 3.62E+04 | 0.40 | 0.89 |
| SARS2-L271H | kcat (s-1) | 0.39 | 0.08 | -3.25 | 5.06 |
| | KM (µM) | 54.92 | 24.64 | -0.51 | 1.54 |
| | kcat/KM (M-1s-1) | 8.25E+03 | 3.45E+03 | -2.56 | 4.43 |
| SARS2-L272H | kcat (s-1) | Not Active | -- | -- | -- |
| | KM (µM) | Not Active | -- | -- | -- |
| | kcat/KM (M-1s-1) | Not Active | -- | -- | -- |
| SARS2-Q273H | kcat (s-1) | Not Active | -- | -- | -- |

| | | | | | |
|---|---|---|---|---|---|
| | KM (μM) | Not Active | -- | -- | -- |
| | kcat/KM (M-1s-1) | Not Active | -- | -- | -- |
| SARS2-Q273E | kcat (s-1) | 4.48 | 1.79 | 0.27 | 1.12 |
| | KM (μM) | 70.56 | 5.60 | -0.15 | 0.77 |
| | kcat/KM (M-1s-1) | 6.40E+04 | 2.77E+04 | 0.39 | 1.14 |
| SARS2-N274E | kcat (s-1) | 3.25 | 0.89 | -0.20 | 1.22 |
| | KM (μM) | 69.86 | 16.21 | -0.16 | 0.64 |
| | kcat/KM (M-1s-1) | 4.84E+04 | 1.51E+04 | -0.01 | 0.31 |
| SARS2-N274Δ | kcat (s-1) | 0.30 | 0.11 | -3.65 | 4.74 |
| | KM (μM) | 47.30 | 16.82 | -0.73 | 1.95 |
| | kcat/KM (M-1s-1) | 6.48E+03 | 1.71E+03 | -2.91 | 4.92 |
| SARS2-M276F | kcat (s-1) | 3.61 | 1.00 | -0.05 | 0.46 |
| | KM (μM) | 68.03 | 10.02 | -0.20 | 1.03 |
| | kcat/KM (M-1s-1) | 5.48E+04 | 2.01E+04 | 0.17 | 0.83 |
| SARS2-N277G | kcat (s-1) | 3.39 | 0.84 | -0.14 | 1.13 |
| | KM (μM) | 68.09 | 10.56 | -0.20 | 0.93 |
| | kcat/KM (M-1s-1) | 5.07E+04 | 1.25E+04 | 0.06 | 0.38 |
| SARS2-R279K | kcat (s-1) | 3.46 | 0.80 | -0.11 | 0.56 |
| | KM (μM) | 68.30 | 10.61 | -0.20 | 0.88 |
| | kcat/KM (M-1s-1) | 5.13E+04 | 1.13E+04 | 0.07 | 0.46 |
| SARS2-T280N | kcat (s-1) | 2.69 | 0.59 | -0.47 | 1.29 |
| | KM (μM) | 74.09 | 14.48 | -0.08 | 0.39 |
| | kcat/KM (M-1s-1) | 3.68E+04 | 8.44E+03 | -0.40 | 1.74 |
| SARS2-S284Y | kcat (s-1) | 2.24 | 0.99 | -0.73 | 2.16 |
| | KM (μM) | 76.01 | 15.83 | -0.04 | 0.46 |
| | kcat/KM (M-1s-1) | 2.91E+04 | 1.09E+04 | -0.74 | 3.15 |
| SARS2-A285S | kcat (s-1) | 2.84 | 0.91 | -0.39 | 1.14 |
| | KM (μM) | 81.36 | 13.08 | 0.06 | 0.70 |
| | kcat/KM (M-1s-1) | 3.50E+04 | 1.04E+04 | -0.48 | 1.37 |
| SARS2-L286S | kcat (s-1) | 4.35 | 0.65 | 0.22 | 0.84 |
| | KM (μM) | 79.93 | 16.32 | 0.03 | 0.37 |
| | kcat/KM (M-1s-1) | 5.62E+04 | 1.42E+04 | 0.21 | 0.77 |
| SARS2-E288C | kcat (s-1) | 0.75 | 0.25 | -2.30 | 4.81 |
| | KM (μM) | 76.44 | 34.98 | -0.03 | 0.36 |
| | kcat/KM (M-1s-1) | 1.10E+04 | 4.58E+03 | -2.14 | 4.23 |
| SARS2-P293L | kcat (s-1) | 1.33 | 0.36 | -1.49 | 3.78 |
| | KM (μM) | 71.77 | 13.09 | -0.12 | 0.44 |
| | kcat/KM (M-1s-1) | 1.94E+04 | 6.68E+03 | -1.33 | 3.12 |
| SARS2-F294A | kcat (s-1) | 4.49 | 1.03 | 0.27 | 1.53 |

| | | | | | |
|---|---|---|---|---|---|
| | KM (µM) | 80.65 | 12.44 | 0.04 | 0.39 |
| | kcat/KM (M-1s-1) | 5.53E+04 | 6.90E+03 | 0.18 | 1.55 |
| SARS2-D295E | kcat (s-1) | 1.96 | 0.38 | -0.92 | 2.32 |
| | KM (µM) | 93.44 | 11.87 | 0.26 | 1.57 |
| | kcat/KM (M-1s-1) | 2.11E+04 | 3.87E+03 | -1.20 | 4.23 |
| SARS2-R298K | kcat (s-1) | 2.58 | 0.95 | -0.53 | 1.92 |
| | KM (µM) | 82.14 | 20.11 | 0.07 | 0.44 |
| | kcat/KM (M-1s-1) | 3.21E+04 | 9.72E+03 | -0.60 | 3.54 |
| SARS2-C300M | kcat (s-1) | 3.22 | 0.64 | -0.21 | 1.00 |
| | KM (µM) | 68.38 | 11.17 | -0.19 | 0.88 |
| | kcat/KM (M-1s-1) | 4.74E+04 | 6.84E+03 | -0.04 | 0.53 |
| SARS2-S301Y | kcat (s-1) | 4.28 | 1.17 | 0.20 | 1.10 |
| | KM (µM) | 41.27 | 8.22 | -0.92 | 2.44 |
| | kcat/KM (M-1s-1) | 1.09E+05 | 4.60E+04 | 1.16 | 2.00 |
| SARS2-T304N | kcat (s-1) | 2.98 | 1.28 | -0.32 | 0.92 |
| | KM (µM) | 93.60 | 17.78 | 0.26 | 1.15 |
| | kcat/KM (M-1s-1) | 3.19E+04 | 1.04E+04 | -0.61 | 1.77 |
| SARS2-F305L | kcat (s-1) | Not Active | -- | -- | -- |
| | KM (µM) | Not Active | -- | -- | -- |
| | kcat/KM (M-1s-1) | Not Active | -- | -- | -- |
| SARS2-DH245-246ΔΔ | kcat (s-1) | Not Active | -- | -- | -- |
| | KM (µM) | Not Active | -- | -- | -- |
| | kcat/KM (M-1s-1) | Not Active | -- | -- | -- |
| SARS2-VD247-248ΔΔ | kcat (s-1) | Not Active | -- | -- | -- |
| | KM (µM) | Not Active | -- | -- | -- |
| | kcat/KM (M-1s-1) | Not Active | -- | -- | -- |
| SARS2-NL63-Loop1 | kcat (s-1) | 1.75 | #DIV/0! | -1.09 | #DIV/0! |
| | KM (µM) | 53.39 | #DIV/0! | -0.55 | #DIV/0! |
| | kcat/KM (M-1s-1) | #VALUE! | #VALUE! | #VALUE! | #VALUE! |
| SARS2-NL63-Loop22 | kcat (s-1) | 1.11 | 0.53 | -1.75 | 3.00 |
| | KM (µM) | 84.42 | 29.79 | 0.11 | 0.46 |
| | kcat/KM (M-1s-1) | 1.23E+04 | 3.53E+03 | -1.99 | 4.55 |
| SARS2-NL63-Loop39 | kcat (s-1) | 0.05 | 0.02 | -6.19 | 3.33 |
| | KM (µM) | 52.91 | 25.13 | -0.56 | 1.57 |
| | kcat/KM (M-1s-1) | 1.03E+03 | 3.40E+02 | -5.56 | 3.99 |
| SARS2-NL63-Loop70 | kcat (s-1) | 3.71 | 0.87 | -0.01 | 0.37 |
| | KM (µM) | 83.10 | 17.00 | 0.09 | 0.60 |
| | kcat/KM (M-1s-1) | 4.48E+04 | 6.78E+03 | -0.12 | 0.95 |

| | | | | | |
|---|---|---|---|---|---|
| SARS2-NL63-Loop83 | kcat (s-1) | 2.34 | 0.50 | -0.67 | 2.05 |
| | KM (µM) | 79.31 | 18.03 | 0.02 | 0.46 |
| | kcat/KM (M-1s-1) | 3.11E+04 | 9.43E+03 | -0.65 | 1.72 |
| SARS2-NL63-Loop91 | kcat (s-1) | 1.37 | 0.28 | -1.44 | 2.90 |
| | KM (µM) | 64.89 | 19.13 | -0.27 | 1.08 |
| | kcat/KM (M-1s-1) | 2.26E+04 | 6.77E+03 | -1.11 | 2.68 |
| SARS2-NL63-Loop129 | kcat (s-1) | Not Active | -- | -- | -- |
| | KM (µM) | Not Active | -- | -- | -- |
| | kcat/KM (M-1s-1) | Not Active | -- | -- | -- |
| SARS2-NL63-Loop166 | kcat (s-1) | 4.39 | 0.76 | 0.24 | 1.11 |
| | KM (µM) | 86.63 | 10.80 | 0.15 | 1.41 |
| | kcat/KM (M-1s-1) | 5.08E+04 | 7.53E+03 | 0.06 | 0.32 |
| SARS2-NL63-Loop175 | kcat (s-1) | Not Active | -- | -- | -- |
| | KM (µM) | Not Active | -- | -- | -- |
| | kcat/KM (M-1s-1) | Not Active | -- | -- | -- |
| SARS2-NL63-Loop275 | kcat (s-1) | 3.82 | 0.62 | 0.03 | 0.80 |
| | KM (µM) | 42.52 | 7.12 | -0.88 | 3.16 |
| | kcat/KM (M-1s-1) | 9.00E+04 | 8.27E+03 | 0.89 | 4.48 |
| SARS2-NL63-Loop300 | kcat (s-1) | 0.52 | 0.17 | -2.85 | 4.12 |
| | KM (µM) | 65.99 | 22.56 | -0.25 | 0.66 |
| | kcat/KM (M-1s-1) | 8.24E+03 | 2.39E+03 | -2.56 | 3.26 |
| SARS2-NL63-Loop39/129 | kcat (s-1) | Not Active | -- | -- | -- |
| | KM (µM) | Not Active | -- | -- | -- |
| | kcat/KM (M-1s-1) | Not Active | -- | -- | -- |
| SARS2-NL63-Loop39/166 | kcat (s-1) | 0.15 | 0.07 | -4.64 | 4.61 |
| | KM (µM) | 122.24 | 57.22 | 0.64 | 1.12 |
| | kcat/KM (M-1s-1) | 1.27E+03 | 2.52E+02 | -5.26 | 5.49 |
| SARS2-NL63-Loop39/175 | kcat (s-1) | Not Active | -- | -- | -- |
| | KM (µM) | Not Active | -- | -- | -- |
| | kcat/KM (M-1s-1) | Not Active | -- | -- | -- |
| SARS2-NL63-Loop39/129/166 | kcat (s-1) | Not Active | -- | -- | -- |
| | KM (µM) | Not Active | -- | -- | -- |
| | kcat/KM (M-1s-1) | Not Active | -- | -- | -- |
| SARS2-NL63-Loop39/129/175 | kcat (s-1) | Not Active | -- | -- | -- |
| | KM (µM) | Not Active | -- | -- | -- |
| | kcat/KM (M-1s-1) | Not Active | -- | -- | -- |
| SARS2-NL63-Loop39/166/175 | kcat (s-1) | Not Active | -- | -- | -- |
| | KM (µM) | Not Active | -- | -- | -- |
| | kcat/KM (M-1s-1) | Not Active | -- | -- | -- |

| | | | | | |
|---|---|---|---|---|---|
| SARS2-NL63-Loop129/166 | kcat (s-1) | Not Active | -- | -- | -- |
| | KM (μM) | Not Active | -- | -- | -- |
| | kcat/KM (M-1s-1) | Not Active | -- | -- | -- |
| SARS2-NL63-Loop129/175 | kcat (s-1) | Not Active | -- | -- | -- |
| | KM (μM) | Not Active | -- | -- | -- |
| | kcat/KM (M-1s-1) | Not Active | -- | -- | -- |
| SARS2-NL63-Loop129/166/175 | kcat (s-1) | Not Active | -- | -- | -- |
| | KM (μM) | Not Active | -- | -- | -- |
| | kcat/KM (M-1s-1) | Not Active | -- | -- | -- |
| SARS2-NL63-Loop166/175 | kcat (s-1) | Not Active | -- | -- | -- |
| | KM (μM) | Not Active | -- | -- | -- |
| | kcat/KM (M-1s-1) | Not Active | -- | -- | -- |
| SARS2-NL63-Loop39/129/166/175 | kcat (s-1) | Not Active | -- | -- | -- |
| | KM (μM) | Not Active | -- | -- | -- |
| | kcat/KM (M-1s-1) | Not Active | -- | -- | -- |

**Table S6. Ensitrelvir Inhibition Values for functional and clinically focused sublibrary**

| Mutant ID | IC50 (M) | St. Dev | Log2FC | log10(q-value) |
|---|---|---|---|---|
| SARS2-WT | 3.21E-09 | 9.02E-10 | 0.00 | N/A |
| SARS2-C145A | Not Active | -- | -- | -- |
| SARS2-TQA-WT | Not Active | -- | -- | -- |
| SARS2-TQA-C145A | Not Active | -- | -- | -- |
| SARS2-R004S | Not Active | -- | -- | -- |
| SARS2-K005F | 2.47E-09 | 8.63E-10 | -0.38 | 1.45 |
| SARS2-K005W | 2.41E-09 | 5.50E-10 | -0.41 | 1.95 |
| SARS2-A007T | 2.77E-09 | 5.78E-10 | -0.21 | 0.59 |
| SARS2-G015S | 3.04E-09 | 5.63E-10 | -0.07 | 0.38 |
| SARS2-G015V | 2.66E-09 | 7.94E-10 | -0.27 | 0.74 |
| SARS2-T021I | 2.30E-09 | 3.77E-10 | -0.48 | 1.89 |
| SARS2-C022I | 3.25E-09 | 8.72E-10 | 0.02 | 0.36 |
| SARS2-T025I | 6.43E-09 | 9.93E-10 | 1.01 | 3.66 |
| SARS2-L030I | 3.02E-09 | 9.21E-10 | -0.08 | 0.82 |
| SARS2-D033F | 2.58E-09 | 5.79E-10 | -0.31 | 1.63 |
| SARS2-D033V | 3.74E-09 | 1.59E-09 | 0.22 | 0.61 |
| SARS2-D033Y | Not Active | -- | -- | -- |
| SARS2-D034M | 4.67E-09 | 1.40E-09 | 0.54 | 1.37 |
| SARS2-H041L | Not Active | -- | -- | -- |
| SARS2-H041Y | Not Active | -- | -- | -- |
| SARS2-T045I | 4.05E-09 | 1.03E-09 | 0.34 | 1.47 |
| SARS2-S046F | 3.09E-09 | 8.00E-10 | -0.05 | 0.37 |
| SARS2-S046P | 3.44E-09 | 1.03E-09 | 0.10 | 0.54 |
| SARS2-E047K | 2.49E-09 | 7.85E-10 | -0.37 | 0.91 |
| SARS2-E047N | 2.89E-09 | 7.16E-10 | -0.15 | 0.53 |
| SARS2-D048N | 2.74E-09 | 4.54E-10 | -0.23 | 0.83 |
| SARS2-M049I | 8.06E-09 | 2.73E-09 | 1.33 | 3.07 |
| SARS2-M049K | 5.33E-09 | 9.45E-10 | 0.73 | 3.39 |
| SARS2-M049L | 1.48E-08 | 2.14E-09 | 2.21 | 4.96 |
| SARS2-M049T | 5.20E-09 | 1.81E-09 | 0.70 | 1.91 |
| SARS2-M049V | 5.65E-09 | 1.52E-09 | 0.82 | 2.60 |
| SARS2-L050F | 3.04E-09 | 2.48E-09 | -0.08 | 0.37 |
| SARS2-V073I | Not Active | -- | -- | -- |
| SARS2-Q083L | 3.15E-09 | 3.63E-10 | -0.03 | 0.30 |
| SARS2-K088R | 2.58E-09 | 5.19E-10 | -0.32 | 1.12 |
| SARS2-L089F | 4.13E-09 | 1.46E-09 | 0.37 | 1.11 |

| | | | | |
|---|---|---|---|---|
| SARS2-L089H | 2.64E-09 | 3.95E-10 | -0.28 | 0.99 |
| SARS2-L089I | 3.44E-09 | 1.73E-09 | 0.10 | 0.48 |
| SARS2-L089P | 2.43E-09 | 4.35E-10 | -0.40 | 1.32 |
| SARS2-L089R | 3.74E-09 | 3.73E-09 | 0.22 | 0.48 |
| SARS2-L089V | 3.44E-09 | 1.06E-09 | 0.10 | 0.54 |
| SARS2-K090E | 3.40E-09 | 7.57E-10 | 0.08 | 0.83 |
| SARS2-K090M | 4.62E-09 | 2.38E-09 | 0.53 | 0.88 |
| SARS2-K090N | 3.71E-09 | 1.16E-09 | 0.21 | 0.46 |
| SARS2-K090Q | 3.41E-09 | 1.36E-09 | 0.09 | 0.42 |
| SARS2-K090T | 4.25E-09 | 1.68E-09 | 0.41 | 0.90 |
| SARS2-K090R | 3.40E-09 | 1.98E-09 | 0.09 | 0.42 |
| SARS2-D092G | 2.93E-09 | 1.19E-09 | -0.13 | 0.49 |
| SARS2-T093I | 3.21E-09 | 8.23E-10 | 0.00 | 0.30 |
| SARS2-P096L | 2.90E-09 | 5.16E-10 | -0.15 | 0.42 |
| SARS2-P096S | Not Active | -- | -- | -- |
| SARS2-P108S | 3.30E-09 | 1.40E-09 | 0.04 | 0.50 |
| SARS2-S113T | 2.58E-09 | 4.73E-10 | -0.31 | 1.25 |
| SARS2-A116E | 7.88E-09 | 1.69E-09 | 1.30 | 4.43 |
| SARS2-A116L | 7.85E-09 | 2.09E-09 | 1.29 | 4.14 |
| SARS2-A116V | 4.36E-09 | 1.06E-09 | 0.44 | 1.56 |
| SARS2-S123F | 3.42E-09 | 5.14E-10 | 0.10 | 0.49 |
| SARS2-Y126F | 3.63E-09 | 1.22E-09 | 0.18 | 0.55 |
| SARS2-Y126M | 2.41E-09 | 4.32E-10 | -0.41 | 1.47 |
| SARS2-Q127H | 2.76E-09 | 5.16E-10 | -0.21 | 1.15 |
| SARS2-A129V | 3.66E-09 | 1.05E-09 | 0.19 | 0.65 |
| SARS2-P132A | 2.85E-09 | 6.32E-10 | -0.17 | 0.86 |
| SARS2-P132H | 3.41E-09 | 1.65E-09 | 0.09 | 0.57 |
| SARS2-P132L | 5.89E-09 | 7.41E-09 | 0.88 | 0.77 |
| SARS2-P132R | 3.09E-09 | 6.31E-10 | -0.05 | 0.40 |
| SARS2-P132S | 3.36E-09 | 1.22E-09 | 0.07 | 0.48 |
| SARS2-P132T | 3.02E-09 | 7.64E-10 | -0.08 | 0.46 |
| SARS2-F140M | 3.97E-09 | 1.23E-09 | 0.31 | 1.22 |
| SARS2-L141A | 4.25E-09 | 5.19E-10 | 0.41 | 1.82 |
| SARS2-L141E | 2.88E-09 | 4.60E-10 | -0.15 | 0.51 |
| SARS2-L141H | 7.91E-09 | 1.29E-09 | 1.30 | 3.47 |
| SARS2-L141I | 3.67E-09 | 5.05E-10 | 0.19 | 1.06 |
| SARS2-L141K | 4.50E-09 | 1.55E-09 | 0.49 | 1.15 |
| SARS2-L141M | 4.02E-09 | 1.38E-09 | 0.33 | 0.81 |
| SARS2-L141N | 5.05E-09 | 3.77E-10 | 0.65 | 3.99 |

| | | | | |
|---|---|---|---|---|
| SARS2-L141Q | 4.61E-09 | 8.44E-10 | 0.52 | 2.53 |
| SARS2-L141R | 4.12E-09 | 6.10E-10 | 0.36 | 1.49 |
| SARS2-L141S | 5.06E-09 | 9.65E-10 | 0.66 | 2.56 |
| SARS2-L141T | 4.10E-09 | 5.48E-10 | 0.36 | 1.99 |
| SARS2-N142A | 2.11E-09 | 1.00E-09 | -0.60 | 1.83 |
| SARS2-N142L | 2.85E-09 | 1.37E-09 | -0.17 | 0.47 |
| SARS2-N142M | 2.13E-09 | 6.65E-10 | -0.59 | 1.89 |
| SARS2-N142P | 4.57E-09 | 1.18E-09 | 0.51 | 1.44 |
| SARS2-N142S | 2.86E-09 | 9.12E-10 | -0.16 | 0.67 |
| SARS2-G143A | 3.03E-09 | 6.11E-10 | -0.08 | 0.48 |
| SARS2-G143C | Not Active | -- | -- | -- |
| SARS2-G143D | 2.22E-09 | 8.61E-10 | -0.53 | 2.15 |
| SARS2-G143R | 3.19E-09 | 1.22E-09 | -0.01 | 0.31 |
| SARS2-G143S | 3.58E-09 | 6.26E-10 | 0.16 | 0.81 |
| SARS2-G143V | Not Active | -- | -- | -- |
| SARS2-S144A | 1.49E-08 | 2.72E-09 | 2.21 | 6.05 |
| SARS2-S144F | 5.70E-09 | 2.33E-09 | 0.83 | 1.57 |
| SARS2-S144G | 4.97E-08 | 1.63E-08 | 3.95 | 4.28 |
| SARS2-S144M | 1.75E-08 | 3.89E-09 | 2.45 | 4.76 |
| SARS2-S144Y | 1.70E-08 | 1.33E-08 | 2.41 | 1.90 |
| SARS2-N151C | 4.05E-09 | 2.61E-09 | 0.34 | 0.61 |
| SARS2-N151F | 7.04E-09 | 7.63E-09 | 1.13 | 1.09 |
| SARS2-N151G | 3.43E-09 | 1.23E-09 | 0.10 | 0.32 |
| SARS2-N151I | 3.60E-09 | 8.80E-10 | 0.17 | 0.85 |
| SARS2-N151M | 3.47E-09 | 8.06E-10 | 0.11 | 0.58 |
| SARS2-N151R | 4.20E-09 | 1.47E-09 | 0.39 | 1.07 |
| SARS2-N151S | 3.21E-09 | 4.67E-10 | 0.00 | 0.31 |
| SARS2-N151T | 5.23E-09 | 5.49E-09 | 0.71 | 0.85 |
| SARS2-N151W | 3.64E-09 | 1.02E-09 | 0.19 | 0.55 |
| SARS2-N151Y | 3.32E-09 | 7.41E-10 | 0.05 | 0.43 |
| SARS2-V157L | 3.19E-09 | 9.24E-10 | 0.00 | 0.31 |
| SARS2-C160F | 2.55E-09 | 1.98E-10 | -0.33 | 1.46 |
| SARS2-C160N | 2.90E-09 | 5.66E-10 | -0.15 | 0.63 |
| SARS2-C160Y | 2.32E-09 | 3.73E-10 | -0.47 | 1.66 |
| SARS2-M162I | 2.24E-09 | 4.34E-10 | -0.52 | 2.30 |
| SARS2-H164N | 6.53E-09 | 9.60E-10 | 1.03 | 4.61 |
| SARS2-H164Q | 8.27E-09 | 1.45E-09 | 1.37 | 3.94 |
| SARS2-M165I | 7.17E-09 | 2.08E-09 | 1.16 | 2.68 |
| SARS2-M165K | Not Active | -- | -- | -- |

| | | | | |
|---|---|---|---|---|
| SARS2-M165L | 3.47E-09 | 3.80E-10 | 0.12 | 0.34 |
| SARS2-M165R | 2.36E-09 | 1.18E-09 | -0.44 | 0.97 |
| SARS2-M165T | 3.20E-09 | 1.10E-09 | 0.00 | 0.31 |
| SARS2-M165V | 5.54E-09 | 9.23E-10 | 0.79 | 3.85 |
| SARS2-E166A | 1.24E-08 | 3.12E-09 | 1.95 | 5.74 |
| SARS2-E166D | 2.80E-09 | 8.26E-10 | -0.19 | 1.46 |
| SARS2-E166G | 6.11E-09 | 1.94E-09 | 0.93 | 2.60 |
| SARS2-E166K | 1.98E-08 | 5.82E-09 | 2.63 | 3.40 |
| SARS2-E166M | 2.09E-08 | 1.13E-08 | 2.71 | 2.40 |
| SARS2-E166Q | 3.79E-09 | 1.16E-09 | 0.24 | 1.27 |
| SARS2-E166V | 9.97E-09 | 3.24E-09 | 1.64 | 3.19 |
| SARS2-L50F/E166V | 9.81E-09 | 2.50E-09 | 1.61 | 3.63 |
| SARS2-T21I/E166V | 6.67E-09 | 3.25E-09 | 1.06 | 2.98 |
| SARS2-P168Δ | 3.20E-09 | 4.04E-10 | 0.00 | 0.30 |
| SARS2-T169S | 3.13E-09 | 1.01E-09 | -0.04 | 0.32 |
| SARS2-T169P | 3.95E-09 | 8.78E-10 | 0.30 | 1.63 |
| SARS2-G170R | 3.92E-09 | 7.57E-10 | 0.29 | 1.33 |
| SARS2-H172F | Not Active | -- | -- | -- |
| SARS2-H172Q | 5.17E-09 | 1.94E-09 | 0.69 | 1.77 |
| SARS2-A173D | Not Active | -- | -- | -- |
| SARS2-A173G | 3.66E-09 | 4.76E-10 | 0.19 | 0.74 |
| SARS2-A173P | Not Active | -- | -- | -- |
| SARS2-A173S | 2.89E-09 | 5.82E-10 | -0.15 | 0.66 |
| SARS2-A173T | 4.10E-09 | 1.76E-09 | 0.35 | 1.11 |
| SARS2-A173V | 4.50E-09 | 1.19E-09 | 0.49 | 1.63 |
| SARS2-N180D | 3.38E-09 | 1.15E-09 | 0.08 | 0.41 |
| SARS2-P184S | 3.94E-09 | 1.32E-09 | 0.30 | 0.66 |
| SARS2-F185S | 3.46E-09 | 1.92E-09 | 0.11 | 0.41 |
| SARS2-V186F | 3.66E-09 | 1.49E-09 | 0.19 | 0.55 |
| SARS2-R188K | 3.12E-09 | 9.04E-10 | -0.04 | 0.55 |
| SARS2-Q189Δ | Not Active | -- | -- | -- |
| SARS2-Q189E | 4.49E-09 | 1.07E-09 | 0.49 | 1.25 |
| SARS2-Q189F | 3.58E-09 | 7.80E-10 | 0.16 | 0.71 |
| SARS2-Q189G | 4.57E-09 | 2.03E-09 | 0.51 | 1.16 |
| SARS2-Q189H | 5.97E-09 | 1.54E-09 | 0.90 | 2.67 |
| SARS2-Q189K | 3.10E-09 | 1.42E-09 | -0.05 | 0.32 |
| SARS2-Q189L | 3.92E-09 | 7.66E-10 | 0.29 | 2.55 |
| SARS2-Q189P | 5.18E-09 | 1.38E-09 | 0.69 | 2.26 |
| SARS2-Q189R | 3.62E-09 | 1.07E-09 | 0.18 | 0.47 |

| | | | | |
|---|---|---|---|---|
| SARS2-Q189S | 4.18E-09 | 1.20E-09 | 0.38 | 1.68 |
| SARS2-T190I | 3.12E-09 | 5.07E-10 | -0.04 | 0.74 |
| SARS2-A191V | 3.44E-09 | 1.71E-09 | 0.10 | 0.40 |
| SARS2-Q192T | 5.53E-09 | 1.89E-09 | 0.79 | 1.94 |
| SARS2-Q192A | 5.44E-09 | 2.29E-09 | 0.76 | 1.46 |
| SARS2-Q192C | 5.72E-09 | 1.43E-09 | 0.84 | 3.31 |
| SARS2-Q192F | 4.34E-09 | 7.91E-10 | 0.44 | 1.85 |
| SARS2-Q192H | 3.82E-09 | 9.10E-10 | 0.25 | 1.22 |
| SARS2-Q192I | Not Active | -- | -- | -- |
| SARS2-Q192L | 6.49E-09 | 1.72E-09 | 1.02 | 2.61 |
| SARS2-Q192P | 6.14E-09 | 2.13E-09 | 0.94 | 2.73 |
| SARS2-Q192S | 4.25E-09 | 1.18E-09 | 0.41 | 1.11 |
| SARS2-Q192V | 6.50E-09 | 1.89E-09 | 1.02 | 3.94 |
| SARS2-Q192W | 4.35E-09 | 1.05E-09 | 0.44 | 2.10 |
| SARS2-T196M | 2.12E-09 | 2.82E-10 | -0.60 | 2.41 |
| SARS2-V202W | 2.91E-09 | 7.90E-10 | -0.14 | 0.68 |
| SARS2-L205V | 3.06E-09 | 8.78E-10 | -0.07 | 0.42 |
| SARS2-A206V | Not Active | -- | -- | -- |
| SARS2-V212F | 2.56E-09 | 2.82E-10 | -0.33 | 1.28 |
| SARS2-I213V | 3.13E-09 | 1.09E-09 | -0.03 | 0.41 |
| SARS2-L220F | 2.47E-09 | 6.50E-10 | -0.38 | 1.27 |
| SARS2-F223L | 2.74E-09 | 2.33E-10 | -0.22 | 0.79 |
| SARS2-Y239A | 3.15E-09 | 7.33E-10 | -0.03 | 0.30 |
| SARS2-Y239G | 2.16E-09 | 4.75E-10 | -0.57 | 1.70 |
| SARS2-E240M | 2.13E-09 | 1.32E-10 | -0.59 | 3.07 |
| SARS2-P241L | 3.25E-09 | 9.86E-10 | 0.02 | 0.33 |
| SARS2-H246Y | 2.25E-09 | 3.49E-10 | -0.51 | 2.25 |
| SARS2-D248F | 2.94E-09 | 1.47E-09 | -0.12 | 0.43 |
| SARS2-I259L | 2.77E-09 | 8.46E-10 | -0.21 | 0.88 |
| SARS2-A260V | 2.95E-09 | 5.85E-10 | -0.12 | 0.52 |
| SARS2-A266S | 3.67E-09 | 2.09E-09 | 0.20 | 0.43 |
| SARS2-G278R | 2.90E-09 | 1.53E-09 | -0.15 | 0.57 |
| SARS2-R279C | 3.22E-09 | 9.95E-10 | 0.00 | 0.45 |
| SARS2-G283S | 3.58E-09 | 6.88E-10 | 0.16 | 0.77 |
| SARS2-S284G | 2.83E-09 | 4.90E-10 | -0.18 | 0.77 |
| SARS2-A285T | 2.80E-09 | 7.27E-10 | -0.20 | 1.10 |
| SARS2-S301F | 3.51E-09 | 2.31E-09 | 0.13 | 0.50 |

**Table S7. Ensitrelvir inhibition values for the evolution-focused NL63 fusion sublibrary.**

| Mutant ID | IC50 (M) | St. Dev | Log2FC | log10(p-value) |
|---|---|---|---|---|
| SARS2-WT | 1.07E-09 | 3.76E-10 | 0.00 | N/A |
| SARS2-C145A | Not Active | -- | -- | -- |
| SARS2-TQA-WT | Not Active | -- | -- | -- |
| SARS2-TQA-C145A | Not Active | -- | -- | -- |
| SARS2-F003L | 8.01E-10 | 2.99E-10 | -0.41 | 1.18 |
| SARS2-R004K | 6.43E-10 | 1.42E-10 | -0.73 | 2.25 |
| SARS2-F008Q | 1.19E-09 | 2.85E-10 | 0.16 | 0.66 |
| SARS2-K012C | 7.54E-10 | 2.78E-10 | -0.50 | 1.05 |
| SARS2-G015R | 7.51E-10 | 9.64E-11 | -0.51 | 1.64 |
| SARS2-M017V | 5.44E-10 | 2.15E-10 | -0.97 | 3.14 |
| SARS2-Q019R | 1.12E-09 | 3.71E-10 | 0.07 | 0.46 |
| SARS2-T021C | 9.57E-10 | 3.70E-10 | -0.16 | 1.07 |
| SARS2-C022Y | 2.31E-09 | 4.57E-10 | 1.11 | 3.56 |
| SARS2-T024S | 8.30E-10 | 2.20E-10 | -0.36 | 1.18 |
| SARS2-T026V | 7.39E-10 | 2.52E-10 | -0.53 | 1.52 |
| SARS2-L030V | 4.65E-10 | 1.18E-10 | -1.20 | 2.67 |
| SARS2-D033G | 6.70E-10 | 2.09E-10 | -0.67 | 2.26 |
| SARS2-V035T | 9.92E-10 | 4.22E-10 | -0.10 | 0.39 |
| SARS2-Y037T | 1.16E-09 | 4.64E-10 | 0.12 | 0.72 |
| SARS2-C044A | 4.02E-09 | 6.32E-10 | 1.91 | 5.39 |
| SARS2-T045P | 1.02E-09 | 3.14E-10 | -0.07 | 0.31 |
| SARS2-E047T | 1.02E-09 | 1.51E-10 | -0.07 | 0.44 |
| SARS2-D048T | 3.98E-10 | 9.10E-11 | -1.42 | 2.92 |
| SARS2-M049Δ | 1.08E-08 | 3.72E-09 | 3.35 | 4.29 |
| SARS2-L050V | 3.52E-10 | 1.34E-10 | -1.60 | 2.98 |
| SARS2-N051L | 3.46E-10 | 8.46E-11 | -1.62 | 3.00 |
| SARS2-N051Δ | 7.42E-09 | 4.43E-09 | 2.80 | 2.43 |
| SARS2-P052I | 1.72E-08 | 5.01E-09 | 4.01 | 4.23 |
| SARS2-N053D | 8.37E-10 | 2.14E-10 | -0.35 | 1.04 |
| SARS2-E055D | 6.79E-10 | 3.30E-10 | -0.65 | 2.27 |
| SARS2-D056H | 8.59E-10 | 2.40E-10 | -0.31 | 1.04 |
| SARS2-L057A | 1.50E-09 | 4.67E-10 | 0.49 | 1.63 |
| SARS2-L058Y | 3.40E-10 | 1.57E-10 | -1.65 | 3.55 |
| SARS2-I059S | 1.48E-09 | 4.57E-10 | 0.47 | 1.20 |
| SARS2-R060T | 6.88E-10 | 1.62E-10 | -0.63 | 1.55 |

| | | | | |
|---|---|---|---|---|
| SARS2-K061M | 1.18E-09 | 4.51E-10 | 0.15 | 0.45 |
| SARS2-S062R | 1.06E-09 | 4.95E-10 | -0.01 | 0.38 |
| SARS2-N063L | 1.06E-09 | 2.99E-10 | -0.01 | 0.40 |
| SARS2-L067S | 1.06E-09 | 2.55E-10 | -0.01 | 0.32 |
| SARS2-Q069S | 9.08E-10 | 1.26E-10 | -0.23 | 0.60 |
| SARS2-A070H | 1.23E-09 | 5.77E-10 | 0.20 | 1.00 |
| SARS2-G071N | 8.30E-10 | 2.67E-10 | -0.36 | 1.15 |
| SARS2-N072G | 7.13E-10 | 2.11E-10 | -0.58 | 1.83 |
| SARS2-Q074F | 6.47E-10 | 1.23E-10 | -0.72 | 1.49 |
| SARS2-R076G | 7.99E-10 | 1.19E-10 | -0.42 | 1.29 |
| SARS2-I078V | 9.22E-10 | 3.11E-10 | -0.21 | 1.07 |
| SARS2-H080V | 1.53E-09 | 4.40E-10 | 0.52 | 3.70 |
| SARS2-S081T | 9.92E-10 | 5.14E-10 | -0.11 | 0.32 |
| SARS2-Q083H | 7.76E-10 | 3.11E-10 | -0.46 | 1.89 |
| SARS2-N084G | 7.47E-10 | 1.23E-10 | -0.51 | 1.31 |
| SARS2-C085S | 1.39E-09 | 2.01E-10 | 0.38 | 1.31 |
| SARS2-K088R | 7.47E-10 | 1.61E-10 | -0.51 | 1.14 |
| SARS2-L089I | 8.27E-10 | 3.49E-10 | -0.37 | 0.62 |
| SARS2-D092S | 6.78E-10 | 1.16E-10 | -0.66 | 1.53 |
| SARS2-T093Q | 8.93E-10 | 2.55E-10 | -0.26 | 0.93 |
| SARS2-A094S | 8.72E-10 | 3.18E-10 | -0.29 | 1.38 |
| SARS2-P096V | 7.23E-10 | 2.12E-10 | -0.56 | 1.20 |
| SARS2-K097H | 7.74E-10 | 1.33E-10 | -0.46 | 1.02 |
| SARS2-Y101H | 6.24E-10 | 8.12E-11 | -0.78 | 2.26 |
| SARS2-K102V | 4.23E-10 | 1.51E-10 | -1.34 | 3.72 |
| SARS2-V104K | 6.54E-10 | 7.09E-11 | -0.71 | 2.22 |
| SARS2-R105T | 6.73E-10 | 1.50E-10 | -0.66 | 1.57 |
| SARS2-I106L | 7.33E-10 | 3.60E-10 | -0.54 | 2.26 |
| SARS2-Q107K | 9.34E-10 | 2.33E-10 | -0.19 | 0.51 |
| SARS2-Q110D | 1.08E-09 | 2.89E-10 | 0.02 | 0.40 |
| SARS2-T111S | 4.99E-10 | 1.25E-10 | -1.10 | 2.35 |
| SARS2-S113N | Not Active | -- | -- | -- |
| SARS2-V114I | 1.40E-09 | 4.19E-10 | 0.39 | 1.99 |
| SARS2-N119E | 8.82E-10 | 2.60E-10 | -0.28 | 0.89 |
| SARS2-S121I | 1.16E-09 | 5.61E-10 | 0.12 | 0.54 |
| SARS2-P122A | 6.66E-10 | 2.37E-10 | -0.68 | 1.76 |
| SARS2-Y126F | 1.40E-09 | 4.95E-10 | 0.39 | 1.69 |
| SARS2-Q127G | 3.03E-10 | 1.55E-10 | -1.82 | 2.76 |

| | | | | |
|---|---|---|---|---|
| SARS2-C128V | 1.40E-09 | 6.21E-10 | 0.39 | 0.95 |
| SARS2-A129N | 2.62E-09 | 4.21E-09 | 1.30 | 0.75 |
| SARS2-M130L | 1.22E-09 | 5.88E-10 | 0.19 | 1.00 |
| SARS2-P132T | 5.70E-10 | 1.83E-10 | -0.90 | 2.40 |
| SARS2-L141I | 9.37E-10 | 1.90E-10 | -0.19 | 0.50 |
| SARS2-S144A | 2.32E-08 | 2.57E-09 | 4.44 | 7.23 |
| SARS2-V148P | 3.47E-10 | 1.42E-10 | -1.62 | 3.54 |
| SARS2-F150Y | 5.40E-10 | 1.68E-10 | -0.98 | 2.70 |
| SARS2-I152V | 1.37E-09 | 4.53E-10 | 0.36 | 1.70 |
| SARS2-D153R | 5.11E-10 | 1.91E-10 | -1.06 | 2.29 |
| SARS2-Y154N | 1.28E-09 | 5.13E-10 | 0.26 | 0.68 |
| SARS2-ins154D | 1.25E-09 | 1.42E-10 | 0.23 | 0.81 |
| SARS2-D155G | 5.75E-10 | 6.84E-11 | -0.89 | 2.49 |
| SARS2-ins155G | 1.62E-09 | 4.21E-10 | 0.60 | 1.56 |
| SARS2-C156T | 1.50E-09 | 3.45E-10 | 0.49 | 2.26 |
| SARS2-S158E | 1.20E-09 | 3.77E-10 | 0.17 | 1.09 |
| SARS2-M162L | 4.22E-07 | 7.89E-07 | 8.63 | 0.98 |
| SARS2-H164Q | 5.02E-09 | 6.56E-10 | 2.24 | 5.07 |
| SARS2-M165I | 3.09E-09 | 3.99E-10 | 1.53 | 5.88 |
| SARS2-P168G | 2.11E-09 | 3.77E-10 | 0.99 | 4.57 |
| SARS2-T169S | 1.29E-09 | 2.78E-10 | 0.28 | 1.46 |
| SARS2-V171A | 1.29E-09 | 3.94E-10 | 0.27 | 1.08 |
| SARS2-A173V | 1.61E-09 | 3.62E-10 | 0.59 | 1.50 |
| SARS2-T175S | 7.12E-10 | 1.37E-10 | -0.58 | 1.72 |
| SARS2-L177F | 4.92E-10 | 2.62E-10 | -1.12 | 4.36 |
| SARS2-E178T | 4.38E-10 | 2.35E-10 | -1.28 | 2.55 |
| SARS2-N180S | 3.89E-10 | 1.76E-10 | -1.46 | 3.28 |
| SARS2-F181V | Not Active | -- | -- | -- |
| SARS2-P184N | 6.00E-10 | 1.82E-10 | -0.83 | 2.60 |
| SARS2-V186D | 3.16E-09 | 3.35E-10 | 1.57 | 5.80 |
| SARS2-R188Q | 7.62E-10 | 3.36E-10 | -0.49 | 1.22 |
| SARS2-Q189P | 2.58E-09 | 2.96E-10 | 1.27 | 3.52 |
| SARS2-T190S | 7.41E-10 | 1.27E-10 | -0.53 | 1.23 |
| SARS2-A191L | 7.51E-10 | 1.41E-10 | -0.51 | 1.53 |
| SARS2-A193V | 7.75E-10 | 2.59E-10 | -0.46 | 1.90 |
| SARS2-A194E | 7.25E-10 | 2.51E-10 | -0.56 | 1.86 |
| SARS2-G195S | 8.75E-10 | 1.40E-10 | -0.29 | 0.88 |
| SARS2-T196A | 8.92E-10 | 1.28E-10 | -0.26 | 1.06 |

| | | | | |
|---|---|---|---|---|
| SARS2-D197N | 1.00E-09 | 4.37E-10 | -0.09 | 0.35 |
| SARS2-T198L | 1.21E-09 | 5.72E-10 | 0.19 | 0.75 |
| SARS2-T199M | 5.88E-10 | 1.69E-10 | -0.86 | 1.68 |
| SARS2-I200L | 5.19E-10 | 5.04E-11 | -1.04 | 2.45 |
| SARS2-T201S | 6.07E-10 | 1.40E-10 | -0.81 | 2.15 |
| SARS2-V202D | Not Active | -- | -- | -- |
| SARS2-L205V | 5.54E-10 | 1.20E-10 | -0.95 | 2.32 |
| SARS2-W207F | 8.13E-10 | 2.32E-10 | -0.39 | 1.09 |
| SARS2-V212L | 4.42E-10 | 1.84E-10 | -1.27 | 2.80 |
| SARS2-I213L | 9.69E-10 | 2.45E-10 | -0.14 | 0.64 |
| SARS2-D216C | 6.06E-10 | 1.93E-10 | -0.82 | 2.09 |
| SARS2-F219W | 9.57E-10 | 2.95E-10 | -0.16 | 0.73 |
| SARS2-N221C | 9.21E-10 | 1.75E-10 | -0.21 | 0.67 |
| SARS2-R222S | 8.83E-10 | 1.99E-10 | -0.27 | 0.70 |
| SARS2-F223T | 1.18E-09 | 4.63E-10 | 0.14 | 0.51 |
| SARS2-T224R | 6.55E-10 | 1.62E-10 | -0.70 | 2.12 |
| SARS2-T225V | Not Active | -- | -- | -- |
| SARS2-T226N | 8.11E-10 | 1.92E-10 | -0.40 | 0.95 |
| SARS2-L227V | 6.30E-10 | 2.17E-10 | -0.76 | 2.75 |
| SARS2-N228D | Not Active | -- | -- | -- |
| SARS2-D229G | 3.50E-10 | 1.76E-10 | -1.61 | 2.78 |
| SARS2-L232E | 1.34E-09 | 4.99E-10 | 0.32 | 0.94 |
| SARS2-V233W | 7.39E-10 | 1.56E-10 | -0.53 | 1.38 |
| SARS2-K236A | 9.20E-10 | 4.02E-10 | -0.21 | 0.68 |
| SARS2-Y237N | 7.16E-10 | 3.14E-10 | -0.58 | 2.45 |
| SARS2-N238G | 1.27E-09 | 4.58E-10 | 0.25 | 0.87 |
| SARS2-E240T | Not Active | -- | -- | -- |
| SARS2-P241S | 5.81E-10 | 1.47E-10 | -0.88 | 2.36 |
| SARS2-L242V | 6.63E-10 | 2.48E-10 | -0.69 | 3.24 |
| SARS2-T243S | 6.87E-10 | 1.72E-10 | -0.64 | 2.02 |
| SARS2-Q244S | 5.90E-10 | 9.63E-11 | -0.85 | 2.13 |
| SARS2-D245V | 7.00E-10 | 1.67E-10 | -0.61 | 1.58 |
| SARS2-D245Δ | Not Active | -- | -- | -- |
| SARS2-H246E | 7.52E-10 | 2.23E-10 | -0.50 | 1.15 |
| SARS2-H246Δ | 4.29E-10 | 8.66E-11 | -1.32 | 3.03 |

| | | | | |
|---|---|---|---|---|
| SARS2-V247Δ | Not Active | -- | -- | -- |
| SARS2-D248Δ | Not Active | -- | -- | -- |
| SARS2-D248E | 9.52E-10 | 3.12E-10 | -0.16 | 0.45 |
| SARS2-I249C | 9.13E-10 | 3.80E-10 | -0.23 | 0.96 |
| SARS2-L250Y | Not Active | -- | -- | -- |
| SARS2-G251S | 9.47E-10 | 4.32E-10 | -0.17 | 1.00 |
| SARS2-P252I | 5.53E-10 | 9.94E-11 | -0.95 | 2.13 |
| SARS2-S254A | 1.21E-09 | 4.83E-10 | 0.18 | 1.16 |
| SARS2-Q256K | 9.18E-10 | 3.94E-10 | -0.22 | 0.97 |
| SARS2-I259V | 7.27E-10 | 1.40E-10 | -0.55 | 1.62 |
| SARS2-A260S | 1.15E-09 | 4.98E-10 | 0.10 | 0.71 |
| SARS2-L262E | 8.81E-10 | 2.36E-10 | -0.28 | 1.15 |
| SARS2-D263Q | Not Active | -- | -- | -- |
| SARS2-M264L | 4.91E-10 | 6.29E-11 | -1.12 | 2.22 |
| SARS2-C265L | 8.03E-10 | 2.52E-10 | -0.41 | 1.15 |
| SARS2-L268I | 1.27E-05 | 2.54E-05 | 13.54 | 1.00 |
| SARS2-K269Q | 1.03E-09 | 2.31E-10 | -0.05 | 0.42 |
| SARS2-E270Δ | Not Active | -- | -- | -- |
| SARS2-E270H | 8.33E-10 | 2.27E-10 | -0.36 | 1.57 |
| SARS2-L271H | 1.20E-09 | 7.06E-10 | 0.18 | 0.74 |
| SARS2-L272H | Not Active | -- | -- | -- |
| SARS2-Q273H | Not Active | -- | -- | -- |
| SARS2-Q273E | 9.87E-10 | 3.93E-10 | -0.11 | 0.66 |
| SARS2-N274E | 8.09E-10 | 1.37E-10 | -0.40 | 1.73 |
| SARS2-N274Δ | 1.98E-10 | 1.41E-10 | -2.43 | 4.11 |
| SARS2-M276F | 7.79E-09 | 1.99E-08 | 2.87 | 0.73 |
| SARS2-N277G | 8.82E-10 | 3.17E-10 | -0.27 | 1.61 |
| SARS2-R279K | 8.12E-10 | 1.95E-10 | -0.39 | 1.16 |
| SARS2-T280N | 4.64E-10 | 6.13E-11 | -1.20 | 2.80 |
| SARS2-S284Y | 3.44E-10 | 1.55E-10 | -1.63 | 3.35 |
| SARS2-A285S | 8.52E-10 | 3.60E-10 | -0.32 | 0.88 |
| SARS2-L286S | 1.00E-09 | 2.90E-10 | -0.09 | 0.43 |
| SARS2-E288C | 2.14E-09 | 7.48E-10 | 1.01 | 2.40 |
| SARS2-P293L | 3.12E-10 | 7.67E-11 | -1.77 | 2.79 |
| SARS2-F294A | 1.35E-09 | 4.26E-10 | 0.34 | 1.88 |

| | | | | |
|---|---|---|---|---|
| SARS2-D295E | 6.72E-10 | 2.60E-10 | -0.67 | 1.56 |
| SARS2-R298K | 7.96E-10 | 2.89E-10 | -0.42 | 2.05 |
| SARS2-C300M | 9.88E-10 | 2.16E-10 | -0.11 | 0.57 |
| SARS2-S301Y | 1.04E-09 | 2.62E-10 | -0.04 | 0.34 |
| SARS2-T304N | 8.42E-10 | 2.15E-10 | -0.34 | 1.06 |
| SARS2-F305L | Not Active | -- | -- | -- |
| SARS2-DH245-246ΔΔ | Not Active | -- | -- | -- |
| SARS2-VD247-248ΔΔ | Not Active | -- | -- | -- |
| SARS2-NL63-Loop1 | Not Active | -- | -- | -- |
| SARS2-NL63-Loop22 | 2.83E-09 | 9.70E-10 | 1.41 | 2.61 |
| SARS2-NL63-Loop39 | 9.90E-07 | 1.90E-06 | 9.86 | 0.96 |
| SARS2-NL63-Loop70 | 8.34E-10 | 4.27E-10 | -0.36 | 1.15 |
| SARS2-NL63-Loop83 | 1.44E-09 | 3.58E-10 | 0.43 | 2.16 |
| SARS2-NL63-Loop91 | 1.09E-09 | 3.19E-10 | 0.04 | 0.35 |
| SARS2-NL63-Loop129 | Not Active | -- | -- | -- |
| SARS2-NL63-Loop166 | 2.51E-09 | 3.52E-10 | 1.24 | 4.58 |
| SARS2-NL63-Loop175 | Not Active | -- | -- | -- |
| SARS2-NL63-Loop275 | 1.40E-09 | 4.21E-10 | 0.39 | 1.48 |
| SARS2-NL63-Loop300 | 1.12E-09 | 2.87E-10 | 0.07 | 0.47 |
| SARS2-NL63-Loop39/129 | Not Active | -- | -- | -- |
| SARS2-NL63-Loop39/166 | 1.49E-07 | 2.90E-07 | 7.13 | 1.07 |
| SARS2-NL63-Loop39/175 | Not Active | -- | -- | -- |
| SARS2-NL63-Loop39/129/166 | Not Active | -- | -- | -- |
| SARS2-NL63-Loop39/129/175 | Not Active | -- | -- | -- |
| SARS2-NL63-Loop39/166/175 | Not Active | -- | -- | -- |
| SARS2-NL63-Loop129/166 | Not Active | -- | -- | -- |
| SARS2-NL63-Loop129/175 | Not Active | -- | -- | -- |
| SARS2-NL63-Loop129/166/175 | Not Active | -- | -- | -- |
| SARS2-NL63-Loop166/175 | Not Active | -- | -- | -- |
| SARS2-NL63-Loop39/129/166/175 | Not Active | -- | -- | -- |

**Table S8. AVI-4516 IC50s**

| Mutant ID | IC50 (M) | St. Dev | Log2FC | -log10(q-value) |
|---|---|---|---|---|
| SARS2-F003L | 2.11E-08 | 9.82E-10 | 0.1 | 0.64 |
| SARS2-R004K | 1.70E-08 | 1.37E-09 | -0.21 | 1.19 |
| SARS2-R004S | 1.34E-08 | 5.34E-10 | -0.43 | 2.22 |
| SARS2-K005F | 8.69E-09 | 8.00E-10 | -1.06 | 3.63 |
| SARS2-K005W | 8.15E-09 | 1.64E-09 | -1.16 | 3.87 |
| SARS2-A007T | 1.16E-08 | 1.15E-09 | -0.65 | 3.08 |
| SARS2-K012C | 1.70E-08 | 1.36E-09 | -0.21 | 1.28 |
| SARS2-G015R | 1.61E-08 | 9.46E-10 | -0.29 | 1.76 |
| SARS2-G015S | 1.15E-08 | 1.76E-09 | -0.66 | 3.12 |
| SARS2-G015V | 1.61E-08 | 3.40E-09 | -0.17 | 0.44 |
| SARS2-M017V | 1.44E-08 | 2.10E-09 | -0.45 | 2.39 |
| SARS2-Q019R | 1.26E-08 | 1.50E-09 | -0.65 | 3.39 |
| SARS2-T021C | 1.51E-08 | 9.11E-10 | -0.38 | 2.28 |
| SARS2-T021I | 1.01E-08 | 1.02E-09 | -0.85 | 3.47 |
| SARS2-T21I/E166V | 1.18E-08 | 7.88E-10 | -0.62 | 2.93 |
| SARS2-C022Y | 1.63E-08 | 4.04E-09 | -0.28 | 0.86 |
| SARS2-T024S | 1.63E-08 | 1.03E-09 | -0.28 | 1.74 |
| SARS2-T025I | 1.56E-08 | 6.31E-10 | -0.22 | 1.13 |
| SARS2-T026V | 1.77E-08 | 1.54E-09 | -0.15 | 0.86 |
| SARS2-L030I | 1.11E-08 | 2.21E-09 | -0.71 | 3.08 |

| | | | | |
|---|---|---|---|---|
| SARS2-L030V | 1.69E-08 | 7.71E-10 | -0.22 | 1.47 |
| SARS2-D033F | 1.17E-08 | 1.17E-09 | -0.63 | 3.04 |
| SARS2-D033G | 1.71E-08 | 1.23E-09 | -0.2 | 1.30 |
| SARS2-D033V | 1.05E-08 | 1.09E-09 | -0.79 | 3.38 |
| SARS2-D034M | 9.71E-09 | 9.64E-10 | -0.9 | 3.53 |
| SARS2-V035T | 9.95E-09 | 4.84E-09 | -0.21 | 0.27 |
| SARS2-Y037T | 1.46E-08 | 1.64E-09 | -0.44 | 2.40 |
| SARS2-H041L | 9.44E-09 | 2.03E-09 | -0.95 | 3.65 |
| SARS2-H041Y | 1.05E-08 | 1.60E-09 | -0.8 | 3.44 |
| SARS2-C044A | 1.12E-08 | 9.19E-10 | -0.81 | 3.80 |
| SARS2-S046F | 1.37E-08 | 3.56E-09 | 0.25 | 0.50 |
| SARS2-D048N | 1.02E-08 | 5.04E-10 | -0.84 | 3.34 |
| SARS2-D048T | 1.68E-08 | 1.16E-09 | -0.23 | 1.49 |
| SARS2-M049I | 3.03E-09 | 5.26E-10 | -2.58 | 4.36 |
| SARS2-M049K | 1.80E-08 | 6.08E-09 | -0.02 | 0.03 |
| SARS2-M049L | 5.77E-10 | 1.88E-10 | -4.98 | 4.52 |
| SARS2-M049T | 2.55E-08 | 2.31E-09 | 0.49 | 3.27 |
| SARS2-M049Δ | 4.96E-08 | 9.57E-09 | 1.33 | 2.56 |
| SARS2-L050F | 1.64E-08 | 1.23E-09 | -0.15 | 0.68 |
| SARS2-L050V | 1.66E-08 | 2.53E-09 | -0.24 | 1.23 |
| SARS2-L50F/E166V | 2.42E-08 | 8.07E-09 | 0.42 | 0.78 |
| SARS2-N051L | 1.55E-08 | 1.90E-09 | -0.35 | 1.85 |

| | | | | |
|---|---|---|---|---|
| SARS2-N051Δ | 2.59E-08 | 7.92E-09 | 0.39 | 0.83 |
| SARS2-P052I | 1.53E-08 | 5.91E-09 | -0.36 | 0.85 |
| SARS2-N053D | 1.79E-08 | 1.09E-09 | -0.14 | 0.81 |
| SARS2-E055D | 1.66E-08 | 8.19E-10 | -0.24 | 1.58 |
| SARS2-D056H | 1.67E-08 | 2.26E-09 | -0.23 | 1.26 |
| SARS2-L057A | 1.37E-08 | 4.76E-09 | -0.52 | 1.19 |
| SARS2-L058Y | 1.54E-08 | 2.12E-09 | -0.36 | 1.70 |
| SARS2-I059S | 2.43E-08 | 1.83E-09 | 0.3 | 2.17 |
| SARS2-R060T | 1.79E-08 | 1.23E-09 | -0.14 | 0.82 |
| SARS2-K061M | 2.25E-08 | 8.38E-10 | 0.19 | 1.43 |
| SARS2-S062R | 1.89E-08 | 1.35E-09 | -0.06 | 0.28 |
| SARS2-NL63-Loop129/166 | 1.23E-08 | 1.49E-09 | -0.68 | 3.52 |
| SARS2-NL63-Loop166 | 3.43E-08 | 1.57E-09 | 0.8 | 4.92 |
| SARS2-NL63-Loop166/175 | 1.54E-08 | 7.67E-10 | -0.36 | 2.17 |
| SARS2-NL63-Loop175 | 1.54E-08 | 1.38E-09 | -0.35 | 2.02 |
| SARS2-NL63-Loop22 | 1.34E-08 | 8.70E-10 | -0.44 | 2.27 |
| SARS2-NL63-Loop275 | 1.59E-08 | 6.22E-10 | -0.3 | 1.85 |
| SARS2-NL63-Loop300 | 1.50E-08 | 7.69E-09 | -0.4 | 0.54 |
| SARS2-NL63-Loop39/129/166 | 1.34E-08 | 2.25E-09 | -0.56 | 2.56 |
| SARS2-NL63-Loop39/129/175 | 1.58E-08 | 1.32E-09 | -0.32 | 1.75 |
| SARS2-NL63-Loop39/166 | 1.78E-08 | 1.05E-09 | -0.15 | 0.86 |

| | | | | |
|---|---|---|---|---|
| SARS2-NL63-Loop39/166/175 | 1.56E-08 | 2.08E-09 | -0.33 | 1.74 |
| SARS2-NL63-Loop39/175 | 1.61E-08 | 9.03E-10 | -0.29 | 1.75 |
| SARS2-NL63-Loop70 | 8.34E-09 | 4.44E-09 | -0.47 | 0.55 |
| SARS2-NL63-Loop83 | 1.61E-08 | 2.53E-09 | -0.29 | 1.37 |
| SARS2-NL63-Loop91 | 1.73E-08 | 1.31E-09 | -0.18 | 1.14 |
| SARS2-L067S | 1.41E-08 | 1.28E-09 | -0.48 | 2.65 |
| SARS2-Q069S | 1.61E-08 | 1.33E-09 | -0.29 | 1.74 |
| SARS2-A070H | 1.85E-08 | 7.98E-10 | -0.09 | 0.54 |
| SARS2-G071N | 1.70E-08 | 4.35E-10 | -0.22 | 1.44 |
| SARS2-N072G | 1.68E-08 | 1.32E-09 | -0.23 | 1.46 |
| SARS2-Q074F | 1.81E-08 | 1.78E-09 | -0.12 | 0.60 |
| SARS2-R076G | 7.91E-09 | 4.56E-09 | -0.54 | 0.95 |
| SARS2-I078V | 1.83E-08 | 1.75E-09 | -0.11 | 0.52 |
| SARS2-H080V | 2.28E-08 | 1.35E-09 | 0.21 | 1.54 |
| SARS2-S081T | 1.34E-08 | 1.98E-09 | -0.56 | 2.70 |
| SARS2-Q083H | 1.73E-08 | 1.34E-09 | -0.18 | 1.14 |
| SARS2-N084G | 1.50E-08 | 1.24E-09 | -0.39 | 2.28 |
| SARS2-C085S | 1.90E-08 | 6.94E-10 | -0.05 | 0.25 |
| SARS2-K088R | 1.31E-08 | 1.25E-09 | -0.47 | 2.43 |
| SARS2-L089F | 8.00E-09 | 1.28E-09 | -1.18 | 3.83 |
| SARS2-L089I | 1.63E-08 | 6.56E-10 | -0.27 | 1.74 |

| | | | | |
|---|---|---|---|---|
| SARS2-L089P | 9.47E-09 | 6.27E-10 | -0.94 | 3.53 |
| SARS2-K090M | 1.37E-08 | 6.24E-10 | -0.41 | 2.10 |
| SARS2-K090Q | 1.40E-08 | 2.01E-09 | -0.38 | 1.91 |
| SARS2-K090R | 1.93E-08 | 2.19E-09 | 0.08 | 0.29 |
| SARS2-K090T | 1.60E-08 | 2.69E-09 | -0.19 | 0.68 |
| SARS2-D092G | 1.48E-08 | 2.16E-09 | -0.3 | 1.30 |
| SARS2-T093I | 1.79E-08 | 1.41E-09 | -0.02 | 0.06 |
| SARS2-T093Q | 1.87E-08 | 2.07E-09 | -0.08 | 0.28 |
| SARS2-A094S | 1.69E-08 | 6.01E-10 | -0.22 | 1.46 |
| SARS2-P096L | 1.32E-08 | 1.65E-09 | -0.47 | 2.33 |
| SARS2-P096V | 1.69E-08 | 8.93E-10 | -0.22 | 1.44 |
| SARS2-K097H | 1.66E-08 | 9.12E-10 | -0.25 | 1.58 |
| SARS2-K102V | 1.61E-08 | 1.37E-09 | -0.29 | 1.70 |
| SARS2-V104K | 1.84E-08 | 3.86E-10 | -0.1 | 0.58 |
| SARS2-I106L | 1.51E-08 | 1.37E-09 | -0.39 | 2.28 |
| SARS2-Q107K | 1.94E-08 | 6.50E-10 | -0.02 | 0.10 |
| SARS2-P108S | 9.16E-09 | 2.28E-09 | -0.99 | 3.65 |
| SARS2-Q110D | 1.97E-08 | 8.86E-10 | 0 | 0.00 |
| SARS2-T111S | 1.33E-08 | 1.13E-09 | -0.56 | 3.00 |
| SARS2-S113T | 1.08E-08 | 9.54E-10 | -0.74 | 3.27 |
| SARS2-V114I | 1.95E-08 | 2.32E-09 | -0.01 | 0.04 |
| SARS2-A116E | 1.03E-08 | 1.10E-09 | -0.81 | 3.44 |

| | | | | |
|---|---|---|---|---|
| SARS2-A116L | 4.53E-08 | 3.26E-09 | 1.32 | 7.74 |
| SARS2-A116V | 5.02E-08 | 7.74E-09 | 1.46 | 3.08 |
| SARS2-N119E | 1.64E-08 | 8.33E-10 | -0.27 | 1.69 |
| SARS2-S121I | 1.25E-08 | 2.66E-09 | 0.12 | 0.21 |
| SARS2-P122A | 1.98E-08 | 1.71E-09 | 0.01 | 0.03 |
| SARS2-S123F | 2.64E-08 | 8.30E-09 | 0.54 | 1.40 |
| SARS2-Y126F | 2.57E-08 | 2.69E-09 | 0.5 | 3.18 |
| SARS2-Y126M | 1.31E-08 | 3.89E-09 | -0.47 | 1.23 |
| SARS2-Q127G | 1.27E-08 | 5.16E-09 | -0.63 | 1.05 |
| SARS2-Q127H | 1.87E-08 | 5.27E-09 | 0.04 | 0.07 |
| SARS2-C128V | 2.75E-08 | 2.94E-09 | 0.48 | 2.28 |
| SARS2-A129N | 8.29E-09 | 4.96E-09 | -1.25 | 1.87 |
| SARS2-A129V | 1.69E-08 | 2.28E-09 | -0.1 | 0.36 |
| SARS2-M130L | 1.73E-08 | 6.33E-10 | -0.19 | 1.27 |
| SARS2-P132A | 1.04E-08 | 1.11E-09 | -0.8 | 3.43 |
| SARS2-P132H | 1.10E-08 | 1.06E-09 | -0.73 | 3.27 |
| SARS2-P132L | 1.10E-08 | 9.69E-10 | -0.73 | 3.24 |
| SARS2-P132R | 1.12E-08 | 1.69E-09 | -0.7 | 3.27 |
| SARS2-P132S | 1.13E-08 | 5.51E-10 | -0.69 | 3.08 |
| SARS2-P132T | 1.35E-08 | 1.12E-09 | -0.54 | 2.95 |
| SARS2-L141E | 1.01E-08 | 5.74E-09 | -0.84 | 1.60 |
| SARS2-L141H | 1.90E-08 | 2.98E-09 | 0.07 | 0.20 |

| | | | | |
|---|---|---|---|---|
| SARS2-L141I | 1.60E-08 | 9.35E-10 | -0.3 | 1.85 |
| SARS2-L141M | 2.57E-08 | 6.25E-10 | 0.5 | 3.27 |
| SARS2-L141Q | 1.64E-08 | 1.65E-09 | -0.15 | 0.61 |
| SARS2-L141R | 1.38E-08 | 1.75E-09 | -0.4 | 1.91 |
| SARS2-L141S | 2.17E-08 | 9.49E-09 | 0.25 | 0.37 |
| SARS2-L141T | 1.95E-08 | 9.72E-09 | 0.1 | 0.13 |
| SARS2-N142A | 1.71E-08 | 2.37E-09 | -0.09 | 0.27 |
| SARS2-N142L | 4.90E-09 | 4.55E-10 | -1.89 | 3.97 |
| SARS2-N142M | 5.11E-09 | 1.17E-09 | -1.83 | 4.36 |
| SARS2-N142S | 1.42E-08 | 2.87E-09 | -0.35 | 1.34 |
| SARS2-G143A | 1.37E-08 | 1.78E-09 | -0.41 | 1.93 |
| SARS2-G143C | 9.45E-09 | 1.28E-09 | -0.94 | 3.63 |
| SARS2-G143D | 8.34E-09 | 1.00E-09 | -1.12 | 3.77 |
| SARS2-G143S | 1.21E-08 | 2.00E-09 | -0.59 | 2.78 |
| SARS2-G143V | 1.26E-08 | 2.16E-09 | -0.53 | 2.6 |
| SARS2-S144A | 3.95E-08 | 3.28E-09 | 1.12 | 5.91 |
| SARS2-S144F | 9.11E-09 | 7.31E-10 | -1 | 3.58 |
| SARS2-S144M | 1.79E-08 | 4.87E-09 | -0.02 | 0.03 |
| SARS2-S144Y | 9.92E-09 | 1.97E-09 | -0.87 | 3.58 |
| SARS2-C145A | 1.53E-08 | 8.63E-10 | -0.36 | 2.19 |
| SARS2-V148P | 1.37E-08 | 1.11E-09 | -0.52 | 2.86 |
| SARS2-F150Y | 1.51E-08 | 7.83E-10 | -0.38 | 2.28 |

| | | | | |
|---|---|---|---|---|
| SARS2-N151C | 1.51E-08 | 3.77E-09 | 0.39 | 0.96 |
| SARS2-N151F | 3.33E-08 | 2.52E-09 | 0.87 | 4.77 |
| SARS2-N151G | 1.29E-08 | 8.63E-10 | -0.49 | 2.48 |
| SARS2-N151R | 2.59E-08 | 6.72E-09 | 0.51 | 1.29 |
| SARS2-N151T | 1.92E-08 | 2.90E-09 | 0.74 | 3.23 |
| SARS2-N151W | 1.66E-08 | 7.05E-09 | 0.53 | 1.15 |
| SARS2-N151Y | 1.99E-08 | 2.68E-09 | 0.79 | 3.55 |
| SARS2-I152V | 2.27E-08 | 9.35E-10 | 0.2 | 1.53 |
| SARS2-D153R | 1.66E-08 | 1.82E-09 | -0.24 | 1.39 |
| SARS2-Y154N | 2.53E-08 | 2.09E-09 | 0.36 | 2.51 |
| SARS2-ins154D | 2.42E-08 | 1.71E-09 | 0.3 | 2.17 |
| SARS2-ins155G | 3.30E-08 | 4.27E-09 | 0.74 | 3.80 |
| SARS2-D155G | 1.59E-08 | 2.50E-09 | -0.3 | 1.46 |
| SARS2-C156T | 2.01E-08 | 3.67E-09 | 0.03 | 0.08 |
| SARS2-S158E | 1.99E-08 | 3.57E-09 | 0.02 | 0.04 |
| SARS2-C160F | 1.18E-08 | 2.24E-09 | -0.63 | 3.01 |
| SARS2-C160N | 9.18E-09 | 9.16E-10 | -0.99 | 3.63 |
| SARS2-C160Y | 1.01E-08 | 3.29E-09 | -0.85 | 2.81 |
| SARS2-M162I | 1.01E-08 | 1.95E-09 | -0.85 | 3.37 |
| SARS2-M162L | 1.28E-08 | 3.43E-09 | -0.62 | 2.28 |
| SARS2-H164N | 1.93E-08 | 3.78E-09 | 0.09 | 0.18 |
| SARS2-H164Q | 2.36E-08 | 1.26E-09 | 0.26 | 1.87 |

| | | | | |
|---|---|---|---|---|
| SARS2-M165I | 7.70E-08 | 3.19E-09 | 1.97 | 9.17 |
| SARS2-M165K | 8.44E-09 | 1.41E-09 | -1.11 | 3.81 |
| SARS2-M165R | 8.96E-09 | 1.09E-09 | -1.02 | 3.65 |
| SARS2-E166Q | 1.30E-08 | 1.28E-09 | -0.48 | 2.47 |
| SARS2-E166V | 1.29E-08 | 2.40E-09 | -0.49 | 2.29 |
| SARS2-P168G | 2.95E-08 | 1.65E-09 | 0.58 | 4.48 |
| SARS2-P168Δ | 8.11E-09 | 1.33E-09 | -1.16 | 3.83 |
| SARS2-T169P | 1.19E-08 | 1.61E-09 | -0.61 | 3.01 |
| SARS2-T169S | 1.45E-08 | 3.03E-09 | -0.32 | 1.16 |
| SARS2-G170R | 9.01E-09 | 8.65E-10 | -1.01 | 3.62 |
| SARS2-V171A | 2.28E-08 | 9.62E-10 | 0.21 | 1.52 |
| SARS2-H172F | 1.33E-08 | 6.81E-09 | -0.45 | 0.72 |
| SARS2-A173G | 1.14E-08 | 2.82E-09 | -0.68 | 2.86 |
| SARS2-A173P | 9.55E-09 | 1.53E-09 | -0.93 | 3.47 |
| SARS2-A173S | 9.45E-09 | 1.42E-09 | -0.94 | 3.63 |
| SARS2-A173T | 1.95E-08 | 8.34E-09 | 0.1 | 0.11 |
| SARS2-A173V | 4.06E-08 | 3.48E-09 | 1.16 | 5.03 |
| SARS2-T175S | 1.33E-08 | 2.29E-09 | -0.57 | 2.56 |
| SARS2-L177F | 1.48E-08 | 2.08E-09 | -0.42 | 2.12 |
| SARS2-E178T | 1.32E-08 | 1.78E-09 | -0.58 | 3.02 |
| SARS2-N180D | 1.06E-08 | 1.91E-09 | -0.78 | 3.24 |
| SARS2-N180S | 1.30E-08 | 1.39E-09 | -0.6 | 3.02 |

| | | | | |
|---|---|---|---|---|
| SARS2-F181V | 1.03E-08 | 3.45E-09 | -0.93 | 1.81 |
| SARS2-P184N | 1.65E-08 | 1.32E-09 | -0.26 | 1.58 |
| SARS2-P184S | 1.07E-08 | 5.01E-10 | -0.76 | 3.21 |
| SARS2-F185S | 8.60E-09 | 5.02E-10 | -1.08 | 3.61 |
| SARS2-V186D | 2.86E-08 | 1.84E-09 | 0.54 | 3.54 |
| SARS2-V186F | 1.37E-08 | 1.69E-09 | -0.41 | 2.05 |
| SARS2-R188K | 1.24E-08 | 3.57E-09 | 0.11 | 0.19 |
| SARS2-R188Q | 2.25E-08 | 1.42E-09 | 0.19 | 1.39 |
| SARS2-Q189E | 3.15E-08 | 4.33E-09 | 0.79 | 2.87 |
| SARS2-Q189F | 2.00E-08 | 1.95E-09 | 0.14 | 0.58 |
| SARS2-Q189G | 1.44E-08 | 4.50E-09 | -0.34 | 0.73 |
| SARS2-Q189H | 2.86E-08 | 3.42E-09 | 0.66 | 2.96 |
| SARS2-Q189K | 7.52E-09 | 8.11E-10 | -1.27 | 3.81 |
| SARS2-Q189L | 1.52E-08 | 3.63E-09 | -0.26 | 0.92 |
| SARS2-Q189P | 2.84E-08 | 1.67E-09 | 0.53 | 3.97 |
| SARS2-Q189R | 7.94E-09 | 1.42E-09 | -1.19 | 3.87 |
| SARS2-Q189S | 2.07E-08 | 2.14E-09 | 0.19 | 0.88 |
| SARS2-Q189Δ | 1.04E-08 | 1.44E-09 | -0.8 | 3.43 |
| SARS2-T190I | 2.04E-08 | 2.90E-09 | 0.17 | 0.64 |
| SARS2-T190S | 1.82E-08 | 1.64E-09 | -0.12 | 0.59 |
| SARS2-A191L | 1.60E-08 | 1.43E-09 | -0.3 | 1.75 |
| SARS2-A191V | 1.33E-08 | 1.67E-09 | -0.45 | 2.27 |

| | | | | |
|---|---|---|---|---|
| SARS2-Q192C | 9.12E-09 | 9.96E-10 | -0.99 | 3.63 |
| SARS2-Q192F | 9.86E-09 | 1.15E-09 | -0.88 | 3.53 |
| SARS2-Q192H | 1.48E-08 | 8.32E-09 | -0.3 | 0.45 |
| SARS2-Q192I | 1.36E-08 | 4.65E-09 | -0.42 | 0.91 |
| SARS2-Q192P | 1.29E-08 | 5.38E-09 | -0.5 | 1.06 |
| SARS2-Q192T | 1.15E-08 | 2.16E-09 | -0.66 | 2.96 |
| SARS2-Q192V | 1.10E-08 | 2.23E-09 | -0.72 | 3.10 |
| SARS2-Q192W | 8.59E-09 | 1.50E-09 | -1.08 | 3.81 |
| SARS2-A193V | 1.80E-08 | 2.00E-09 | -0.13 | 0.65 |
| SARS2-A194E | 1.42E-08 | 1.56E-09 | -0.47 | 2.65 |
| SARS2-G195S | 1.78E-08 | 1.90E-09 | -0.15 | 0.79 |
| SARS2-T196A | 1.96E-08 | 1.29E-09 | -0.01 | 0.04 |
| SARS2-T196M | 1.72E-08 | 2.56E-09 | -0.08 | 0.23 |
| SARS2-D197N | 1.26E-08 | 2.89E-09 | -0.65 | 2.65 |
| SARS2-T198L | 2.01E-08 | 1.36E-09 | 0.03 | 0.12 |
| SARS2-T199M | 1.55E-08 | 2.87E-09 | -0.35 | 1.60 |
| SARS2-I200L | 1.65E-08 | 1.19E-09 | -0.25 | 1.58 |
| SARS2-T201S | 1.61E-08 | 1.84E-09 | -0.29 | 1.66 |
| SARS2-V202D | 1.46E-08 | 1.75E-09 | -0.43 | 2.31 |
| SARS2-V202W | 8.92E-09 | 2.58E-09 | -1.03 | 3.65 |
| SARS2-L205V | 1.12E-08 | 4.78E-10 | -0.7 | 3.08 |
| SARS2-A206V | 1.02E-08 | 1.25E-09 | -0.84 | 3.48 |

| | | | | |
|---|---|---|---|---|
| SARS2-W207F | 1.93E-08 | 2.46E-09 | -0.03 | 0.08 |
| SARS2-V212F | 9.13E-09 | 3.28E-09 | -0.33 | 0.36 |
| SARS2-V212L | 1.37E-08 | 9.16E-10 | -0.53 | 2.86 |
| SARS2-I213L | 2.06E-08 | 8.71E-10 | 0.07 | 0.37 |
| SARS2-I213V | 1.22E-08 | 4.66E-10 | -0.58 | 2.77 |
| SARS2-D216C | 1.51E-08 | 1.84E-09 | -0.38 | 1.91 |
| SARS2-F219W | 1.56E-08 | 9.01E-10 | -0.34 | 2.06 |
| SARS2-N221C | 1.67E-08 | 6.65E-10 | -0.23 | 1.50 |
| SARS2-R222S | 1.71E-08 | 1.79E-09 | -0.2 | 1.14 |
| SARS2-F223L | 1.44E-08 | 1.12E-09 | -0.33 | 1.70 |
| SARS2-F223T | 1.99E-08 | 1.57E-09 | 0.01 | 0.05 |
| SARS2-T224R | 1.64E-08 | 1.13E-09 | -0.26 | 1.66 |
| SARS2-T225V | 1.58E-08 | 2.32E-09 | -0.32 | 1.01 |
| SARS2-T226N | 1.82E-08 | 1.95E-09 | -0.12 | 0.54 |
| SARS2-L227V | 1.42E-08 | 2.95E-09 | -0.47 | 1.84 |
| SARS2-N228D | 2.05E-08 | 1.32E-09 | 0.06 | 0.28 |
| SARS2-D229G | 1.39E-08 | 1.88E-09 | -0.5 | 2.56 |
| SARS2-L232E | 1.87E-08 | 1.12E-09 | -0.08 | 0.41 |
| SARS2-V233W | 1.65E-08 | 1.61E-09 | -0.25 | 1.44 |
| SARS2-Y237N | 1.75E-08 | 1.68E-09 | -0.17 | 0.92 |
| SARS2-N238G | 2.06E-08 | 1.15E-09 | 0.07 | 0.33 |
| SARS2-Y239A | 1.09E-08 | 2.25E-09 | -0.74 | 3.27 |

| | | | | |
|---|---|---|---|---|
| SARS2-Y239G | 8.69E-09 | 1.53E-09 | -1.06 | 3.79 |
| SARS2-E240M | 1.47E-08 | 2.37E-09 | -0.31 | 1.38 |
| SARS2-E240T | 1.72E-08 | 7.74E-10 | -0.2 | 1.30 |
| SARS2-T243S | 1.55E-08 | 2.15E-09 | -0.35 | 1.75 |
| SARS2-Q244S | 1.77E-08 | 2.11E-09 | -0.15 | 0.72 |
| SARS2-D245V | 1.62E-08 | 1.04E-09 | -0.28 | 1.75 |
| SARS2-D245Δ | 1.66E-08 | 1.14E-09 | -0.25 | 1.52 |
| SARS2-H246E | 1.72E-08 | 5.93E-10 | -0.2 | 1.33 |
| SARS2-H246Y | 1.02E-08 | 1.13E-09 | -0.83 | 3.47 |
| SARS2-H246Δ | 1.60E-08 | 1.48E-09 | -0.3 | 1.75 |
| SARS2-V247Δ | 1.66E-08 | 1.17E-09 | -0.24 | 1.53 |
| SARS2-D248E | 1.71E-08 | 4.27E-10 | -0.2 | 1.37 |
| SARS2-D248F | 1.10E-08 | 1.25E-09 | -0.73 | 3.27 |
| SARS2-D248Δ | 1.67E-08 | 9.95E-10 | -0.24 | 1.41 |
| SARS2-I249C | 1.72E-08 | 2.61E-09 | -0.19 | 0.91 |
| SARS2-L250Y | 1.52E-08 | 2.20E-09 | -0.37 | 1.51 |
| SARS2-G251S | 1.26E-08 | 6.32E-09 | 0.13 | 0.18 |
| SARS2-P252I | 1.62E-08 | 2.00E-09 | -0.28 | 1.51 |
| SARS2-S254A | 1.43E-08 | 2.90E-09 | 0.31 | 0.78 |
| SARS2-Q256K | 1.64E-08 | 2.83E-09 | -0.26 | 1.14 |
| SARS2-I259L | 1.03E-08 | 1.07E-09 | -0.82 | 3.44 |
| SARS2-I259V | 1.10E-08 | 2.52E-09 | -0.06 | 0.11 |

| | | | | |
|---|---|---|---|---|
| SARS2-A260S | 1.92E-08 | 2.58E-09 | -0.03 | 0.11 |
| SARS2-A260V | 9.59E-09 | 5.52E-10 | -0.92 | 3.47 |
| SARS2-L262E | 1.69E-08 | 8.83E-10 | -0.22 | 1.46 |
| SARS2-D263Q | 1.30E-08 | 9.96E-10 | -0.6 | 3.15 |
| SARS2-M264L | 1.87E-08 | 3.41E-09 | -0.08 | 0.22 |
| SARS2-C265L | 1.68E-08 | 2.43E-09 | -0.23 | 1.25 |
| SARS2-A266S | 1.36E-08 | 2.70E-09 | -0.42 | 1.91 |
| SARS2-K269Q | 2.08E-08 | 1.14E-09 | 0.08 | 0.45 |
| SARS2-E270H | 1.72E-08 | 8.51E-10 | -0.19 | 1.28 |
| SARS2-L271H | 1.70E-08 | 3.66E-09 | -0.21 | 0.62 |
| SARS2-L272H | 1.58E-08 | 1.65E-09 | -0.32 | 1.79 |
| SARS2-Q273E | 1.87E-08 | 2.55E-09 | -0.07 | 0.28 |
| SARS2-Q273H | 1.60E-08 | 6.27E-10 | -0.3 | 1.84 |
| SARS2-N274E | 1.63E-08 | 7.64E-10 | -0.28 | 1.75 |
| SARS2-N274Δ | 1.20E-08 | 2.60E-09 | -0.71 | 2.45 |
| SARS2-M276F | 1.57E-08 | 1.74E-09 | -0.33 | 1.74 |
| SARS2-N277G | 9.33E-09 | 3.10E-09 | -0.3 | 0.55 |
| SARS2-R279K | 1.88E-08 | 1.76E-09 | -0.07 | 0.29 |
| SARS2-T280N | 1.63E-08 | 1.29E-09 | -0.27 | 1.68 |
| SARS2-G283S | 1.11E-08 | 4.40E-10 | -0.71 | 3.09 |
| SARS2-S284G | 9.83E-09 | 4.17E-10 | -0.89 | 3.43 |
| SARS2-S284Y | 1.61E-08 | 1.08E-09 | -0.29 | 1.76 |

| | | | | |
|---|---|---|---|---|
| SARS2-A285T | 1.60E-08 | 2.67E-09 | -0.18 | 0.68 |
| SARS2-L286S | 1.94E-08 | 2.66E-09 | 0.1 | 0.31 |
| SARS2-E288C | 1.71E-08 | 1.12E-09 | -0.09 | 0.37 |
| SARS2-F294A | 2.31E-08 | 3.87E-09 | 0.35 | 1.19 |
| SARS2-D295E | 1.13E-08 | 1.15E-09 | -0.68 | 3.17 |
| SARS2-R298K | 1.67E-08 | 5.10E-10 | -0.12 | 0.56 |
| SARS2-C300M | 1.29E-08 | 1.11E-09 | -0.49 | 2.48 |
| SARS2-S301F | 1.98E-08 | 5.72E-09 | 0.12 | 0.24 |
| SARS2-S301Y | 1.88E-08 | 4.82E-09 | 0.05 | 0.10 |
| SARS2-F305L | 1.50E-08 | 2.50E-09 | -0.28 | 1.18 |
| SARS2-TQA | 9.00E-09 | 2.91E-09 | -1.13 | 3.83 |

**Table S9. Sequence of 11 SARS-CoV-2 peptide substrates.**

| **NSP Junction** | **Substrate Sequence** |
|---|---|
| NSP 4-5 | K(5-FAM)TSAVLQSGFRKM(dnp) |
| NSP 5-6 | K(5-FAM)SGVTFQSAVKRT(dnp) |
| NSP 6-7 | K(5-FAM)KVATVQSKMSDV(dnp) |
| NSP 7-8 | K(5-FAM)NRATLQAIASEF(dnp) |
| NSP 8-9 | K(5-FAM)SAVKLQNNELSP(dnp) |
| NSP 9-10 | K(5-FAM)ATVRLQAGNATE(dnp) |
| NSP 10-11 | K(5-FAM)REPMLQSADAQS(dnp) |
| NSP 12-13 | K(5-FAM)PHTVLQAVGACV(dnp) |
| NSP 13-14 | K(5-FAM)NVATLQAENVTG(dnp) |
| NSP 14-15 | K(5-FAM)TFTRLQSLENVA(dnp) |
| NSP 15-16 | K(5-FAM)TYPKLQSSQAWQ(dnp) |

**Table S10. Sequence of Matriptase, CMV protease, and the catalytic domain of MMP12, and peptide substrates, each with a 5-Carboxyfluorescein (5-FAM) fluorescent dye and a Dinitrophenolate (dnp) quencher.**

| **Matriptase Amino Acid Sequence** |
|---|
| VVGGTDADEGEWPWQVSLHALGQGHICGASLISPNWLVSAAHCYIDDRGFRYSDPTQWTAFLGLHDQSQRSAPGVQERRLKRIISHPFFNDFTFDYDIALLELEKPAEYSSMVRPISLPDASHVFPAGKAIWVTGWGHTQYGGTGALILQKGEIRVINQTTCENLLPQQITPRMMCVGFLSGGVDSCQGDSGGPLSSVEADGRIFQAGVVSWGDGCAQRNKPGVYTRLPLFRDWIKENTGV |
| **Matriptase Substrate** |
| dRK(5-FAM)RQARVVGGK(DNP) |
| **CMV Amino Acid Sequence** |
| MTMDEQQSQAVAPVYVGGFLARYDQSPDEAELLLPRDVVEHWLHAQGQGQPSLSVALPLNINHDDTAVVGHVAAMQSVRDGLFCLGCVTSPRFLEIVRRASEKSELVSRGPVSPLQPDKVVEFLSGSYAGLSLSSRRCDDVEVATSLSGSETTPFKHVALCSVGRRRGTLAVYGRDPEWVTQRFPDLTAADRDGLRAQWQRCGSTAVDVSGDPFRSDSYGLLGNSVDALYIRERLPKLRYDKQLVGVTERESYVKA |
| **CMV Substrate** |
| dRK(5-FAM)-Tbg-Tbg-NASSRLK(DNP), Tbg=tert-butylglycine |
| **cdMMP12 Amino Acid Sequence** |
| MGPVWRKHYITYRINNYTPDMNREDVDYAIRKAFQVWSNVTPLKFSKINTGMADILVVFARGAHGDDHAFDGKGGILAHAFGPGSGIGGDAHFDEDEFWTTHSGGTNLFLTAVHEIGHSLGLGHSSDPKAVMFPTYKYVDINTFRLSADDIRGIQSLY |
| **cdMMP12 Substrate** |
| K(5-FAM)RPKPVE-Nval-WRK(DNP), Nval= norvaline |
